# Roughening instability of growing 3D bacterial colonies

**DOI:** 10.1101/2022.05.09.491177

**Authors:** Alejandro Martínez-Calvo, Tapomoy Bhattacharjee, R. Kōnane Bay, Hao Nghi Luu, Anna M. Hancock, Ned S. Wingreen, Sujit S. Datta

**Author notes:** These authors contributed equally to this work (Alejandro Martínez-Calvo, Tapomoy Bhattacharjee).

## Abstract

How do growing bacterial colonies get their shapes? While colony morphogenesis is well-studied in 2D, many bacteria grow as large colonies in 3D environments, such as gels and tissues in the body, or soils, sediments, and subsurface porous media. Here, we describe a morphological instability exhibited by large colonies of bacteria growing in 3D. Using experiments in transparent 3D granular hydrogel matrices, we show that dense colonies of four different species of bacteria—*Escherichia coli, Vibrio cholerae, Pseudomonas aeruginosa*, and *Komagataeibacter sucrofermentans*—generically roughen as they consume nutrients and grow beyond a critical size, eventually adopting a characteristic branched, broccoli-like, self-affine morphology independent of variations in the cell type and environmental conditions. This behavior reflects a key difference between 2D and 3D colonies: while a 2D colony may access the nutrients needed for growth from the third dimension, a 3D colony inevitably becomes nutrient-limited in its interior, driving a transition to rough growth at its surface. We elucidate the onset of roughening using linear stability analysis and numerical simulations of a continuum model that treats the colony as an ‘active fluid’ whose dynamics are driven by nutrient-dependent cellular growth. We find that when all dimensions of the growing colony substantially exceed the nutrient penetration length, nutrient-limited growth drives a 3D morphological instability that recapitulates essential features of the experimental observations. Our work thus provides a framework to predict and control the organization of growing colonies—as well as other forms of growing active matter, such as tumors and engineered living materials—in 3D environments.

---

Bacteria are known to thrive in diverse ecosystems and habitats [1–5]. In nature, bacteria can be found growing on surfaces, which, in addition to the ease of visualization in 2D, have led laboratory studies to typically focus on colony growth on 2D planar surfaces. However, in many cases in nature, bacteria grow in 3D habitats such as gels and tissues inside of hosts [6–8], soils and other subsurface media [9, 10], wastewater treatment devices, and naturally-occurring bodies of water [11–13]. Nonetheless, despite their prevalence, the morphodynamics of bacterial colonies growing in such 3D environments remains largely unknown. Here, we ask: what determines the shape of a bacterial colony growing in 3D? And are there general characteristics and universal principles that span across species and specific environmental conditions?

Studies of bacteria growing on 2D planar surfaces have revealed a large variety of growth patterns, ranging from circular-shaped colonies [14–21] to herringbone patterns [22] and ramified, rough interfaces [14–20,23]. Some of these patterns become 3D as the colony can grow and deform into the third dimension. These morphologies are now understood to arise from friction between the growing colony and the surface, and differential access to nutrients, which may also be available from the third dimension [14–20, 22, 24–28]. The emergent patterns have been rationalized by incorporating these key ingredients into reaction-diffusion equations [2, 15–18, 20, 24, 29–36], active continuum theories [21–23, 30, 32, 37–50], and agent-based models [30, 32, 38, 40, 45, 46, 51–54]. Moreover, it has been suggested that these morphologies and growth patterns may in turn influence the global function and physiology of bacterial colonies [55–57], including resistance to antibiotics [58–60] and parasites [61], resilience to environmental changes [62, 63], and genetic diversity [53, 55, 56, 64–70]. In stark contrast to the considerable scientific effort devoted to studying 2D growth, little is known about the collective processes, the resulting morphologies, or their functional consequences for bacteria growing in 3D.

From a physics standpoint, growing in 3D is fundamentally different from growing in 2D, both in terms of nutrient access and ability to grow and deform into an additional dimension. Consequently, we expect colony morphodynamics—the way a colony’s overall shape changes over time—to also be different. Some recent studies hint that this is indeed the case, showing how specific mechanical interactions imposed by a 3D environment can influence the morphology of growing biofilms. For instance, external fluid flows are now known to trigger the formation of streamers that stem from an initially surface-attached colony [11, 71]. Under quiescent conditions, small (≲ 10s of cells across) biofilm colonies constrained in strongly-cross-linked bulk gels adopt internally-ordered structures as they grow and push outward [72], mediated by elastic stresses arising at the interface between the colony and its stiff environment. However, the behavior of larger bacterial colonies growing freely in quiescent 3D environments remains underexplored, despite the fact that they represent a fundamental building block of more complex natural colonies.

Here, we combine experiments, theoretical modeling, and numerical simulations to unravel the morphodynamics of large colonies growing in 3D environments. By performing experiments with four different species of bacteria growing in transparent and easily-deformed granular hydrogel matrices, we find that dense colonies growing in 3D undergo a morphological instability when they become nutrient-limited in their interior, i.e. when their size sufficiently exceeds the nutrient penetration length into the colony bulk. Independent of variations in cell type and environmental conditions, growing colonies eventually adopt a generic highly-branched, broccoli-like, self-affine morphology with a characteristic roughness exponent and power spectrum. Employing a continuum ‘active fluid’ model that incorporates the coupling between nutrient diffusion, consumption, and cell growth, we trace the origin of this instability to an interplay of competition for nutrients with growth-pressure-driven colony expansion. In particular, we find that these dense 3D colonies inevitably become nutrient-limited in their interior, which eventually drives the transition to a rough, branched periphery of the colony. Our results thus help establish a framework to predict and control the organization of growing colonies in 3D habitats. These principles could also extend to other morphogenesis processes driven by growth in 3D environments, such as developmental processes [73, 74], tumor growth [75–79], and the expansion of engineered soft living materials [80, 81].

## RESULTS

### Growing 3D bacterial colonies undergo a common morphological instability

To explore the morphodynamics of dense bacterial colonies growing in 3D environments, we use a bioprinter to inject densely-packed (number density *ρ* ~ 10^12^ cells mL^-1^) colonies of bacteria inside granular hydrogel matrices, as sketched in Fig. 1*A* (see *Materials and Methods* and *SI Appendix*). To start, we focus on experiments using *E. coli* as a model system. The colonies have long cylindrical shapes with radii *R* that vary between ~ 20 – 250 μm, at least tens of cells across. The matrices have four notable characteristics [34, 85–87]: (i) they are transparent, enabling us to directly visualize colony morphodynamics in situ; (ii) they are easily mechanically deformed and rearranged (yielded), and thus, do not strongly constrain colony growth but simply keep the cells suspended in 3D unlike in previous work [72]; (iii) they can be designed to be replete with oxygen and nutrients, given that the internal mesh size of the individual hydrogel grains is ~ 40 – 100 nm [88] and thus permissive of free diffusion of small molecules throughout each matrix, thereby sustaining cellular proliferation over many generations; and (iv) the sizes of the interstitial pores between adjacent hydrogel grains can be precisely tuned by changing the hydrogel grain packing density [34, 88]. To isolate the influence of cellular growth on colony morphodynamics, we use hydrogel matrices with mean pore sizes between ≈ 0.1 – 1.0 μm, i.e., smaller or comparable to the diameter of a single cell. Hence, the bacteria are stuck inside the pores; even if they are nominally motile, they cannot self-propel through the pore space. Nevertheless, as the cells consume nutrients, they grow—transiently deforming and yielding the surrounding matrix—and push their progeny out into the neighboring available pores (Movies S1–S2).

**FIG. 1.**
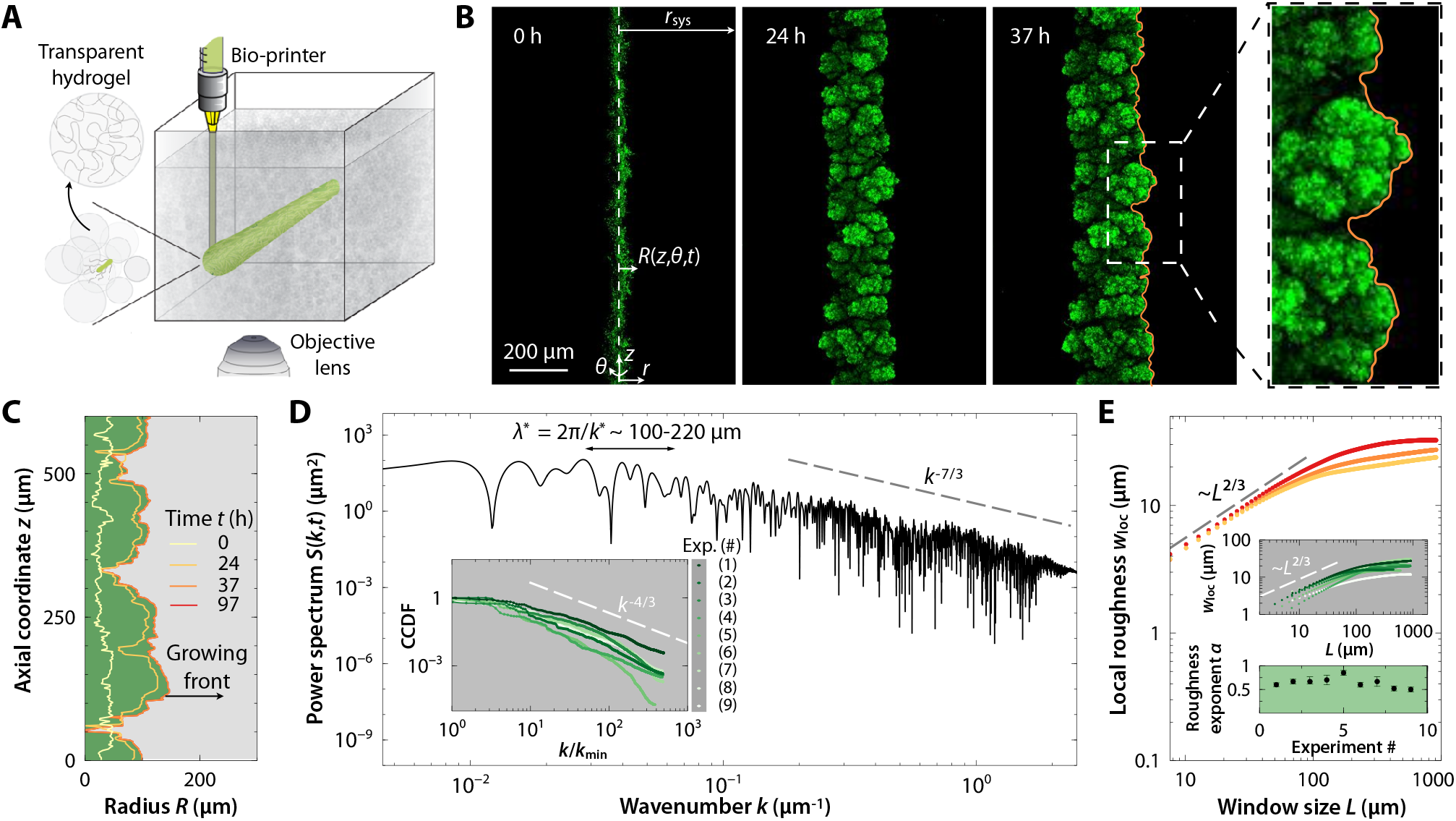
Dense bacterial colonies roughen as they grow in 3D. (*A*) Schematic of the experiments in which a dense bacterial colony (green cylinder) is 3D-printed within a transparent matrix made of jammed hydrogel grains (gray). The matrix locally yields and fluidizes as cells are injected into the pore space, and then rapidly re-jams around the dense-packed cells, holding them in place. The individual hydrogel grains are highly swollen in liquid media containing salts and nutrients, which can freely pass through the hydrogels. However, the interstitial pores between hydrogel grains are smaller than the cell body size, thereby suppressing any motility and holding the cells in place; the colony then expands outward solely through cellular growth and division into adjacent available pores. We use confocal microscopy to obtain 3D stacks of optical slices of cell body fluorescence at different depths in the medium. (*B*) Snapshots of the time evolution of a colony of *E. coli* displaying a roughening instability as the colony grows. The rightmost panel displays a magnified of the colony shown in *B* at 37 hours. The images show maximum-intensity projections of the confocal optical slices taken at different depths in the medium. (*C*) Profiles of the leading edge of the colony shown in *B* at the same times as in *B*. (*D*) Power spectrum *S*(*k,t*) of the leading edge of the colony in *B* at time *t* = 37 h as a function of the axial (along *z*) wavenumber *k*. The lower and upper limits of the wavenumbers displayed correspond to the size of the domain and the experimental resolution limit, respectively. (*Inset*) Complementary cumulative distribution function (CCDF) of the power spectra of different experiments for *E. coli* colonies, testing different strains, hydrogel matrix stiffnesses and mean pore sizes, and nutrient concentrations, as summarized in Table S1; *k*_min_ is the wavenumber corresponding to the domain size that we use to normalize the the different CCDFs. We observe similar power-law scaling, with a plateau at small *k*, in all cases. (*E*) Local roughness *w*_loc_ as a function of the observation window size *L* for the experiment shown in *B* at times *t* = 24, 37, and 97 h. (*Insets*) Local roughness *w*_loc_ as a function of *L* (top), and corresponding roughness exponent *α* (bottom), for the same experiments shown in the *Inset* to *D* at the same times.

It was recently shown that colonies of motile *E. coli* in similar matrices but with larger pores—sufficiently large for cells to swim through—smooth out any perturbations to their colony morphology using directed motility, retaining flat, cylindrical surfaces [35, 36]. We observe starkly-differing behavior for the case of purely growth-driven colony expansion considered here. When the matrix pore size is small enough to abolish the influence of motility, a growing colony of *E. coli* instead exhibits a striking morphological instability— adopting a rough, highly-branched, broccoli-like interface, as shown by the example in Fig. 1*B* and Movie S3. The cells constitutively express green fluorescent protein (GFP) in their cytoplasms, which enables us to directly visualize the colony morphodynamics in 3D across scales spanning from the width of a single cell to that of the entire colony using confocal microscopy; Fig. 1*B* shows bottom-up projections of the cellular fluorescence intensity measured using a 3D stack of confocal micrographs, with successive planar slices taken at different depths within the colony. This growth-induced roughening is markedly different from that observed in Ref. [72], where much smaller bacterial colonies growing as inclusions in stiff, cross-linked bulk gels retained smooth surfaces. Moreover, while highly-branched shapes have been previously observed for colonies growing on planar 2D surfaces, they are thought to only arise when the underlying surface is depleted in nutrients, thereby generating stochasticity in the ability of cells to access nutrient [14–20, 24, 25, 37]; in our case, however, the surrounding matrix is nutrient-replete.

To characterize the colony shape, we track its onedimensional (1D) leading edge over time, as shown in Fig. 1 *C*. The rough colony surface exhibits fluctuations in the axial direction *z* over a broad range of length scales, ranging from the size of a single cell to hundreds of cells. However, these fluctuations appear to have a characteristic maximal size ~ 100 – 200 μm, as shown by the ‘florets’—to use the analogy of broccoli—in Figs. 1 *B* and *C*. These features can be quantified using the power spectral density of spatial fluctuations, 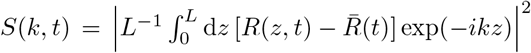, as shown in Fig. 1*D* at time *t* = 37 h; here, 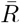 is the average radius of the leading edge, *k* = 2*π*/λ is the axial wavenumber corresponding to a wavelength λ, and *L* is the length of the analyzed region of the colony along the axial direction *z*. In particular, *S*(*k*) eventually becomes time-independent (Fig. S1) and exhibits a power-law decay ~ *k*^-*ν*^ with *v* ≈ 7/3 at sufficiently small length scales (large *k*), as indicated by the dashed line in Fig. 1*D*, which implies that the spatial fluctuations of the rough colony surface have a fractal structure over multiple scales. However, *S*(*k*) becomes bounded at sufficiently large length scales (small *k*), as indicated in panel *D*. Indeed, we again find that the majority of the power is confined to wavenumbers corresponding to characteristic wavelengths λ* ~ 100 – 200 μm—as highlighted by the complementary cumulative distribution function (CCDF) of the power spectrum shown in the inset to Fig. 1*D*.

We further characterize the colony surface roughening by computing the local roughness of the leading edge, quantified by the local variance of the colony radius, 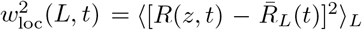, where 〈·〉 denotes the spatial average over different windows of size *L* along the axial direction *z*, and 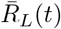 is the average radius in this window. We find that the local roughness of the leading edge scales as *w*_loc_ ~ *L^α^* up to a window size comparable to the characteristic wavelength λ* ~ 100 – 200 μm, above which *w*_loc_ saturates (Fig. 1*E*). The so-called roughness exponent *α* ≈ 2/3, as shown by the dashed line, corresponding to a fractal dimension *d*_f_ = 2 – *α* ~ 1.2 – 1.4 [89]. Indeed, the power-law decay of the power spectrum *S*(*k*) and this roughness exponent agree well with the relation expected from the celebrated Family-Vicsek dynamic scaling of fractal interfaces in one dimension, namely that *v* = 1 + 2*α* [89, 90]—confirming that over length scales smaller than the characteristic ‘floret’ size λ* ~ 100 – 200 μm, the colony surface is a fractal.

To investigate the generality of this morphological instability, we perform replicates of this same experiment, exploring a broad range of different cell types and environmental conditions (Table S1, Figs. S1–S14). Remarkably, we find similar colony roughening that arises within just a few hours—i.e., comparable to the doubling time of the cells—in all cases. Growing colonies of cells that are either motile or non-motile in unconfined liquid, but are too confined to be motile in the hydrogel matrices, show similar morphodynamics (Figs. S1–S2, S4–S6, S9)—confirming that roughening is driven solely by cellular growth and division. For this case of growth-driven colony expansion, we also observe similar overall features of roughening independent of the hydrogel matrix pore size and deformability (Figs. S1–S6, S9–S14)—suggesting that this morphological instability is not strongly sensitive to specific granular features of, or mechanical interactions with, the surrounding matrix. Finally, we also find similar roughening for colonies in matrices with differing nutrient characteristics and concentrations, with differing initial radii, and importantly, across different strains of either *E. coli* or the biofilm-formers *V. cholerae, P. aeruginosa*, and *K. sucrofermentans* (Figs. S1–S14, S23)—indicating that this morphological instability is not a manifestation of a specific nutrient environment or cell type. Taken together, these results establish that large, dense bacterial colonies growing freely in 3D generically roughen, adopting the same characteristic broccoli-like rough morphology (see insets to Figs. 1*D* and *E*) that is fractal at small length scales, and bounded at larger length scales by a characteristic ‘floret’ size spanning hundreds of cells across.

### The interiors of large, dense colonies are depleted of a substrate essential for growth, causing surface roughening

What causes colony roughening? Close inspection of the confocal micrographs for the experiment in Fig. 1 provides a clue: we find that cells only ≲ 20 μm from the colony surface are fluorescent, while those deeper inside the colony lose fluorescence, as shown in Fig. S15. Because the GFP fluorescence acts as a proxy indicator for live and metabolically-active cells, and because cellular metabolism of carbon-containing nutrients such as sugars is both nutrient- and oxygen-dependent, this observation suggests that colony growth is limited to this small surface layer.

This suggestion is corroborated by a simple balance of either nutrient or oxygen 1D diffusion into the cylindrical colony with consumption by the cells (see *SI Appendix*), which suggests that they penetrate into the colony over a length scale 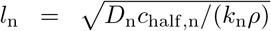 or 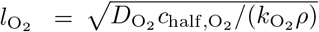, where *ρ* is cell density, *D*_n_, *D*_O_2__ are the diffusivities, *k*_n_, *k*_O_2__ are the maximal uptake rates per cell, and *c*_half,n_, *c*_half,O_2__ are characteristic Michaelis-Menten concentrations of nutrient or oxygen. In particular, using parameter values representative of our experiments (Table I) yields *l_n_* ≈ 1 – 9 μm and *l*_O_2__ ≈ 5 μm, comparable to the length scale over which fluorescence persists. (Because both *l*_n_ and *l*_O_2__ are comparable to each other for all our experiments, hereafter, we will use the subscript “s” and the term *substrate* for whichever limits cellular growth and division; indeed, the theoretical results that follow are not noticeably altered when considering nutrient versus oxygen as the limiting substrate, as detailed in the *SI Appendix* and shown e.g., in Fig. S16.) Thus, for a large and dense 3D bacterial colony, growth is localized to just a few layers of cells at the colony surface, with either nutrient or oxygen acting as a limiting substrate. Consequently, we expect that the relative strength of random fluctuations in cellular growth and division along the colony surface is amplified, potentially driving a roughening instability in a manner similar to 2D colonies [30, 32, 33, 52, 91–93].

**Table I.**
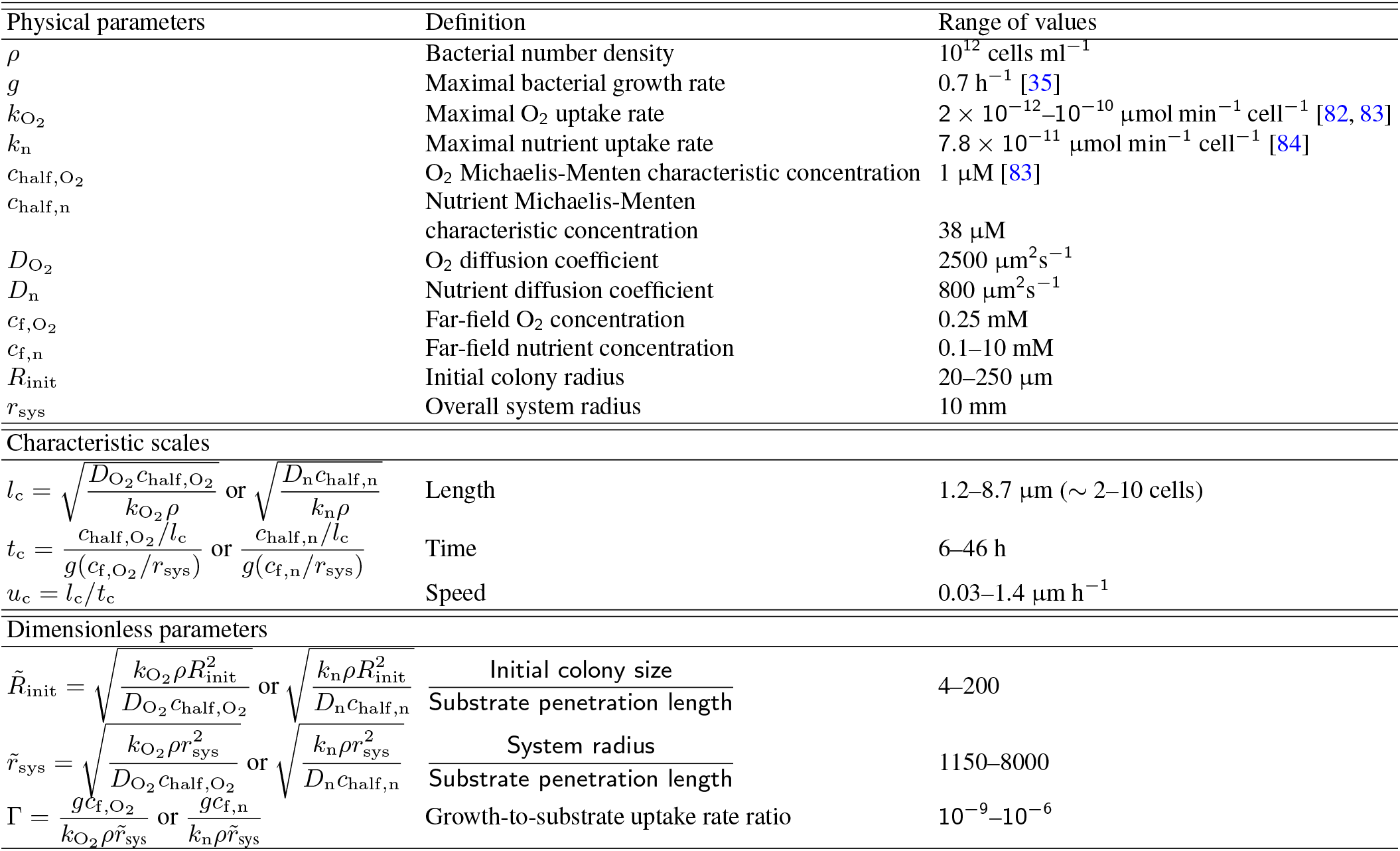
Characteristic parameters for experiments with *E. coli*.

As a final test of this hypothesis, we repeat our experiments, but now with each colony initially inoculated on the planar surface atop a granular hydrogel matrix, exposed to humid air above—just as in conventional studies of growth on 2D gels. In this case, the colony can either grow laterally in 2D on the surface, accessing nutrients and oxygen from the third dimension akin to purely-2D colonies; or it can grow *into* the 3D matrix below, becoming substrate-limited in its interior akin to purely-3D colonies. Thus, when the hydrogel matrix is nutrient-replete, we expect that all cells in the colony on the surface of the matrix remain metabolically-active and fluorescent, and the lateral colony edge remains smooth, as expected for purely-2D colonies that are not nutrient-depleted [14–21]; by contrast, the cells that grow into the underlying matrix should eventually lose fluorescence, and the corresponding colony surface will roughen, similar to purely-3D colonies. These expectations are indeed borne out by the experiments, as shown in Figs. S17–S18. Thus, unlike 2D colonies that can use growth substrates that are available from the third dimension, dense colonies growing in 3D environments inevitably become substrate-limited in their interiors—causing growth to be localized at their surfaces, amplifying inherent random fluctuations in growth along the surface, and thereby generating a roughening instability.

### Continuum model of a growing 3D bacterial colony

To rationalize the experimental observations and test our hypothesis regarding the surface-roughening instability, we model a densely-packed growing bacterial colony embedded in a 3D hydrogel by means of a minimal continuum theory, where bacteria are treated as an ‘active fluid’ whose expansion is driven by substrate-dependent growth. A similar approach was established previously to model growing bacterial colonies in other settings [30, 32, 33, 37, 39, 50, 70], as well as other forms of biological morphodynamics [94–100]. The model is schematized for a slab geometry, which can represent a small region of the surface of a cylindrical colony, in Fig. 2*A*. In our approach, the colony is described in terms of a substrate concentration field, *c*(***x**, t*), a velocity field, ***u***(***x**,t*), and a growth pressure field inside the colony, *p*(***x**,t*), where ***x*** and *t* denote positional coordinate and time, respectively. Because the bacteria inside the colony are effectively incompressible and close-packed, we assume a constant cellular density *ρ* ~ 10^12^ cells mL^-1^, and assume that the velocity field in the interior of the growing colony obeys Darcy’s law in which the local velocity field is proportional to the local pressure gradient ∇*p* inside the colony, which yields a curl-free velocity field.

**FIG. 2.**
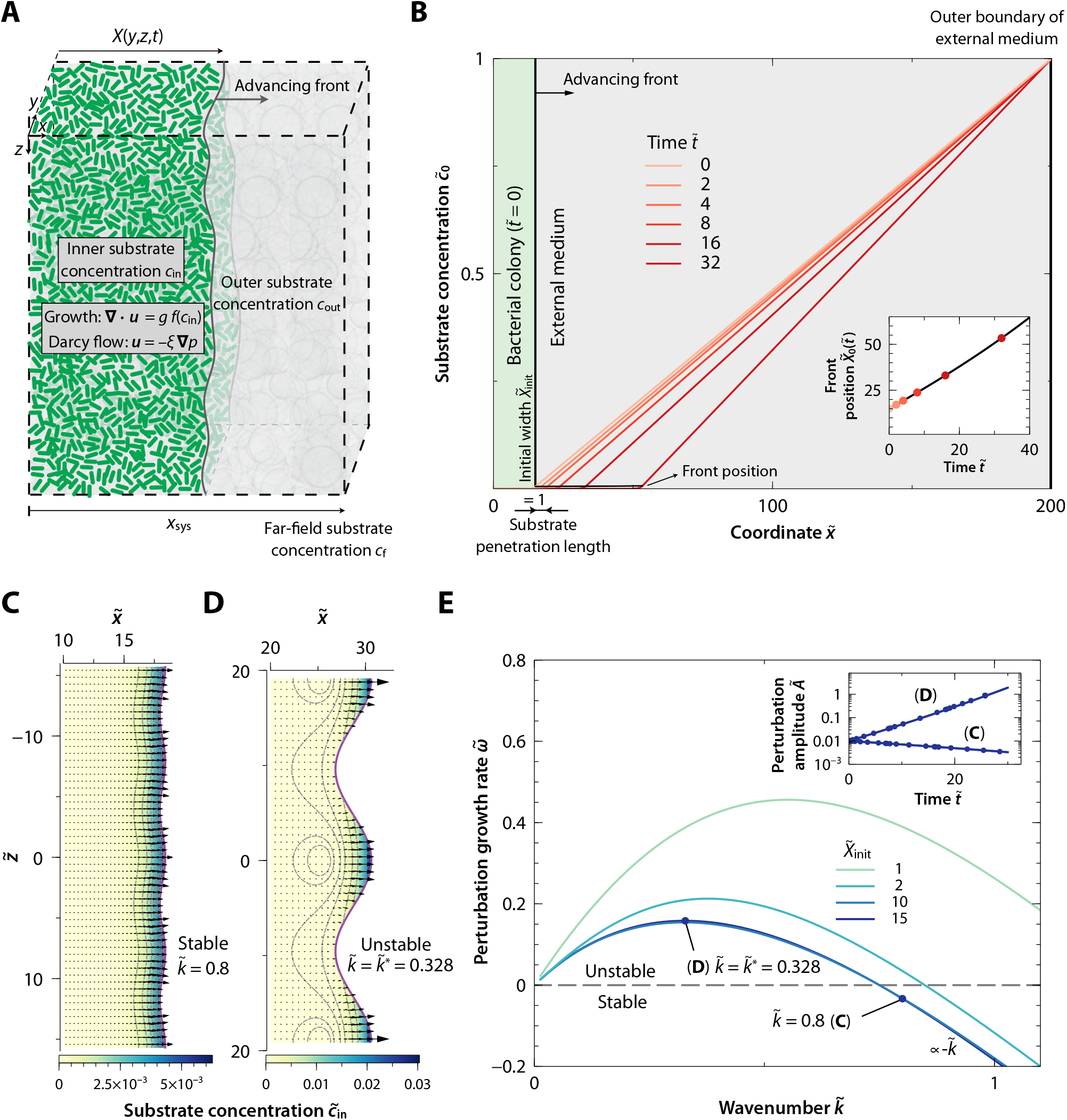
A minimal continuum model suggests that colonies growing in 3D are intrinsically unstable. (*A*) Schematic of the model, in which we treat a slab-shaped bacterial colony (green) as an ‘active fluid’ that expands along the *x* coordinate due to the pressure generated by growth, mediated by substrate availability and consumption. The model is 2D i.e., taken to be uniform and infinite along the *y* coordinate. (*B*) Dimensionless substrate concentration as a function of position 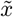 at different times, displaying both the inner and outer solutions, 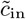 and 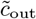, for an initial dimensionless colony width 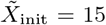 and system width 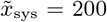, for the case where the front has been forced to be flat. The substrate penetrates only a small distance into the colony before it is completely consumed, and thus, growth is localized to a thin surface layer. (*Inset*) Evolution of the advancing planar front 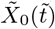 as a function of time as given by Eq. (9); points show specific times, while the black curve shows the numerical solution for all times. (*C,D*) Color plots of the dimensionless substrate concentration inside the bacterial colony, 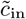, the cellular velocity vector field 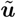 (arrows), and contours of the growth pressure field 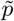 for an initially sinusoidally-perturbed colony surface with (*C*) a stable wavenumber, 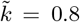, at time 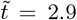, and (*D*) the same but for a perturbation with the most unstable wavenumber, 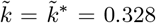 at time 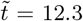. Both cases are obtained for an initial colony width 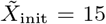, system width 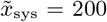 (same as in *B*), and initial perturbation amplitude 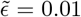. (*E*) Short-time perturbation growth rate 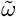 as a function of wavenumber 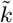 for the slab geometry with 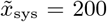 and different values of the initial colony width 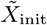. The dots correspond to the wavenumbers of the fronts shown in panels *C* and *D*. (*Inset*) Perturbation amplitude *Ã* as a function of time 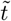 obtained from linear stability analysis (solid lines) and full numerical simulations (dots) for the two cases in panels *C* and *D*.

Our model is further detailed in *Materials and Methods*. We consider both a slab geometry, to more readily elucidate the relevant physics, and a cylindrical geometry, to directly compare to the experiments. We therefore adopt Cartesian and cylindrical coordinates, respectively, to describe the evolution of the growing colony, whose surface is located at *x* = *X*(*y,z,t*) for a slab-shaped colony (Fig. 2*A*), and at *r* = *R*(*z, θ, t*) for a cylindrical colony (Fig. 1*B*). The mass, momentum, and substrate conservation equations then read

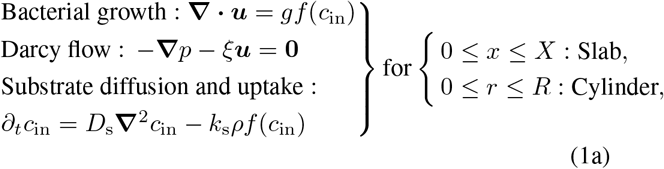

and are coupled to substrate diffusion outside the colony:

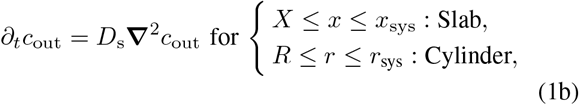

where ***x*** = (*x, y, z*), ***u*** = (*u_x_, u_y_, u_z_*) for a slab-shaped colony, and ***x*** = (*z, r, θ*), ***u*** = (*u_z_, u_r_, u_θ_*) for a cylindrical colony. Here, *g* is the maximal bacterial growth rate, *c*_in_(***x**, t*) and *c*_out_(***x**, t*) denote the substrate concentration inside and outside the colony, respectively, *f* (*c*_in_) = *c*_in_/(*c*_half,s_ + *c*_in_) is the Michaelis–Menten function reflecting the substrate-dependence of consumption relative to the characteristic concentration *c*_half,s_ [101, 102], and is an effective cell-matrix friction coefficient that reflects the proportionality between the gradient in growth pressure driving colony expansion and the speed with which it expands. Additionally, *x*_sys_ and *r*_sys_ describing the size of the overall system are given by the outermost width and radius of the hydrogel matrix, respectively.

As boundary conditions, we impose surface stress balance, substrate continuity, and the kinematic condition, i.e. *∂_t_ **x***_surf_ · ***n*** = ***u*** · ***n***, which relates the front velocity and the local velocity of the colony at the moving front, *x* = *X*(*z, y, t*) and *r* = *R*(*z, θ, t*), where ***x***_surf_ and ***n*** are the parametrization and the unit normal vector of the moving surface, respectively. We also impose a constant substrate concentration, *c*_f,s_, at the outermost system boundary, *x* = *x*_sys_ and *r* = *r*_sys_.

### Parameters characterizing 3D colony growth

To distill the relevant physics from our theoretical model, we first reduce the number of parameters in Eq. (1) by employing dimensional analysis and selecting appropriate quantities to scale the system of equations—as detailed further in the *SI Appendix.*

First, given that colony growth is a slow process (see e.g., Fig. 1*B*), we assume that the spatial profile of substrate rapidly adjusts to any change in the colony due to growth. We verify the validity of this assumption *a posteriori*, as further detailed in the *SI Appendix*. Moreover, as discussed previously, because substrate uptake is much faster than substrate diffusion at the scale of the overall colony (Table I), *c*_in_ ~ 0 inside the colony. Thus, there are two distinct regions in the colony: the substrate-depleted interior, and an actively growing boundary layer at the colony surface whose thickness is much smaller than *R*. In this surface layer, substrate diffusion and uptake are always in quasi-steady state balance—again yielding the substrate penetration length into the colony 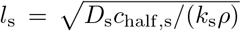 introduced earlier. This penetration length sets the characteristic length scale *l*_c_ = *l*_s_. We expect that *c*_in_ ≪ *c*_f,s_, *c*_half,s_ inside this boundary layer, as the substrate source is far from the colony surface in the experiments, i.e. *r*_sys_ ≫ *R*. Therefore, in what follows, we consider the limit *c*_in_ ≪ *c*_half,s_, where the Michaelis–Menten function simplifies to 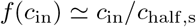.

As the main mechanism driving the expansion of the bacterial colony is growth mediated by substrate consumption, the characteristic scales for time *t*_c_ and front velocity *u*_c_ are obtained from the balance between mass conservation and bacterial growth in Eq. (1a), as well as the kinematic condition Eq. (5) of the front, which sets *u*_c_ = *l*_s_/*t*_c_. The natural time scale *t*_c_ is the inverse of the mean growth rate of cells within the growing front. In the substrate-limited regime, this mean growth rate is ~ *gc_s_/c*_half,s_, where *c*_s_ is the characteristic substrate concentration scale within the front. The latter can be estimated by matching the substrate gradient coming from diffusion outside the colony ~ *c*_f,s_/{*x*_sys_, *r*_sys_} with the gradient in the growing layer ~ *c_s_/l_s_*, yielding *c*_s_ ~ *l*_s_*c*_f,s_/{*x*_sys_, *r*_sys_}. Thus, *t*_c_ = *g*^-1^*c*_half,s_{*x*_sys_, *r*_sys_}/(*l*_s_*c*_f,s_).

Using these natural scales we obtain the following dimensionless parameters that govern colony growth:

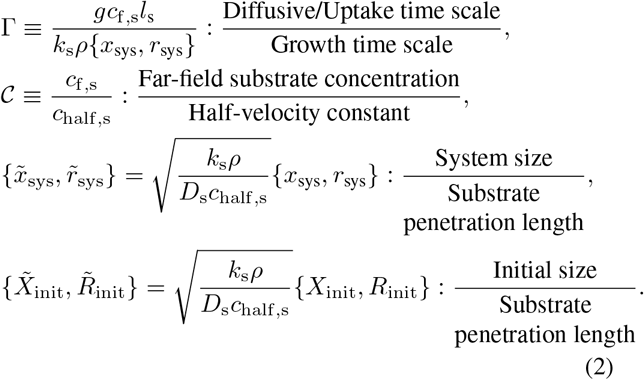

The first parameter Γ compares the characteristic diffusive time scale 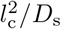, which is equivalent to the substrate uptake time scale *c*_half,s_/(*kρ*) since both diffusion and uptake are in quasi-steady state balance within the growing surface layer, to the characteristic time scale *t*_c_ for growth. As expected, in the experiments, the dimensionless parameter Γ ≪ 1, which means that growth occurs on a much slower time scale than the turnover rate of substrate due to uptake and diffusion within the colony. This allows us to consider that *c*_in_ is always at quasi-steady state.

The dimensionless parameter 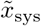 or 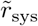, which describes a slab-shaped or cylindrical colony, respectively, is the ratio between the overall system size, *x*_sys_ or *r*_sys_, and the substrate penetration length *l*_s_. Since 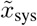 and 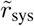 are large in our experiments, we expect that the qualitative growth behavior does not depend on the precise value of 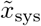 or 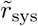.

The last dimensionless parameter 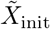 or 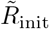 compares the initial width *X*_init_ or radius *R*_init_ of a slab-shaped (Fig. 2) or cylindrical (Fig. 1) colony, respectively, with the substrate penetration length *l*_s_. This parameter thus prescribes the initial condition in our theoretical model. In our experiments with cylindrical 3D colonies, 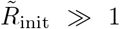 and Γ ≪ 1 as expected; hence, substrate becomes quickly depleted from the interior of the colony and growth is localized to the surface boundary layer. We therefore expect that the subsequent morphodynamics are independent of the initial colony shape. For this reason, we first consider the slab geometry schematized in Fig. 2*A*, which simplifies the analysis of the roughening instability, and then extend our analysis to the cylindrical geometry that describes the experiments (Fig. 1*A*).

### Continuum model recapitulates the key experimental findings

Having established that growth is localized to a thin layer at the surface of a colony, we next ask: how does this localization influence the subsequent colony morphodynamics? Because growth is localized to only a small fraction of the overall bacterial colony, we expect that the relative strength of random fluctuations in substrate availability and thus cellular growth is amplified—potentially engendering this morphological instability. To quantitatively explore this possibility, we first seek a 1D solution of the continuum model equations for the slab geometry (Fig. 2*A*) with a flat surface, and then examine its stability to shape perturbations.

To obtain a 1D solution of the flat-slab geometry, we assume that all the variables only depend on the coordinate 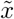 along which the front propagates: 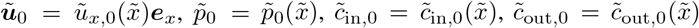, where the position of the moving interface 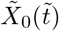 is only a function of time 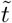, tildes denote dimensionless variables, and the nought subscripts denote the 1D flat-slab solution. As detailed in *Materials and Methods*, the solution is given by equations (8) and (9). Fig. 2*B* shows the resulting spatial profile of substrate concentration at different times 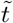 for the illustrative case in which the initial colony width 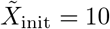. The initial leading edge of the colony is shown by the vertical line, and the inset depicts its subsequent position as a function of time as the colony grows. As in the experiments, because 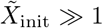, the substrate is mostly depleted inside the colony and only the surface boundary layer is able to grow.

Having established the 1D flat-slab solution, we next perform a linear stability analysis of this solution to assess whether this colony is intrinsically unstable to developing a rough surface (detailed in *Materials and Methods*). In particular, we analyze the rate 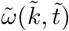 at which small periodic perturbations in the colony surface of wavenumber 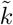, which corresponds to a wavelength 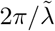, grow in time; thus, if 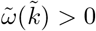, shape perturbations become amplified over time and the surface of the colony is unstable, whereas if 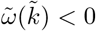, the perturbations become suppressed as the colony grows and its surface tends to be flat. The results for two exemplary values of 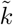 are shown in Figs. 2*C* and *D*. Intriguingly, the shorter-wavelength perturbation with larger 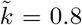 is stable, flattening out over time; by contrast, the longer-wavelength perturbation with a smaller 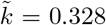 is unstable, causing the undulations in the colony surface to amplify as the colony grows.

This wavelength-dependence is summarized in Fig. 2*E*, which shows the perturbation growth rate 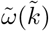 evaluated at the initial time for slab-shaped colonies of different initial widths 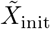; the simulations of Figs. 2*C* and *D* are indicated by the dots. As mentioned before, we expect that when 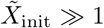 the colony width does not play a role in the stability of the growing front. In agreement with this expectation, we find that when the colony width substantially exceeds the substrate penetration length, the perturbation growth rate 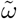 becomes independent of the width, as shown by the collapse of the curves for 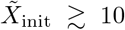. In this limit, the colony is morphologically unstable 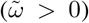 at small 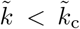, where 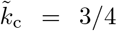 is a cut-off wavenumber—consistent with the results shown in Figs. 2*C* and *D*. Moreover, the perturbation growth rate takes on its maximal value 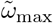 at a parameter-free most-unstable wavenumber 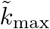, which corresponds to a value 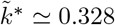 in the substrate-depleted limit 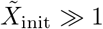. That is, our continuum model predicts that dense colonies of bacteria generically roughen as they grow—just as in the experiments. Moreover, this morphological instability is initiated by surface growth over a most-unstable wavelength 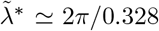 i.e., we expect λ* ≈ 19.2*l*_s_ ~ 20 – 200 μm in the experiments, comparable to the characteristic ‘floret’ size we observe for *E. coli*. Thus, despite its simplicity, our ‘active fluid’ continuum model of growing bacterial colonies captures the essential features of the roughening instability found experimentally.

### Continuum model reveals how substrate-limited growth causes colony surface roughening

What is the biophysical origin of this morphological instability? Our continuum model provides an answer; for example, compare Figs. 2*C* (stable) and *D* (unstable). In the latter unstable case, one sees that that the concentration of substrate at the peaks of perturbed colony surface is larger than in the valleys, resulting in faster subsequent growth at the peaks than in the valleys, which further amplifies the separation between the two. By contrast, in the former stable case, growth at the peaks transverse (in 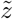) to the overall propagation direction 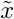 fills in the valleys, which reduces the separation between the two. Thus, differential access to substrate at different locations along the surface of a colony, which in turn imparts differences in the rate of cellular growth, is the primary mechanism driving colony roughening. Indeed, we can quantitatively express this intuition by analyzing the different contributions to the perturbation growth rate described by Eq. (12) in the *Materials and Methods*, for a given wavenumber *k*:

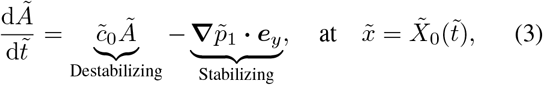

where 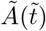 is the perturbation amplitude corresponding to the specific *k* being considered. The first term on the right-hand side corresponds to the driving mechanism of the instability: it reflects the fact that the concentration of substrate is higher in the peaks than in the valleys of the perturbed front. In particular, the substrate concentration of the 1D solution at the flat surface of an unperturbed colony, 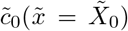 is positive for all values of the wavenumber 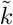 and initial colony width 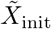—thus, all perturbations are amplified. By contrast, the second term on the right-hand side of Eq. (3) is the leading-order perturbation of the pressure gradient inside the colony, and reflects the stabilizing contribution from transverse growth. This stabilizing mechanism becomes very weak for large colonies 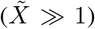, where substrate is depleted and growth mainly occurs in the 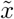 direction, but is able to overcome the destabilizing mechanism when 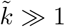. This behavior is in stark contrast to that of bacterial colonies that are not substrate-limited in their interior and thus grow uniformly at the same rate—for which all wavelengths become stable at sufficiently large colony width 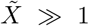 (see *Materials and Methods*), since in this case, transverse growth is able to smooth out any perturbation.

### When they are sufficiently large, growing cylindrical colonies exhibit the same morphological instability

In our experiments, the shapes of the 3D bacterial colonies do not correspond to a slab geometry, as their initial shapes are cylindrical (Fig. 1). However, the same biophysical ideas apply. We therefore employ the same theoretical framework and perform the equivalent linear stability analysis and time-dependent numerical simulations, but for the cylindrical case (as detailed in *Materials and Methods* and Figs. S19–S21). In this case, however, the shape perturbations to the colony surface can be either in the direction along the cylinder axis or along its azimuth—denoted by *z* or *θ* in Fig. 1*B*. Thus, we describe the harmonic perturbations using both the axial wavenumber 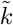 and azimuthal mode number *m*.

The results of the linear stability analysis are summarized in Figs. 3*A* and *B*. The color plots show the perturbation growth rate 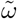 as a function of 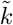 and the ratio between the initial colony radius and the substrate penetration length, 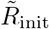, for two different azimuthal modes. Redder regions indicate increasingly unstable perturbations (larger 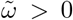), while blacker regions indicate increasingly stable perturbations (more negative 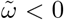). The orange dashed curve corresponds to the loci of the most-unstable mode 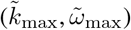 for each corresponding value of m.

**FIG. 3.**
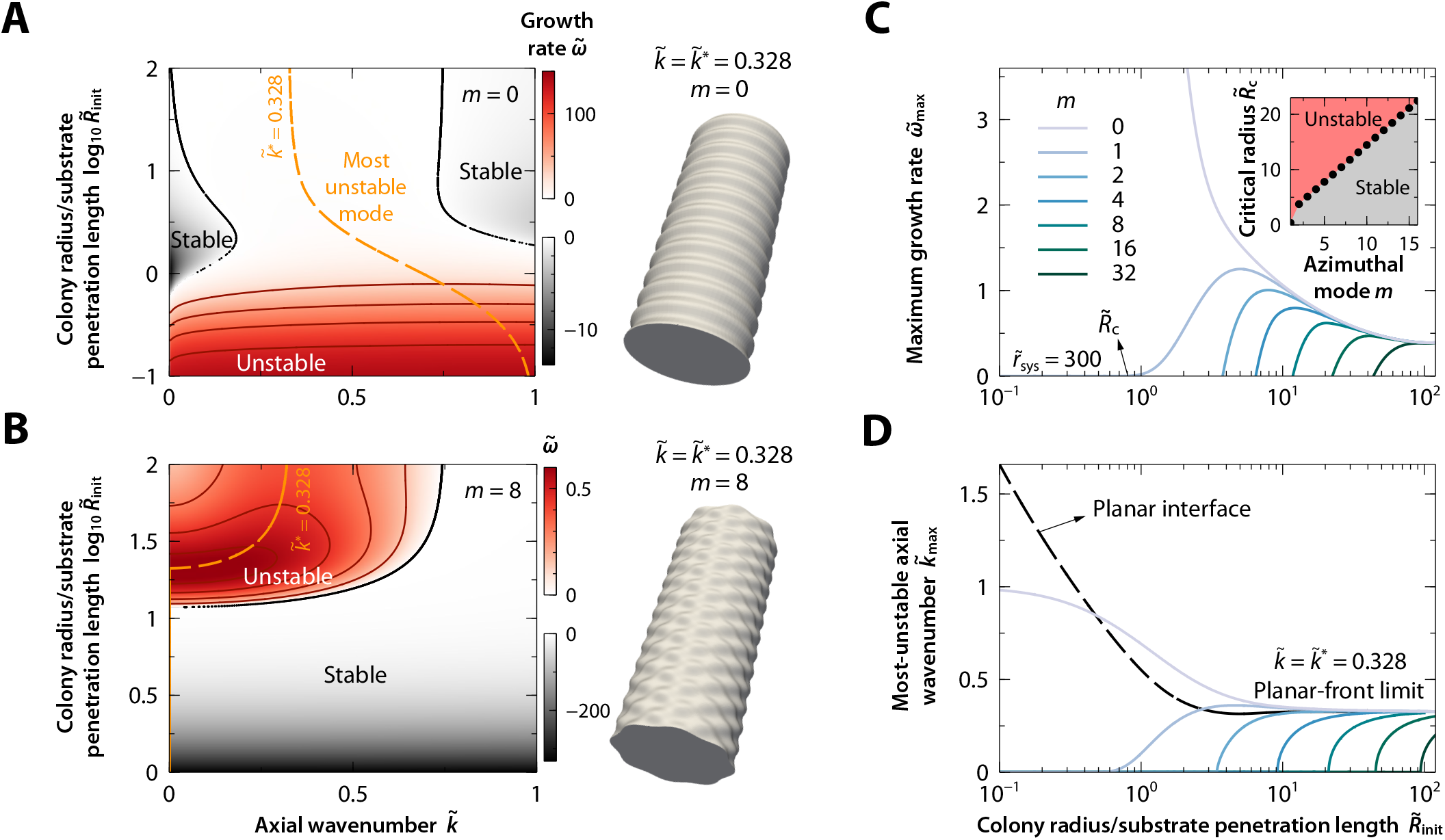
Continuum model of a growing cylindrical colony reveals a similar roughening instability. (*A,B*) (Left) Color plots of the smalltime growth rate 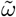 of perturbations as a function of the axial wavenumber 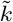 and the dimensionless ratio 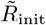 between the initial colony radius and the substrate penetration length, for dimensionless system radius 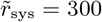. The azimuthal mode is *m* = 0 (axisymmetric perturbation) in *A* and *m* = 8 (non-axisymmetric perturbation) in *B*. The orange dashed curves indicate the loci of the most-unstable mode at short times, indexed by 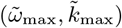. When 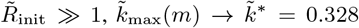 as found for a planar slab geometry. (Right) 3D renderings of a colony at the indicated values of *m* and 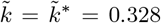, which corresponds to the most unstable mode for a planar slab. (*C*) Short-time maximum growth rate of perturbations 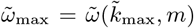 as a function of 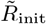 for different azimuthal modes *m*. (*Inset*) Critical radius 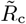 beyond which each non-axisymmetric mode becomes unstable. For 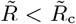 all modes are initially stable, i.e. decaying with growth. (*D*) Most-unstable wavenumber 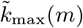 as a function of 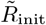 for the same values of *m* shown in *C*.

First, we examine the case of axisymmetric perturbations (*m* = 0). As shown in the top of Fig. 3*A*, large colonies 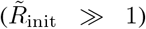 are unstable to perturbations at small axial wavenumbers (large wavelengths), with a most unstable wavenumber 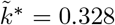—just as in the slab geometry, as expected. This most unstable mode is shown in the 3D rendering to the right of panel *A*. As shown in the bottom of Fig. 3*A*, with decreasing 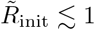, fluctuations in the growing surface layer increasingly dominate the morphodynamics, and all axial wavenumbers are increasingly unstable, as expected. Intriguingly, however, small-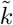 modes become stabilized for intermediate colony sizes 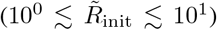—which likely reflects a non-trivial interplay between substrate availability and colony surface growth in these geometries.

Next, we study the case of non-axisymmetric perturbations, taking *m* = 8 as an example. As shown in the bottom of Fig. 3*B*, in this case, all axial wavenumbers are instead *stable* at small 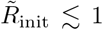—indicating that the stabilizing influence of transverse growth overcomes the roughening instability. However, as expected, large colonies 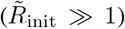 are again unstable to small-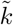 perturbations, but stable to large-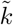 perturbations, with a most unstable wavenumber 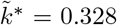—just as in the slab geometry and the *m* = 0 case. This most unstable mode is shown in the 3D rendering to the right of panel *B*. Thus, when colonies become large enough for growth to be localized to their surface, the roughening instability becomes independent of colony size or geometry.

To further compare the influence of axisymmetric (*m* = 0) and non-axisymmetric (*m* ≠ 0) perturbations, we examine the variation of the growth rate 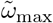 with 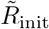 for the most unstable mode, indicated by the orange dashed curves in Figs. 3*A* and *B*. The results are shown in panel *C* for different azimuthal mode numbers *m*; the corresponding most unstable axial wavenumber 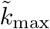 is shown in panel *D*. As expected, we find that the results for both axisymmetric and non-axisymmetric perturbations converge to the same curve when 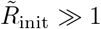 (see the collapse of the curves in the right of *C–D*), which corresponds to the size-independent curve obtained for a slab-shaped colony shown in Fig. 2*E*. This collapse again corroborates the finding that, when colonies become large enough for growth to be localized to their surface, the roughening instability appears to become independent of colony size or geometry. Moreover, for each non-axisymmetric mode, there is a critical dimensionless colony radius, 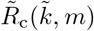, below which the growing colony is stable for all axial wavenumbers 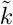, i.e., 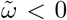, as shown in the inset to Fig. 3*C*—again indicating that the stabilizing influence of transverse growth overcomes the roughening instability for these modes. Indeed, the data in the inset indicate that this critical radius varies as 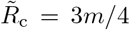, which agrees with the cut-off wavenumber obtained for a planar front, 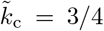. Hence, modes with a larger value of *m* need larger values of 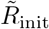 to merge into the planar slab limit. That is, more non-axisymmetric modes become unstable when the radius of the colony increases; these grow with the same rate and a parameter-free most-unstable wavelength corresponding to the slab geometry, i.e. 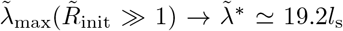, or equivalently 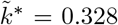. We confirm this behavior in which more non-axisymmetric modes become unstable via full numerical simulations (Figs. S20–S21). Further investigating these interesting morphodynamic features that arise due to the interplay between substrate availability and cellular growth, mediated by the geometry of the colony surface, will be a useful extension of our theoretical framework.

Taken together, our theoretical and simulation results for cylindrical colonies imply that when growth is localized to the surface of a colony—as is inherently the case when the colony becomes sufficiently large and dense—a generic morphological instability arises, independent of the initial colony size or geometry, and without requiring a specific nutrient environment or cell type, just as in our experiments. This instability is characterized by the growth of perturbations over many scales, but has a most-unstable wavelength whose size also agrees well with our experimental observations.

## DISCUSSION

Despite the ubiquity of bacteria in natural 3D environments, the morphodynamics of colonies growing in 3D has remained largely unexplored. One reason is the challenge of visualizing bacteria in 3D media, e.g. soil or tissues and organs, which are usually opaque. We overcame this challenge by performing experiments in transparent granular hydrogel matrices. Strikingly, we found that large, dense colonies growing in 3D undergo a generic roughening instability, adopting the same characteristic broccoli-like morphology independent of variations in cell type and environmental conditions. We elucidated the onset of the instability using a minimal continuum ‘active fluid’ model that incorporates the diffusion and uptake of a substrate essential for growth, coupled to cellular proliferation. In particular, we found that the *sine qua non* condition for the morphological instability is substrate depletion in the interior of the colony, causing the colony to expand via growth only at its surface. As schematized in Fig. 4, this surface growth amplifies perturbations in the overall colony shape—which inevitably arise in complex settings. In particular, we find there exists a universal most-unstable wavelength λ* ≃ 19.2*l*_s_, where *l*_s_ is the substrate penetration length into the colony. Hence, our experimental and modeling results reflect a key difference between growing in 2D and 3D in terms of access to critical substrates: while 2D colonies may access the substrates needed for growth from the third dimension and only roughen when growth is limited to a metabolically-active peripheral layer [30, 32, 33, 52, 91–93, 103–105], large 3D colonies inevitably become internally substrate-limited even under globally nutrient-replete conditions, eventually guaranteeing rough growth at their surface.

**FIG. 4.**
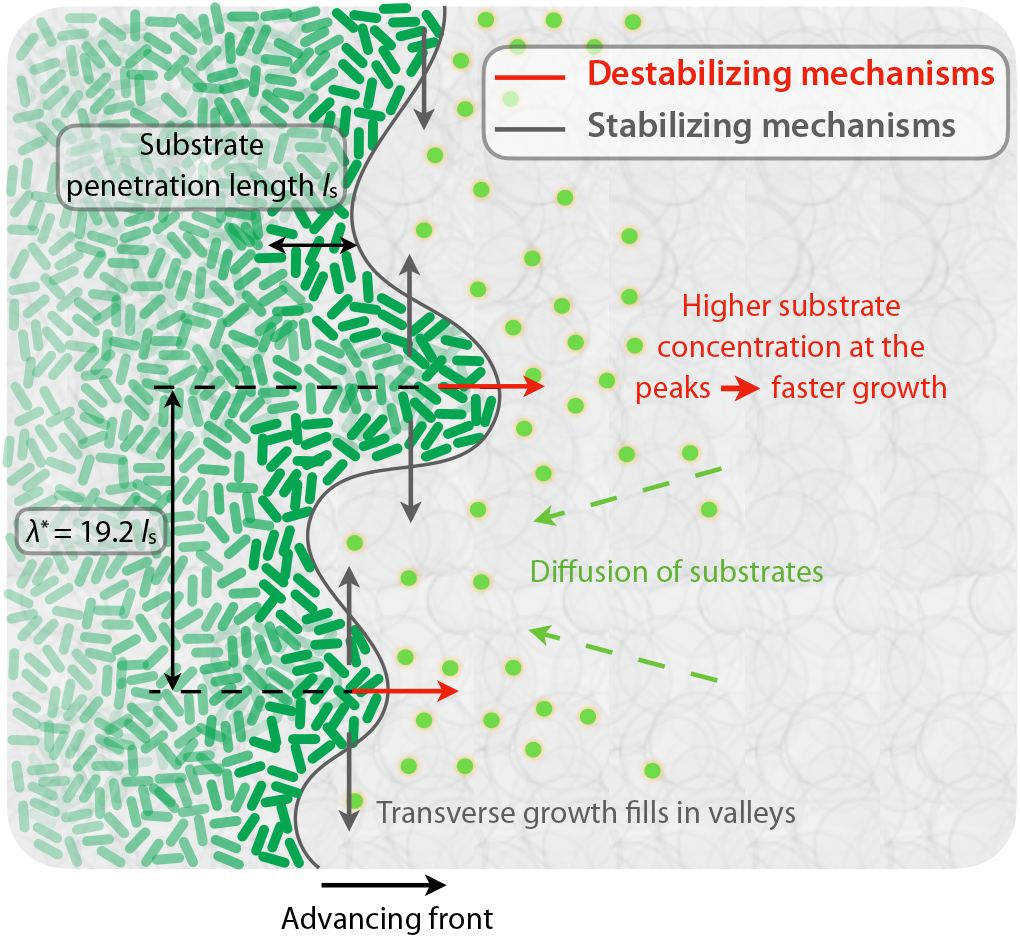
Biophysical mechanisms by which a growing colony is stabilized (gray) or destabilized (red). Substrates diffuse from the surroundings and are consumed by the cells in the colony (green), resulting in subsequent growth and outward colony expansion. The competition between substrate diffusion and consumption results in a small penetration length, causing growth to be localized at the surface of the colony. Growth transverse to the overall outward expansion direction fills in valleys and is thus stabilizing, while the increased availability of nutrients at peaks enables them to grow faster and is thus destabilizing. The competition between these mechanisms establishes the most-unstable wavelength λ* that emerges from our continuum model.

### Scaling behavior of 3D growing colonies

Surface growth is known to cause roughening in diverse processes, ranging from the growth of snowflakes and agglomeration of colloids to the formation of conductive dendrites that impede the operation of batteries [89, 90, 103, 106–117]. Similar roughening also arises in the growth of mammalian cell clusters when they are in nutrient-depleted spaces [76, 100, 118]. Despite the strikingly different underlying physical details, many of these processes exhibit remarkably universal growth dynamics and morphologies that can be described using the celebrated Edwards-Wilkinson (EW) or Kardar-Parisi-Zhang (KPZ) continuum growth models (or variations thereof) [89, 90, 103, 106–117]. Hence, to explore if growing 3D bacterial colonies share some of these universal characteristics, we computed the power spectrum *S*(*k,t*) and the local roughness *w*_loc_(*L,t*) of the colony surface profiles. We found similar results across the multiple cell types and environmental conditions tested, suggesting that our results reflect generic features of growing 3D bacterial colonies. As expected, the majority of the power corresponds to length scales around the most-unstable wavelength λ*/*l*_s_; however, as in many other growing systems, the spectra also exhibit a power-law decay, *S*(*k,t*) ~ *k*^-*ν*^, with *v* ≈ 7/3 at large *k*—reflecting the multi-scale nature of the surface roughening. Indeed, in analogy to the transfer of energy in classic inertial turbulence, where kinetic energy at large scales is dissipated by viscous forces at the smallest scales via an *energy cascade*, our results suggest that differential access to essential substrates and pressure-driven growth sets a range of most “energetic” wavelengths, with power transferred to smaller and smaller length scales down to the size of a single bacterium. Moreover, as in many other growing systems, the local roughness scales with the length *L* of the analyzed region as *w*_loc_ ~ *L^α^*—in our case, with *α* ≈ 0.7. The measured power-spectra-decay and roughness exponents show excellent agreement with the prediction of so-called Family-Vicsek dynamic scaling in 1D, *v* = 1 + 2*α*, which indicates that the surface of growing colonies is self-similar [89]. Thus, analogous to many other growing systems, 3D bacterial colonies adopt universal, self-similar, fractal shapes as they grow, independent of their starting geometry. Furthermore, the fact that the relation between the local roughness exponent and the power spectrum corresponds to 1D Family-Vicsek scaling supports the idea that the geometry or dimension of the front is not relevant for determining the stability or characteristics of the roughening process, when growth is localized to the surface.

However, the measured local roughness exponent *α* ≈ 0.7 is notably different from the value *α* =1/2 that is characteristic of the EW and KPZ universality classes (and which corresponds to an undirected random walk). Hence, growing 3D colonies appear to become rougher than many other growing systems that are well-described by the KPZ model. Values of *α* larger than 1/2 have been found in other systems (both non-living and living), typically when quenched disorder is present [89]—for instance in fluid flows through porous environments [109–112, 117, 119], burning fronts [113], directed percolation [114, 120–122], or in bacterial colonies growing on 2D agar gel [91, 103–105], for which case the origin is still under debate [91, 105]. What mechanisms could explain the large roughness exponent in our experiments? One possibility is that it reflects unstable growth associated with the coupling between cellular growth and substrate diffusion/uptake, as predicted by linear stability analysis of our model, and as described for other reaction-diffusion systems employed to model 2D growing colonies [91, 105]. The fact that the experimental power spectra exhibit a range of wavelengths where the power takes maximum values may be indicative of unstable growth during the expansion of the 3D colony. However, another possibility is that the large value of *α* reflects the influence of the granularity of the hydrogel matrices, which could impart additional heterogeneity in cellular growth due to, e.g., slight variations in pore sizes and hydrogel deformations. We neglected such complexities from our model for simplicity; however, we note that such forms of disorder are indeed thought to give rise to anomalously large growth exponents [89, 91, 103–105, 109–113, 117, 120–126]. Building on our work to study the influence of these complexities will be an important direction for the future.

### Strengths and limitations of our theoretical model

Given the complexities inherent in the experiments, and the simplifications made in the linear stability analysis of our active fluid model, can we still make quantitative comparisons between experiments and theory? Indeed, as summarized below, our theoretical analysis captures the three essential features of the experimental observations: (i) The width of the actively-growing layer of cells at each colony surface, (ii) the time at which colonies begin to roughen, and (iii) the characteristic wavelength of the rough morphology that colonies eventually produce.

We expect that (i) is set by the penetration length of the growth-limiting essential substrate, which we estimated using our theory as 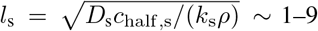 μm (Table I). Our experiments yielded a comparable value: in all the experiments, independent of other cell properties, substrate availability, or the initial colony geometry, we directly visualized a thin layer of fluorescent bacteria (which indicates cells are metabolically active) extending over a distance ≲ 20 μm from the colony front. Indeed, consistent with our findings, previous theoretical work on 2D growing colonies has indicated that a rough front arises when growth occurs only in a sufficiently thin layer located close to the advancing front, i.e. under globally scarce nutrient conditions in the case of a 2D colony [30, 32, 33, 52, 91–93].

The time scale for the colony to expand by the width of the surface layer, which we expect sets (ii), is predicted by our theory to be *t*_c_ ~ 6–50 h (Table I). Consistent with this expectation, in all the experiments, we found that colonies become rough after several hours—which also agrees well with our full numerical simulations in the substrate-depleted limit (Figs. S20–S21).

Regarding (iii), our experiments reveal that the power spectrum of shape fluctuations *S*(*k, t*) plateaus for *k* ≪ 2*π*/λ_c_ and exhibits a power-law decay for *k* ≳ 2*π*/λ_c_, where the characteristic crossover wavelength is λ_c_ ≈ 60 – 600 μm for *E. coli.* As shown in Fig. S22 *A*, our model similarly implies a plateau in the power spectrum for small 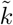, for which the growth rate 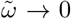 (with noise injecting additional power), and a powerlaw decay for large 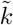—with the characteristic wavelength demarcating the transition between these regimes predicted to be λ_c_ = (8*π*/3)*l*_s_ ~ 10 – 70 μm, roughly comparable to the experimental values. This agreement between the experiments and theory is noteworthy, given that the experimental spectra do not correspond to the earliest stages of growth where the linear stability analysis applies. Moreover, the experiments revealed a similar roughening instability in four different species of bacteria having different natural habits and physiologies, highlighting the generality of this phenomenon as predicted by the theory. Finally, we also note that recent *in vivo* studies have reported similar rough morphologies of bacterial aggregates growing in diverse 3D environments, such as snow [127], microgels prevalent in marine environments [128, 129], mouse urine [130], and sludge [131, 132].

Nevertheless, our model has limitations and does not quantitatively reproduce the entirety of the experimental findings—suggesting refinements that could be considered in future work. For example, while the model correctly predicts a spectral power-law decay by introducing Gaussian white noise, the predicted exponent *ν* =1 (Fig. S22) is at odds with the experimental value *ν* ≈ 7/3. Along the same lines, the linear regime of our model, in which some wavelength perturbations decay and other grow without bound, cannot directly account for the saturation of the power spectra observed in the experiments at long time, where *S*(*k, t*) reaches a fixed curve. Nonlinearities could account for these discrepancies—for instance, by damping unstable growth and increasing the magnitude of the power-law exponent e.g. via an “energy cascade” analogous to that in turbulence as discussed above. To explore the effect of these nonlinearities, direct numerical simulations of the noisy version of Eq. (7) could be conducted, introducing either thermal or quenched noise, possibly along with renormalization group techniques [89]. Finally, we note that while our quantitative analysis focused on experiments with *E. coli*—given that the physical parameters describing these cells (Table I) are well-characterized, that the cellular fluorescence enabled high-resolution imaging via confocal microscopy, and that this strain does not appreciably secrete extracellular materials—we also observed a similar morphological instability for the biofilm-formers *V. cholerae*, *P. aeruginosa*, and *K. sucrofermentans*. However, we note that the characteristic ‘floret’ wavelength for these latter three species is noticeably larger than for *E. coli* (see Figs. S10–S14, S23). This difference could reflect the added influence of the extracellular polymeric matrix secreted by the cells as they grow, which potentially leads to complexities that are not taken into account in our minimal model. Other experimental effects that are also not contained within the model—but may nevertheless play a role—include the granularity and deformability of the hydrogel matrices noted earlier, the mechanical properties of and cellular orientations within the bacterial colony, variations of cellular density within the growing layer, and limitations on the space and amount of nutrient available to the growing colony.

### Implications for microbiology

Our work has demonstrated that, much like snowflakes, large, dense, 3D bacterial colonies cannot avoid developing rough surfaces as they grow. What are the biological implications of this morphological instability? The highly-branched fractal colony shapes studied here are characterized by large surface area-to-volume ratios, which likely promote the exposure of metabolically-active cells at the colony surfaces to their surroundings. Thus, we speculate that the roughening of growing bacterial colonies could be harmful to them by increasing susceptibility to antibiotics [58–60] or phages [61], but could also be beneficial in the long term by providing more space for mutants to grow, thereby supporting genetic diversity [53, 55, 56, 64–69] and potentially enabling colonies to be more resilient against environmental changes [62, 63].

Cell-cell signaling might also be affected by this morphological instability, since different regions of the rough surface of a colony could become decoupled from each other—which in turn could also affect overall colony function and internal colony morphology [133]. Previous work on 2D growing colonies has shown metabolic cooperation between cells growing at the periphery of the colony and those located in the interior––ion channels enable long-range electric signaling between both regions of the colony, giving rise to spatially propagating metabolic waves [134–139]. Hence, although our results suggest that the core of 3D colonies plays a passive role in the colony morphodynamics, the interaction and signaling between the metabolically-active growing layer at the surface and the core may have important consequences for the global function and structure of the colony, and this also constitutes an important direction for future research.

More broadly, given our finding of a morphodynamic instability that emerges generically from the coupling between diffusion and uptake of essential substrates for cellular growth in 3D, we expect that similar behavior could arise in other growing systems, with possible implications for the evolution of form and function [50, 55, 140–142]. Indeed, similar 3D patterns of rough growth have been observed in other living systems, namely in multicellular clusters of *Saccharomyces cerevisiae*, usually referred to as ‘snowflake yeast clusters’, or in aggregation clusters of the green alga *Volvox carteri*. Recent research has shown that the sizes and morphologies of these clusters are crucially determined by growth and crowding-induced mechanical stresses coupled to environment conditions [143–147]. These works have suggested the tantalizing idea that the morphological adaptation of large 3D clusters to environmental constraints could provide insight into the origins of multicellularity. Altogether, our experimental approach and findings, as well as our theoretical framework, could provide a foundation for these promising future avenues for research.

## MATERIALS AND METHODS

### Preparing hydrogel matrices

The 3D granular hydrogel matrices are prepared following our prior work [34, 35] by dispersing dry granules of crosslinked acrylic acid/alkyl acrylate copolymers (Carbomer 980, Ashland or Carbopol 980NF, Lubrizol) in liquid cell culture media.

We use EZ Rich (Teknova Inc.), a defined rich medium, for the experiments with *E. coli;* 2% Lennox LB (Sigma Aldrich) for the experiments with *V. cholerae* and *P. aeruginosa;* and Mannitol-Yeast Extract-Peptone (MYP) medium (Sigma Aldrich) for the experiments with *K. sucrofermentans.* The components to prepare the EZ Rich are are mixed following manufacturer directions; specifically, the liquid medium is an aqueous solution of 10X MOPS Mixture (M2101), 10X ACGU solution (M2103), 5X Supplement EZ solution (M2104), 20% glucose solution (G0520), 0.132 M potassium phosphate dibasic solution (M2102), and ultrapure milli-Q water at volume fractions of 10%, 10%, 20%, 1%, 1%, and 58%, respectively. All components except glucose are combined and then autoclaved. After the medium cools, sterile glucose is added. The MYP medium is made of D-Mannitol (25 g/L), Yeast Extract (5 g/L), and Peptone (3 g/L) all mixed in ultrapure milli-Q water and autoclaved.

In all cases, the hydrogel granules are then homogeneously dispersed by mixing each dispersion for at least 2 h at 1600 rpm using magnetic stirring or in a stand mixer (Hamilton Beach 730C Classic DrinkMaster Mixer) for 2 minutes. We also adjust the pH to 7.4 by adding 10 M NaOH to ensure cell viability. For the experiments with *K. sucrofermentans*, 2,3,5-Triphenyltetrazolium chloride (TTC), a red dye for cells, is also added to the swollen hydrogel particles at 5 mg TTC per 10 mL of swollen hydrogel matrix. In all cases, the swollen hydrogel granules, ~ 5 to 10 μm diameter, pack tightly against each other, resulting in a jammed matrix; the individual hydrogel granules have an internal mesh size of ~ 40 to 100 nm, as we established previously [88]. Thus, small molecules (e.g., amino acids, glucose, oxygen) can freely diffuse throughout the medium, while the cells themselves are confined in the interstitial pore space between hydrogel granules.

### Characterizing hydrogel matrices

Rheological properties of these matrices are measured in an Anton Paar MCR 301 or 501 shear rheometer, as previously described [34, 88]. We load ~ 2 – 3 mL of a given hydrogel matrix into the 1 mm gap between roughened 50 mm-diameter parallel plates. We measure storage and loss moduli as a function of frequency using small-amplitude oscillatory rheology, with a strain amplitude of 1% and frequencies between 0.01 to 1 Hz. For all hydrogel matrices used in this work the storage modulus *G*’ is larger than the loss modulus *G*”, indicating the matrices are jammed elastic solids. At higher strain rates, the matrices fluidize, indicating that they are yield-stress materials; we quantify this behavior using unidirectional shear measurements in which we measure the shear stress as a function of applied shear rate. At low shear rates, the shear stress is nearly constant and independent of shear rate, indicating a nonzero yield stress (~ 1 – 100 Pa) characteristic of a solid material. At higher shear rates, however, the shear stress follows a power-law dependence on shear rate, indicating that the solid matrix becomes fluidized—the individual hydrogel particles rearrange with respect to each other, and the medium yields [86]. This features allows for 3D printing of bacterial colonies in the hydrogel matrices.

We characterize the sizes of the interstitial pores between hydrogel granules as previously described [34, 88]. Specifically, we homogeneously disperse 10^-4^ wt% of 100 nm or 200 nm-diameter carboxylated polystyrene fluorescent nanoparticles (FluoSpheres, Invitrogen, Carlsbad, Ca) within the interstitial pore space between hydrogel granules by gentle mixing. Because the tracer particles are larger than the mesh size of the hydrogel granules, they thermally diffuse through the interstitial pore space. For each hydrogel matrix, we track the motion of 50 to 200 tracer particles on a Nikon A1R+ inverted laser-scanning confocal microscope. We use a custom written script in MATLAB employing the classic Crocker-Grier algorithm to track the center of mass of the particles using a peak finding function with subpixel precision in the micrographs. From the particle tracking, we calculate the mean square displacement, MSD. The tracer MSDs exhibit free diffusion in the pore space at short length and time scales, and then transitions to subdiffusive scaling at sufficiently large length and time scales because of pore-scale confinement. Thus, we identify the transition length scale at which the MSD becomes subdiffusive, and estimate the local pore size d as being the sum of the MSD transition length scale and the tracer size. Repeating this measurement for many different tracers dispersed at random locations through the pore space yields the pore size distribution. TIn particular, we obtain the complementary cumulative distribution function 1-CDF(*d*); 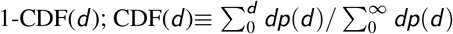 where *p*(*d*) is the number fraction of pores having a characteristic size *d*. To determine the mean pore size, we then fit an exponential ~ exp(–*d/D*) to the decay of 1-CDF(*d*) for each medium and report the mean pore size as 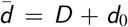 where *d*_0_ is the largest pore size with CDF= 0.

### 3D-printing bacterial colonies within hydrogel matrices

Our experiments used five different bacterial strains: *E. coli* W3110 (motile) and ΔflhDC (non-motile), both of which constitutively express GFP; *V. cholerae* JY030 [148]; *P. aeruginosa* PAO 1 ΔfliC; and *K. sucrofermentans* ATCC-700178. In each experiment, we first prepare a dense suspension of cells in the liquid cell culture medium used to swell the granular hydrogel matrix, and then use this suspension as the inoculum that is 3D-printed into the matrix, following our previous work [34, 35].

For the experiments with *E. coli* and *P. aeruginosa*, to obtain the suspension for the initial inoculum, we prepare an overnight culture of cells in LB media at 30°C, then incubate a 1% solution of this culture in fresh LB media for 3 h until the optical density reaches ~ 0.6, and then re-suspend this culture in the specific liquid cell culture medium used to swell the granular hydrogel matrix (EZ Rich for *E. coli* and LB for *P. aeruginosa*) at a concentration ~ 9 × 10^10^ cells mL^-1^. For the experiments with *V. cholerae*, to obtain the suspension for the initial inoculum, we first grow the cells overnight on a 2% Lennox LB agar plate at 37°C, then inoculate the cells into 3 mL of liquid LB with glass beads and grow them with shaking at 37 °C for 5-6 hours until mid-exponential phase, and then similarly re-suspend this culture in LB at a similar concentration~ 9 × 10^10^ cells mL^-1^. For the experiments with *K. sucrofermentans*, to obtain the suspension for the initial inoculum, we first prepare a culture in stationary phase in MYP at 30 °C for 4 days, then incubate a 1% solution of this culture in 3 mL fresh MYP with 0.2% cellulase at 30 °C for 4 days, and then similarly re-suspend this culture in fresh MYP at a similar concentration ~ 9 × 10^10^ cells mL^-1^.

We then 3D-print the inocula in the hydrogel matrices. For each experiment with *E. coli*, ~ 4 mL of the hydrogel matrix is confined in a transparent-walled glass-bottom petri dish of 35 mm diameter and 10 mm height, and we use a pulled glass capillary with a ~ 100 – 200μm-wide opening as an injection nozzle for the colony. For each experiment with *V. cholerae* or *P. aeruginosa*, ~ 20 mL of the hydrogel matrix is confined in a transparent tissue culture flask or ~ 3 mL of the the hydrogel matrix is confined in a transparent plastic macro cuvette, and we use a 20-gauge needle as an injection nozzle for the colony. For the *E. coli*, *V. cholerae*, and *P. aeruginosa* experiments, a motorized translation stage subsequently drives the injection nozzle to trace out a pre-programmed 3D cylindrical shape within the hydrogel matrix at a constant speed of 1 mm s^-1^. As the injection nozzle moves through the medium, it locally rearranges the hydrogel grains and gently extrudes the inoculum into the interstitial space using a flow-controlled syringe pump at 50 μL h^-1^, which corresponds to a gentle shear rate of ~ 4 to 36 s^-1^ at the tip of the injection nozzle. As the nozzle continues to move, the surrounding hydrogel grains rapidly densify around the newly-introduced cells, re-forming a jammed solid matrix [85, 86, 149] that compresses the cellular suspension until the cells are close-packed to an approximate density of 0.95 × 10^12^ cells mL^-1^. Moreover, as we showed in our previous work [34, 35], this process does not appreciably alter the properties of the hydrogel packing and is sufficiently gentle to maintain the viability and motility of the cells. The resulting cylindrical colony serves as the starting point of the experiments detailed in this paper. In all cases, we ensure that the colony is placed at least ~ 500 to 1000 μm away from any boundaries enclosing the hydrogel matrix. For the experiments with *E. coli*, after 3D printing, a thin layer of 1 – 2 mL paraffin oil is placed atop the top surface of the hydrogel matrix to minimize evaporation while allowing unimpeded oxygen diffusion; for the experiments with *V. cholerae* and *P. aeruginosa*, we accomplish this by sealing the tissue culture flasks with their caps. For the experiments with *K. sucrofermentans*, instead of using an injection nozzle mounted on a motorized stage, we slowly pipette 10 μL of the inoculum in ~ 1.5 mL of the hydrogel matrix and then cover the top of the matrix it with another ~ 1.5 mL of MYP hydrogel matrix, all in a sealed plastic macro cuvette.

### Inoculating bacterial colonies on the hydrogel surface

For the experiments testing *E. coli* growth on the surface of a granular hydrogel matrix, we use a transparent-walled glass-bottom petri dish 35 mm in diameter and 10 mm in height. Initially, the glass bottom (20 mm in diameter and ~ 1 mm in height) is filled with the hydrogel matrix and its top surface is smoothed out. Next, the edge of a glass cover slip (~ 100 μm in thickness) is dipped in a bacterial suspension at a concentration of 8.6 × 10^10^ cells mL^-1^. This edge of the cover glass is then gently touched to the surface of the hydrogel matrix to define an initial cylindrical inoculum on the surface. We then maintain the petri dish within a temperature- and humidity-controlled environmental chamber (Memmert HPP110) at 30 ± 1 °C and 90% relative humidity to reduce evaporation.

### Imaging growth of bacterial colonies

For the experiments with fluorescent *E. coli*, we use a Nikon A1R+ inverted laser-scanning confocal microscope maintained at 30 ± 1 °C to acquire vertical stacks of planar fluorescence images separated by 2.58 μm in depth. For the experiments with *V. cholerae*, *P. aeruginosa*, and *K. sucrofermentans*, we use a Sony *α*6300 camera with a Thorlabs 6.5X Zoom Lens to acquire color images. Between imaging time points the samples are placed in a 37 °C incubator (for *V. cholerae*) or a 30 °C (for *P. aeruginosa* and *K. sucrofermentans*). As time progresses, the colonies continue to grow until at long times (≳ 100 h), when colony expansion ceases presumably due to complete substrate depletion.

### Measurement of the characteristic Michaelis-Menten nutrient concentration of *E. coli*

To estimate *c*_half,s_, we measure the growth rates of wild-type *E. coli* (strain background W3110) in serial dilutions of EZ Rich media. In particular, we inoculate an overnight culture at a ratio of 1:100 into 200 μL of liquid medium and measure growth by optical density on a Biotek Epoch 2 Plate Reader for 24 hours at 30 °C for 6 replicates. Diluted nutrient media have constant concentrations of MOPS buffer (Teknova M2101) and potassium diphosphate (M2102), but diluted concentrations of glucose (Teknova G0520), amino acids (Teknova M2105), and nucleic acids (Teknova M2103) such that the concentration of L-serine, which we take as an example metabolized nutrient, spans 0.1 μM–10 mM. For each growth curve, the maximum growth rate *g* is extracted by fitting a linear curve to log(OD_600_) of the exponential growth regime. Then, the value of *c*_half,s_ is obtained by fitting the growth rates for each L-serine concentration to the Michaelis-Menten function using non-linear least squares. The fit value of *c*_half,s_ is 38 μM with 95% confidence bounds ranging from 29.6 μM to 46.8 μM.

### Measurement of the maximal growth rate of *V. Cholerae*

To obtain *g*, we measure the growth rates of the *V. Cholerae* C6706str2 in Lennox LB. We inoculate an overnight culture at a ratio 1:100 into 50 mL of LB with glass beads and measure the optical density on a WPA Biowave Reader for 10 hours at 37 °C for 3 replicates.

### Boundary conditions of the continuum model

The system of equations Eq. (1) is complemented with the following boundary conditions:

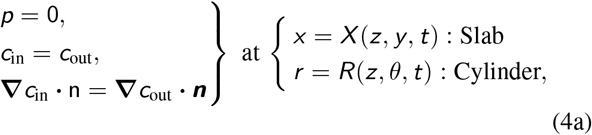

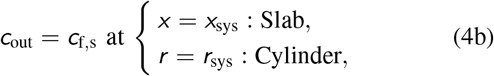

where ***n*** is the unit vector normal to the bacterial colony surface. In our experiments we do not impose a constant far-field source of nutrients, but we expect our theoretical results to be accurate due to the separation of time scales between colony growth and nutrient diffusion and uptake.

Moreover, we impose the following kinematic condition at the growing front:

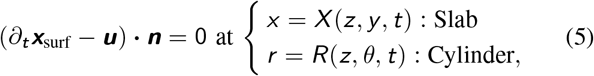

where ***x***_surf_ is the parametrization of the surface of the colony, i.e. the position vector representing any point at the surface.

### Non-dimensionalization

To reduce the number of parameters in Eq. (1), we nondimensionalize the system of equations. To this end, we choose the characteristic scales obtained in the main text and introduce the following dimensionless variables for the position vector, time, and the velocity, pressure, and nutrient fields, respectively,

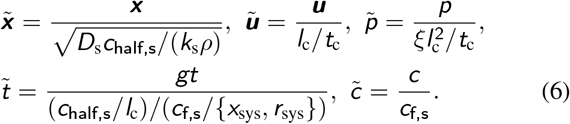

Using the above characteristic scales, the dimensionless bacterial mass, momentum, and substrate conservation equations read, respectively

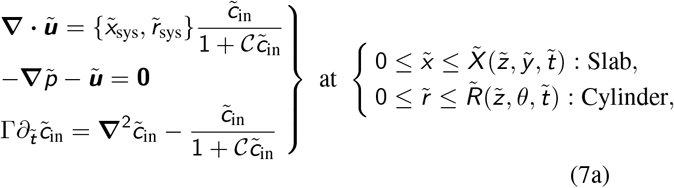

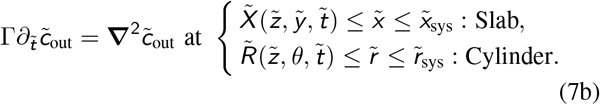

In the limit where *c*_in_ ≪ *c*_half,s_, which is a good approximation under our assumption of the limiting substrate being depleted inside the bacterial colony, i.e. {*X*, *R*} ≫ *l*_s_, we can approximate 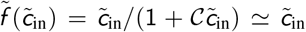, where 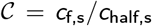. All the analytical calculations are performed using this approximation, as well as the full numerical simulations unless otherwise specified.

To conduct numerical simulations, we integrate Eq. (7) imposing the dimensionless version of the boundary conditions Eq. (4) and Eq. (5), together with axially periodic boundary conditions, i.e. 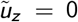at 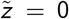 and 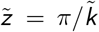, where 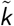 is the dimensionless axial wavenumber of perturbations. To trigger the instability, the interface is slightly perturbed by a harmonic disturbance at 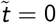, for the cylindrical geometry, 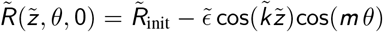, and the slab geometry, 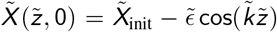 for 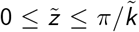, where 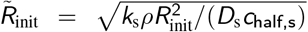 compares the initial radius of the cylinder *R*_init_ = *R*(*t* = 0), with the substrate penetration length *l*_s_ (and equivalently 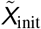 for a slab-shaped colony), and 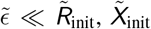 is the relative amplitude of the initial small perturbation.

### 1D solution for slab-shaped and cylindrical colonies

We seek a one-dimensional (1D) solution of the dimensionless version of the system of equations for the slab geometry in Fig. 2*A*, with the growth front constrained to be flat and assuming that 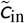 is at steady state, i.e. Γ ≪ 1. This solution is a necessary starting point for our subsequent linear stability analysis. Specifically, we assume that all the variables only depend on the coordinate 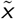, along which the front propagates, 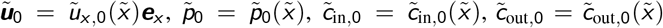, where the position of the moving interface 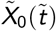 is only a function of time 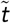. We impose the boundary conditions given by Eq. (4), which yields the solution

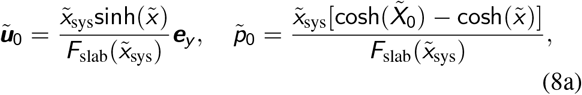

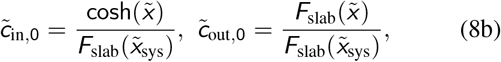

where 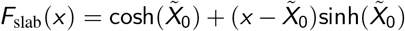. The position of the colony’s interface is obtained via the kinematic condition in Eq. (5), which simplifies to

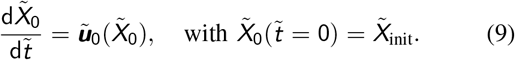

This first-order ordinary differential equation does not admit an analytical solution, thus we solve it numerically by standard methods.

As for the slab geometry, we can also calculate the 1D solution for a cylindrical geometry. This solution is a necessary starting point for the linear stability analysis of a growing cylindrical colony. To this end, we assume that the colony grows radially maintaining a perfect cylindrical shape, thus the velocity, pressure, and nutrient concentration fields only depend on the radial coordinate 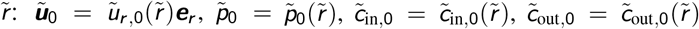. Hence, imposing the dimensionless boundary conditions given by Eq. (4), and the kinematic condition, Eq. (9), to obtain the position of the cylindrical interface as a function of time, 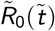, we obtain the following 1D solution:

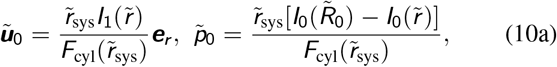

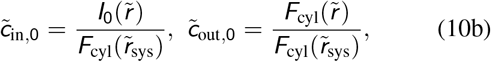

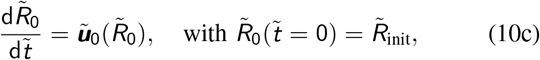

where for the cylindrical colony 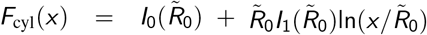, and *l_n_*(*x*) is the nth-order modified Bessel function of the first kind.

### Linear stability analysis of growing 3D bacterial colonies

To elucidate if a 3D colony is intrinsically unstable to surface roughening when nutrient is depleted inside and growth is confined to the advancing front, we perform linear stability analysis on the 1D solutions for the slab and cylinder geometries obtained previously. To this end, all the variables are perturbed around the 1D solutions by small-amplitude harmonic disturbances, and the time-dependent perturbations are decomposed as Fourier wave-like modes, for the cylindrical case defined by the axial wavenumber 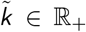 and the azimuthal number 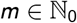:

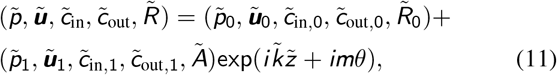

where 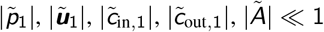. Introducing Eq. (11) into the system Eq. (7), allows us to solve for 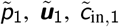, and 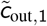, using the dimensionless version of the linearized boundary conditions Eq. (4). Since the colony is expanding with time, and we expect that the perturbation evolves on the same time scale as the moving front, we cannot derive a standard dispersion relation between an exponential growth rate and the wavenumber 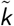 and azimuthal modes *m*. Instead, the evolution of the interface perturbation 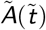 is obtained from the linearized kinematic condition Eq. (5) at leading order:

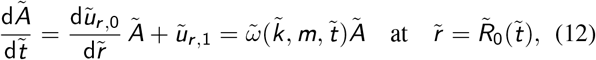

where we have taken advantage of 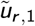 being proportional to 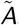, to define 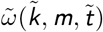 as the time-dependent growth rate. This allows us to solve Eq. (12), which yields,

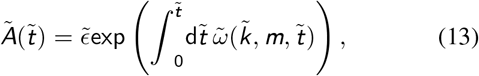

where 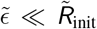 is the initial small amplitude of the perturbation, and the lengthy expression for 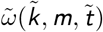 is omitted here for the sake of simplicity (see *SI Appendix*). The linear stability analysis of a slab-shaped colony is performed in the same manner (see *SI Appendix*).

### Derivation of the structure factor

Here we derive the theoretical structure factor after obtaining the deterministic growth rate of perturbations 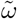. To this end, the evolution equation for the perturbation amplitude 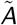 in Fourier space, Eq. (12), is modified by introducing Gaussian white noise 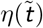, resulting in the following Langevin equation,

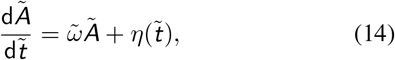

where 〈*η*〉 = 0, and 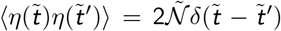, and 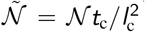 denoting the dimensionless intensity of the noise, which we assume to be independent of 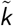. After solving the amplitude equation, the structure factor is obtained as follows,

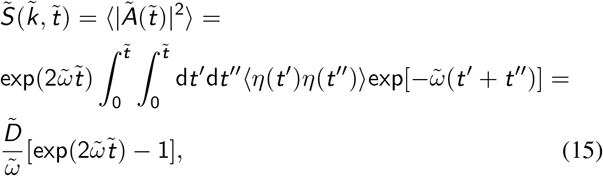

where we have considered an initial structure factor corresponding to a flat front in a slab geometry, and corresponding to a perfect cylinder in the cylindrical geometry. Knowing that 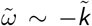 when 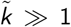, as shown in Fig. 2*E*, we obtain 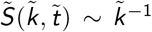 in the same limit (Fig. S22A). Moreover, at long wavelengths the structure factor grows as 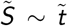 when 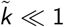. The roughness of the surface can be obtained by integrating the structure factor over the wavenumber,

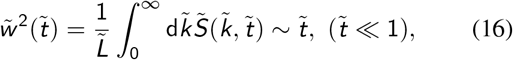

where 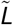 is the dimensionless length of the cylinder, or the slab-shaped colony. Hence, 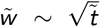, at short time (Fig. S22*B*). The same scaling is found in random deposition or in the Edwards-Wilkinson equation at short time.

### Bacterial colonies growing uniformly

Here we consider the limit where the entire bacterial colony grows at a uniform rate. This limit applies when *c*_half,s_ ≪ *c*_in_, i.e. when substrate is sufficiently abundant to be saturating everywhere in the colony (which is the opposite limit of our experiments at sufficiently long time, where 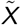 or 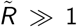). Hence, in this scenario, growth is not limited by nutrient or oxygen concentration, and the equations of motion become decoupled from the substrate dynamics. For uniform growth, if we employ the appropriate characteristic scales, i.e. *t*_c_ = *g*^-1^, *l*_c_ = *R*_init_, *u*_c_ = *gR*_init_, and 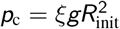, the dimensionless equations are parameter-free and read:

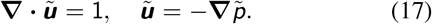

The 1D solutions for the planar and cylindrical growing fronts then read:

#### Slab geometry

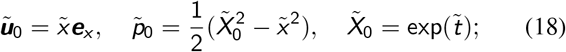

#### Cylindrical/circular geometry

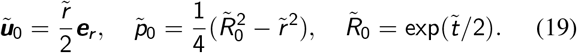

We now assess the linear stability of these 1D solutions, considering the cases of a purely planar front, a circular colony expanding radially, and a cylindrical colony growing in the radial direction. Following the same procedure detailed before, we obtain the leading-order eigenfunctions of the pressure field 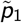 and velocity field 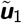, and then use Eq. (12) to derive the evolution of the perturbation amplitude.

#### Slab-shaped colony

For the planar growing front, the evolution equation for the perturbation amplitude 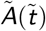 given by Eq. (13) reads

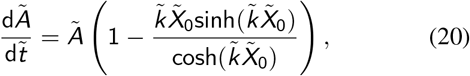

which admits the following analytical solution:

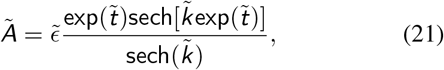

where 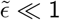 is the small initial value of the amplitude.

Since in the case of uniform expansion, growth is proportional to the local width of the colony, we expect that wavelengths shorter than the initial colony width 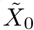 will be stable as growth-pressure-driven expansion transverse to the overall growth direction will be effective at filling in the valleys of the perturbation. Indeed, expanding the term on the right-hand side of Eq. (20) in powers of 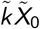, we obtain a reasonably accurate estimate of the cut-off wavenumber 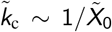 above which all modes are stable. Moreover, at long time, i.e. when 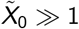, the critical wavenumber 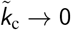 as 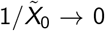, which means that all modes are stable at sufficiently long time, since the stabilizing contribution stemming from transverse growth becomes large enough to stabilize all perturbations. The stabilizing mechanism is transverse growth, which is taken into account by the second term on the right-hand side of Eq. (12), and becomes comparable to the growth perpendicular to the colony interface.

#### Circular colony

In the case of a planar circular colony growing uniformly, the perturbation amplitude also admits an analytical solution:

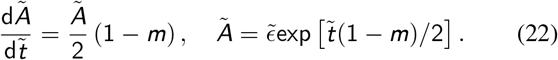

All modes are stable for all time except *m* = 1, which simply corresponds to a body translation and is therefore neutrally stable.

#### Cylindrical colony

If we now consider a cylindrical colony growing uniformly in the radial direction, whose 1D solution is given by Eq. (19), the equation for the perturbation amplitude is

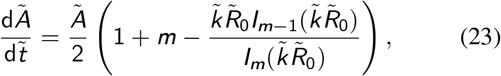

which does not admit an analytical solution. If we take the limit 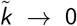, we recover the case of a 2D circular colony given by Eq. (22). To gain more insight into the stability of the cylindrical growing colony, we can expand the rightmost term in parentheses in Eq. (23) in powers of 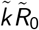, which yields 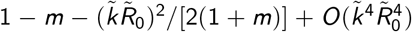. By inspection, we observe that non-axisymmetric modes *m* ≥ 2 are stable for all values of 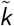 and 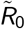, as in the case of the 2D circular colony. Regarding the axisymmetric modes, i.e. *m* = 0, we recover the same scenario obtained for a purely planar growing front, up to geometrical factors.

Finally, all these limits can be obtained from the complete model given in Eq. (7), where the growth of the colony depends on the local concentration of oxygen and nutrients, when 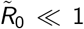, since substrates are then essentially uniform inside the colony.

### Numerical simulations

To perform 2D and 3D numerical simulations, all the dimensionless equations are written in weak form by means of the corresponding integral scalar product, defined in terms of test functions for the pressure 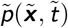 and nutrient concentration fields 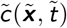. By using Green identities we obtain an integral bilinear system of equations for the set of variables and their corresponding test functions:

Pressure-driven growth:

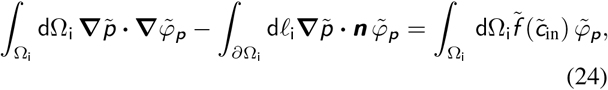

substrate diffusion and uptake:

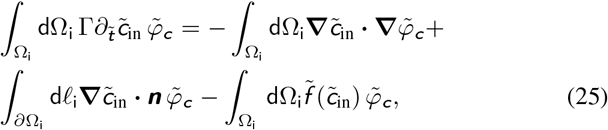

substrate diffusion outside the colony:

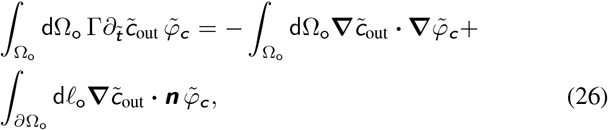

where 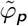 and 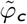 are the test functions for the pressure and substrate concentration fields, respectively, Ω_i_ and Ω_o_ are the domains inside and outside the bacterial colony, respectively, dΩ and d*ℓ* are the volume (3D)/surface (2D) and surface (3D)/line (2D) elements, respectively, and 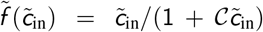 is the dimensionless Michaelis-Menten function. The velocity is calculated using Darcy’s law, 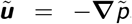, and the moving interface is computed according to the kinematic condition in Eq. (5), where nodes at the interface are advanced using the normal component of the velocity field. We impose the boundary and initial conditions specified in Eq. (4), together with the axially periodic condition. The equations are discretized using Taylor-Hood triangular/tetrahedral elements for the pressure and nutrient concentration fields, ensuring numerical stability. To account for the deformation of the domain we use the Arbitrary Lagrangian-Eulerian method implemented in COMSOL Multiphysics, where the mesh elements in both domains move according to Laplace’s equation for the change of variable between the material and the spatial frames of reference. Regarding the time-stepping, we employ a 4th-order variable-step BDF method. The relative tolerance of the nonlinear method is always set below 10^-6^.

## Data availability

The data and codes that support the plots and findings of this study are available from the authors upon reasonable request.

## ACKNOWLEDGMENTS

A.M.C. acknowledges support from the Princeton Center for Theoretical Science and the Human Frontier Science Program through the grant LT000035/2021-C. R.K.B. acknowledges support from the Presidential Postdoctoral Research Fellows Program. H.N.L. acknowledges support from the Lidow Independent Work/Senior Thesis Fund at Princeton University. This material is also based upon work supported by a National Science Foundation Graduate Research Fellowship Program to A.M.H. under Grant No. DGE-2039656. N.S.W. acknowledges support from the NSF through the Center for the Physics of Biological Function PHY-1734030 and the NIH through grant R01 GM082938. S.S.D. acknowledges support from NSF grants CBET-1941716, DMR-2011750, and EF-2124863, as well as the Eric and Wendy Schmidt Transformative Technology Fund, New Jersey Health Foundation, and Pew Biomedical Scholars Program. We are grateful to Daniel Amchin, Alejandro Sevilla, Howard Stone, and Sankaran Sundaresan for thoughtful discussions, and Sebastián González La Corte for assistance with experiments using *P. aeruginosa*. We also thank the labs of Bob Austin, Bonnie Bassler, and Zemer Gitai for providing strains of *E. coli*, *V. cholerae*, and *P. aeruginosa*, respectively.

## Authors contributions

T.B. and S.S.D. designed the experiments; T.B. performed all experiments with *E. coli*; R.K.B. performed all experiments with *V. cholerae* and *P. aeruginosa*; T.B., R.K.B., A.M.H., and H.N.L. performed additional characterization experiments; R.K.B. and H.N.L. performed experiments with *K. sucrofermentans*; A.M.C., N.S.W., and S.S.D. designed the theoretical model and numerical simulations; A.M.C. performed all theoretical analysis and numerical simulations, as well as analysis of the experimental images; A.M.C., T.B., R.K.B., A.M.H., H.N.L., N.S.W., and S.S.D. analyzed the data; S.S.D. designed the overall project; A.M.C., N.S.W., and S.S.D. discussed the results and implications and wrote the manuscript.

## SI APPENDIX

### Dimensional analysis

We will first identify characteristic scales of length, time, pressure, velocity, and nutrient concentration, denoted as *l*_c_, *t*_c_, *p*_c_, *u*_c_, and *c*_c_, respectively, and we will then use these scales to non-dimensionalize the corresponding quantities in Eq. (3) of the Main Text. To choose the appropriate scales, we start with the kinematic equation relating the front speed with the local velocity inside the colony, which sets the following balance:

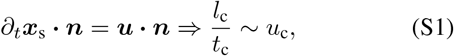

yielding the relation *l*_c_ = *u*_c_*t*_c_, so that both terms in the equation are of order unity. We then consider the nutrient conservation equation in the interior of the colony, whose different terms have the following orders of magnitude:

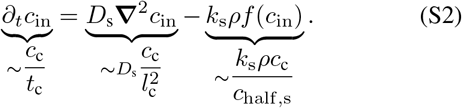

As illustrated by Fig. 1*B* in the Main Text, our experiments reveal that the growth of the colony is a slow process, thus we assume *a priori* that the growth of the colony is much slower than nutrient diffusion and uptake, whereby 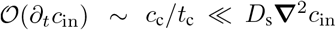 and *k*_s_*ρ*. We validate this assumption *a posteriori* using our parameter estimates, as described further in this section. Note that this assumption implies a separation of time scales such that the nutrient profile will rapidly adjust to any change in the colony due to growth. This separation of time scales is assumed in Eqs. (3c) and (3d) of the Main Text. Moreover, a natural length scale of the system is either the initial size of the colony, *R*_init_, or the system size *r*_sys_, i.e. the overall size of the hydrogel matrix where the bacterial colony is embedded. If we choose either of these lengths as the characteristic length scale and then consider the estimates in Table 1 of the Main Text, one can see that 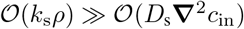, which yields *c*_in_ ~ 0 in the bulk of the colony. This result reveals two distinct regions inside the colony: the nutrient-depleted bulk, and an active growing boundary layer at the surface of the colony whose thickness is much smaller than the radius *R* of the colony, and where nutrient diffusion and uptake are in balance. Given that this growing region is the main cause of colony expansion, we choose its thickness as the characteristic length scale in our system of equations, i.e. 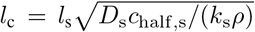, which is the substrate penetration length arising from the balance between nutrient diffusion and uptake.

Next, to determine the characteristic time scale, we examine the mass conservation equation describing growth as the driving mechanism of the moving front:

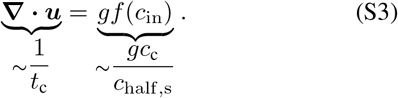

From the only possible balance in the above equation, the natural time scale *t*_c_ is the inverse of the mean growth rate of cells within the growing front. In the nutrient-limited regime, this mean growth rate is ~ *gc*_s_/*c*_half,s_, where *c*_s_ is the characteristic substrate concentration scale within the front. The latter can be estimated by matching the substrate gradient coming from diffusion outside the colony ~ *c*_f,s_/{*x*_sys_,*r*_sys_} (where *x*_sys_ is the size of the outer medium in the case of a slab-shaped colony) with the gradient in the growing layer ~ *c*_s_/*l*_s_, yielding *c*_s_ ~ *l*_s_*c*_f,s_/{*x*_sys_, *r*_sys_} and thus *t*_c_ = *g*^-1^*c*_half,s_{*x*_sys_, *r*_sys_}/(*l*_s_*c*_f,s_).

We now address the assumption that *∂*_*t*_*c*_in_ is much smaller than the remaining terms in Eq. (S2) by comparing their orders of magnitude, i.e. 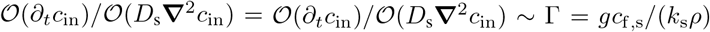, which is indeed a small parameter given the estimates in Table 1 of the Main Text, thus our choice of *t*_c_ is appropriate and self-consistent.

Finally, the characteristic pressure scale is obtained from the balance in Darcy’s law,

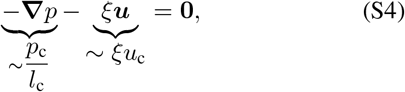

which yields *p*_c_ = *ξu*_c_*l*_c_ as the characteristic pressure scale.

### Linear stability analysis

To analyze the linear stability of slab-shaped or cylindrical bacterial colonies, we first obtain a one-dimensional (1D) solution of the dimensionless version of Eq. (9) in the Main Text, with the growth front constrained to be either flat or cylindrical, respectively (see *Materials and Methods* and Figs. S20, S22–23). Then, all the variables are perturbed around these 1D solutions by small-amplitude harmonic disturbances, and the time-dependent perturbations are decomposed as Fourier wave-like modes (which in the cylindrical case are defined by the axial wavenumber 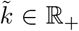 and the azimuthal number 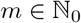):

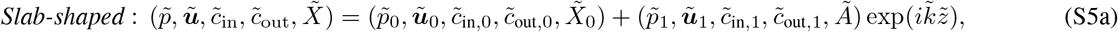

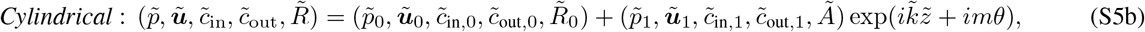

where 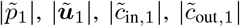 are the eigenfunctions of the small-amplitude perturbations, and *Ã* is the amplitude of the front with 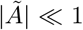. Introducing Eq. (S5) into Eq. (9) of the Main Text and retaining only leading-order terms allows us to solve for 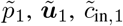, and 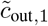, which for a slab-shaped colony yields:

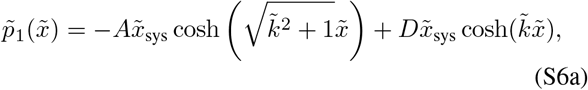

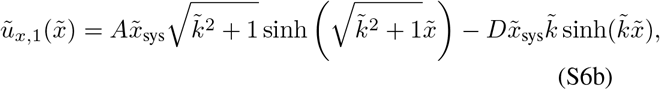

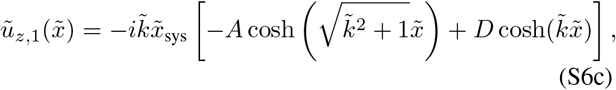

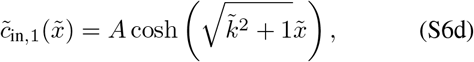

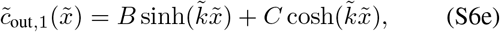

and for a cylindrical colony yields:

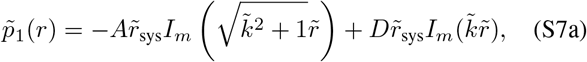

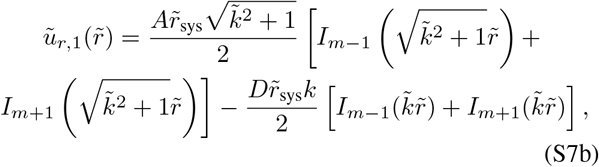

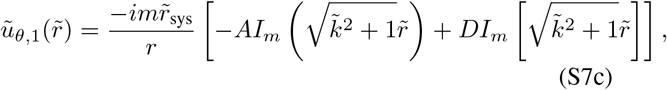

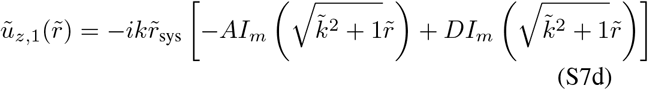

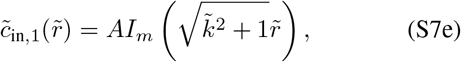

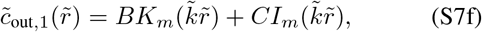

where *A, B, C*, and *D* are integration constant that we obtain by imposing the linearized boundary conditions given by Eqs. (6) and (7) of the Main Text. Due to their lengthiness, their expressions are omitted here. Axisymmetric modes are obtained for the case *m* = 0. Moreover, the eigenfunctions of an expanding circular 2D colony are recovered by taking the limit *k* → 0.

Finally, the growth rate of perturbations 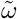 is obtained from the remaining boundary condition, i.e. the kinematic condition for the moving surface, which relates the local velocity of the colony ***u***, with the velocity of the moving front 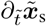 (see Eq. (7) in the Main Text), which upon linearization yields a first-order ODE for the perturbation amplitude *Ã*:

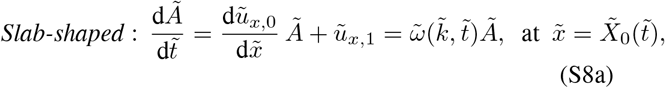

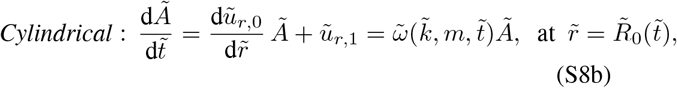

where we have taken advantage of 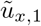 and 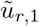 being proportional to *Ã* to define 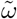 as the time-dependent growth rate. Since the colony is expanding with time, i.e. the 1D solution depends on time, and we expect that the perturbation evolves on the same time scale as the moving front, we cannot consider the front to be frozen and derive a standard dispersion relation between an exponential growth rate and the wavenumber *k* and azimuthal mode *m*, as in our case 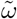 depends on time. Instead, the evolution of the interface perturbation 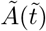 reads:

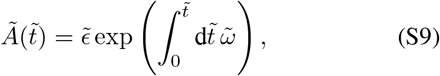

where 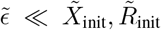 is the initial small amplitude of the perturbation. Hence, the growth rate of perturbations can be defined as:

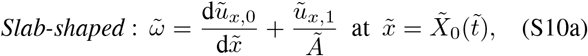

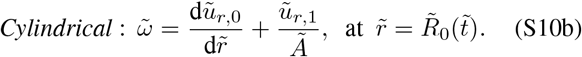

The expression for a cylindrical colony is omitted due to its lengthiness, but the growth rate of perturbations of a slabshaped colony reads,

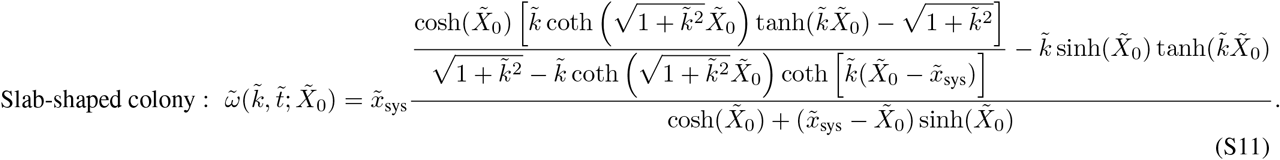

To deduce the main driving mechanism of the morphodynamical instability, we examine the different contributions to the time-dependent growth rate of the perturbation amplitude *Ã* for a slab-shaped colony, i.e. Eq. (S8a). To this end, we expand the local velocity of the colony at the surface, which we express using Darcy’s law ***u*** = – ∇*p*, as follows:

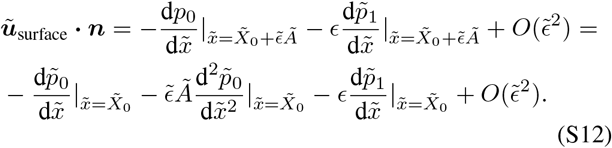

Hence, at leading order, the evolution of the perturbation amplitude 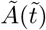 is given by the following ODE:

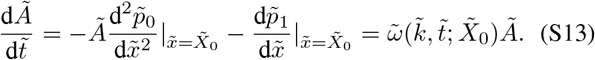

This equation is identical to Eq. (S8a), but expressed in terms of the pressure field rather than the local velocity field. The first term in the right-hand-side of Eq. (S13) arising from the 1D solution is not zero since the pressure gradient is not uniform, i.e. the pressure field is not harmonic due to the growth of the bacterial colony, and thus neither the 1D pressure drop nor the velocity field are constant. Indeed, we can write this contribution in terms of the nutrient concentration by combining Darcy’s law, 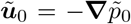, and the mass conservation equation, 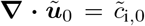, to yield 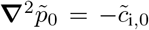, or equivalently 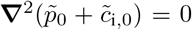, since we consider the steady-state nutrient concentration, i.e. 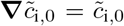. Hence, we can also write the equation for the perturbation amplitude *Ã* as follows,

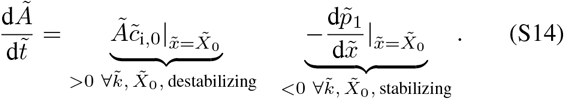

The first term in the right-hand-side of Eq. (S14) is independent of the wavenumber 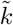, and thus is always destabilizing for every value of 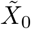 since the nutrient concentration is always positive. This term reflects the fact that peaks at the surface of the bacterial colony have more access to nutrients and oxygen than the valleys, and therefore grow faster—amplifying the shape perturbation; that is, differential access to nutrients is the main driving mechanism of the instability. The second term in the right-hand side is negative for every value of 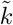 and 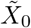, thus reflecting a stabilizing contribution. In particular, this term contains contributions arising from nutrient diffusion whose effect is to smooth nutrient gradients along the moving front, and also the pressure gradient associated with growth in such transverse direction to the overall growth direction, that also contributes to smooth the perturbations. To gain a better understanding of the latter mechanism, we explore a simplified case of the complete problem, in particular when the colony grows uniformly in space, i.e. with growth not limited by the local nutrient concentration.

### Uniform growth

To isolate the role of growth and its associated pressure gradient, we consider the limit where the bacterial colony grows at a rate that is spatially uniform and constant in time. This limit applies when *c*_half,n_ ≪ *c*, i.e. typically when nutrient is sufficiently abundant to be saturating everywhere in the colony, which is the opposite limit of our experiments where 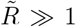. Hence, in this scenario, growth is not limited by nutrient or oxygen concentration, and the equations of motion become decoupled from the nutrient dynamics:

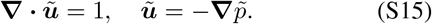

The 1D solutions for the planar and cylindrical configurations are then, respectively:

#### Slab geometry

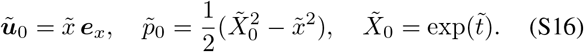

#### Cylindrical/circular geometry

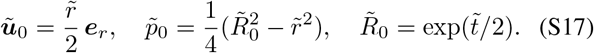

We now assess the linear stability of these 1D solutions, considering the cases of a purely planar front, a circular colony expanding radially, and a cylindrical colony growing in the radial direction.

### Uniform growth — Linear stability

Following the same procedure detailed before and in *Materials and Methods*, we obtain the leading-order eigenfunctions of the pressure field 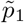 and velocity field 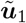, and then use Eq. (S13) to derive the evolution of the perturbation amplitude.

#### Slab-shaped colony

For the planar growing front, the evolution equation for the perturbation amplitude 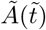 given by Eq. (S13) reads

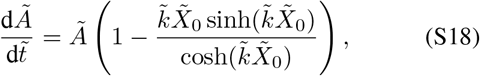

which admits the following analytical solution:

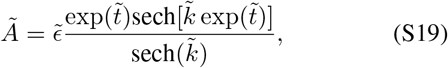

where 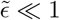 is the small initial value of the amplitude.

Since growth is proportional to the local width of the colony, we expect that wavelengths shorter than the initial colony width 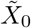 will be stable as growth-pressure-driven expansion transverse to the overall growth direction will be effective at filling in the valleys of the perturbation. Indeed, expanding the term on the right-hand side of Eq. (S18) in powers of 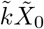, we obtain a reasonably accurate estimate of the cut-off wavenumber 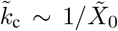 above which all modes are stable. Moreover, at long time, i.e. when 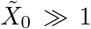, the critical wavenumber 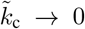 as 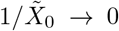, which means that all modes are stable at sufficiently long time, since the stabilizing contribution stemming from transverse growth becomes large enough to stabilize all perturbations.

#### Circular colony

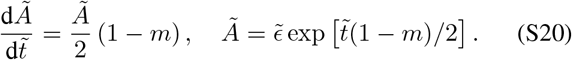

#### Cylindrical colony

If we now consider a cylindrical colony growing uniformly in the radial direction, whose 1D solution is given by Eq. (S17), the equation for the perturbation amplitude is

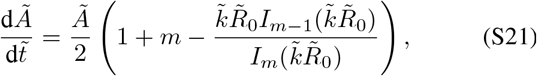

which does not admit an analytical solution. If we take the limit 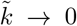, we recover the case of a 2D circular colony given by Eq. (S20). To gain more insight into the stability of the cylindrical growing colony, we can expand the rightmost term in parentheses in Eq. (S21) in powers of 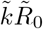, which yields 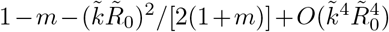. By inspection, we observe that non-axisymmetric modes *m* ≥ 2 are stable for all values of 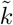 and 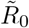, as in the case of the 2D circular colony. Regarding the axisymmetric modes, i.e. *m* = 0, we recover the same scenario obtained for a purely planar growing front, up to geometrical factors.

### Influence of nutrient and oxygen consumption

To determine whether the consumption of nutrients or oxygen influences our results, we modify Eq. (7) of the Main Text to consider that nutrient and oxygen uptake, as well as growth, depend on both the local concentration of oxygen and nutrient via Michaelis-Menten kinetics. Hence, the new system of equations for a cylindrical geometry reads:

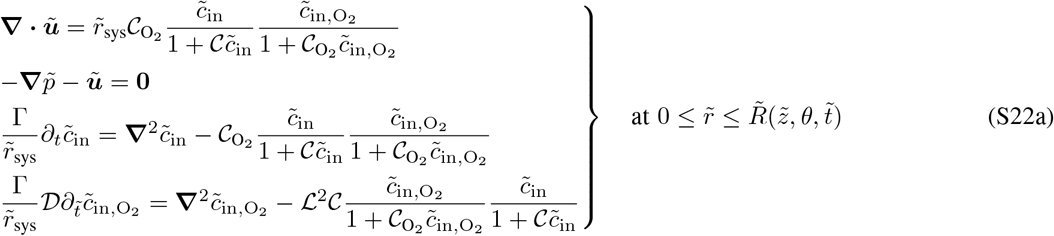

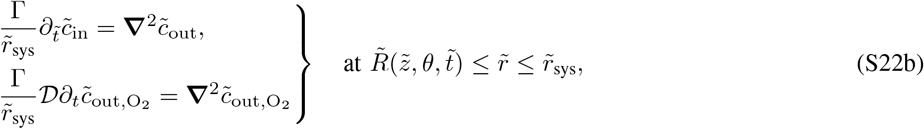

where 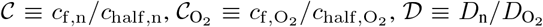 is the nutrient-to-oxygen diffusivity ratio, and 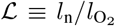 is the ratio between the nutrient and oxygen penetration lengths, the latter being 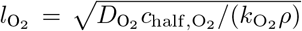. In particular, given that 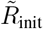 is sufficiently large, when 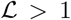 oxygen is limiting since it is depleted inside the colony, whereas when 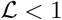 nutrient is limiting. Considering the estimates of Table 1 in the Main Text, we perform numerical simulations of the above system of equations for 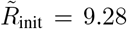, Γ = 0, 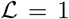, 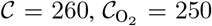, and 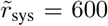. In particular, Fig. S16 shows the front position 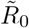 of the 1D solution as a function of time, computed using system Eq. (9) of the Main Text. We find nearly identical results to that of the simplified system of equations. Moreover, the linearization of Eq. (S22) yields a very similar system of equations to that obtained in the simplified case considered in the Main Text. Thus we expect the growth rate of perturbations and the physics of the linear regime not to change significantly with respect to the results obtained in the Main Text.

**Table S1.**
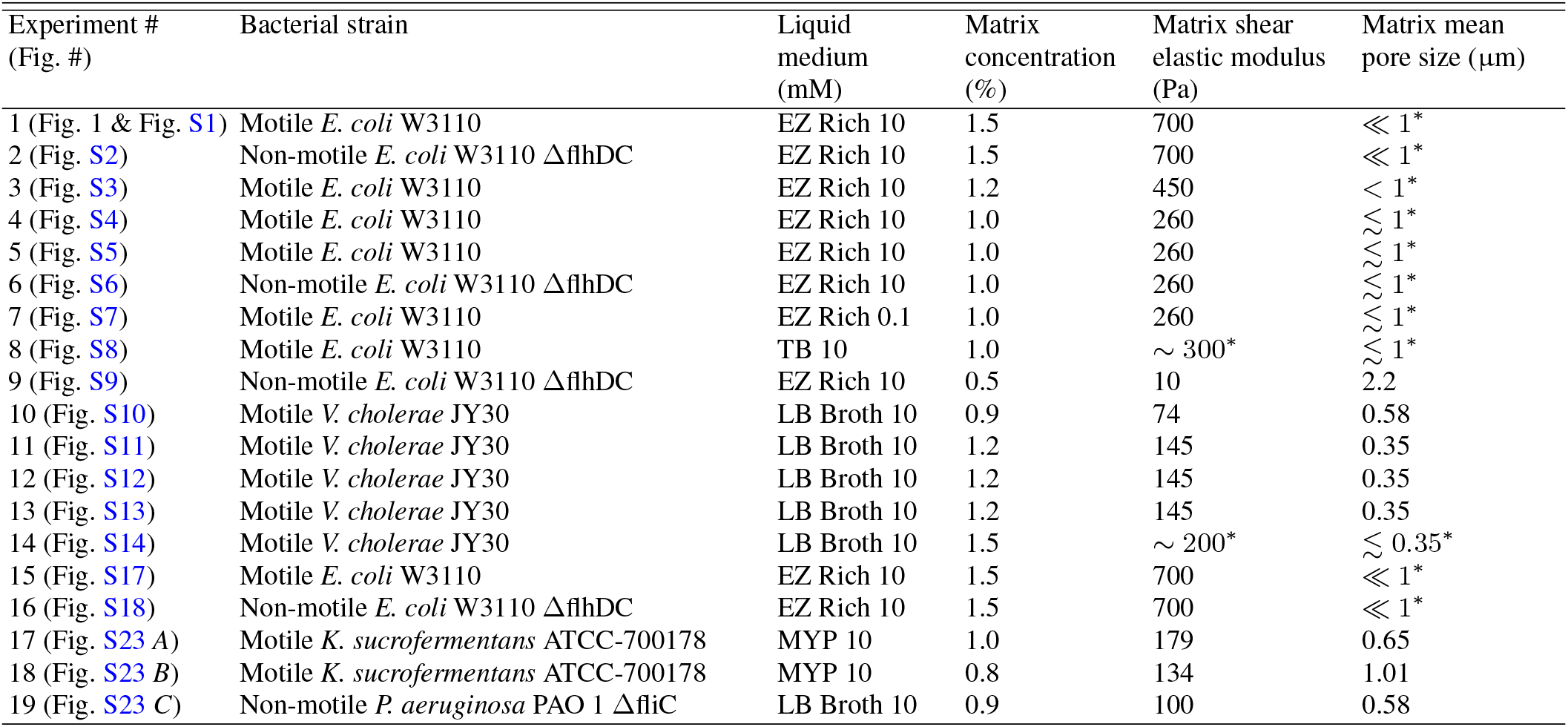
Summary of experiments. Values listed are either obtained from direct measurements, as described in *Materials and Methods*, or when indicated by * are estimated by extrapolating our current or previous measurements [34, 88] on hydrogel matrices prepared identically at different matrix concentrations.

**FIG. S1.**
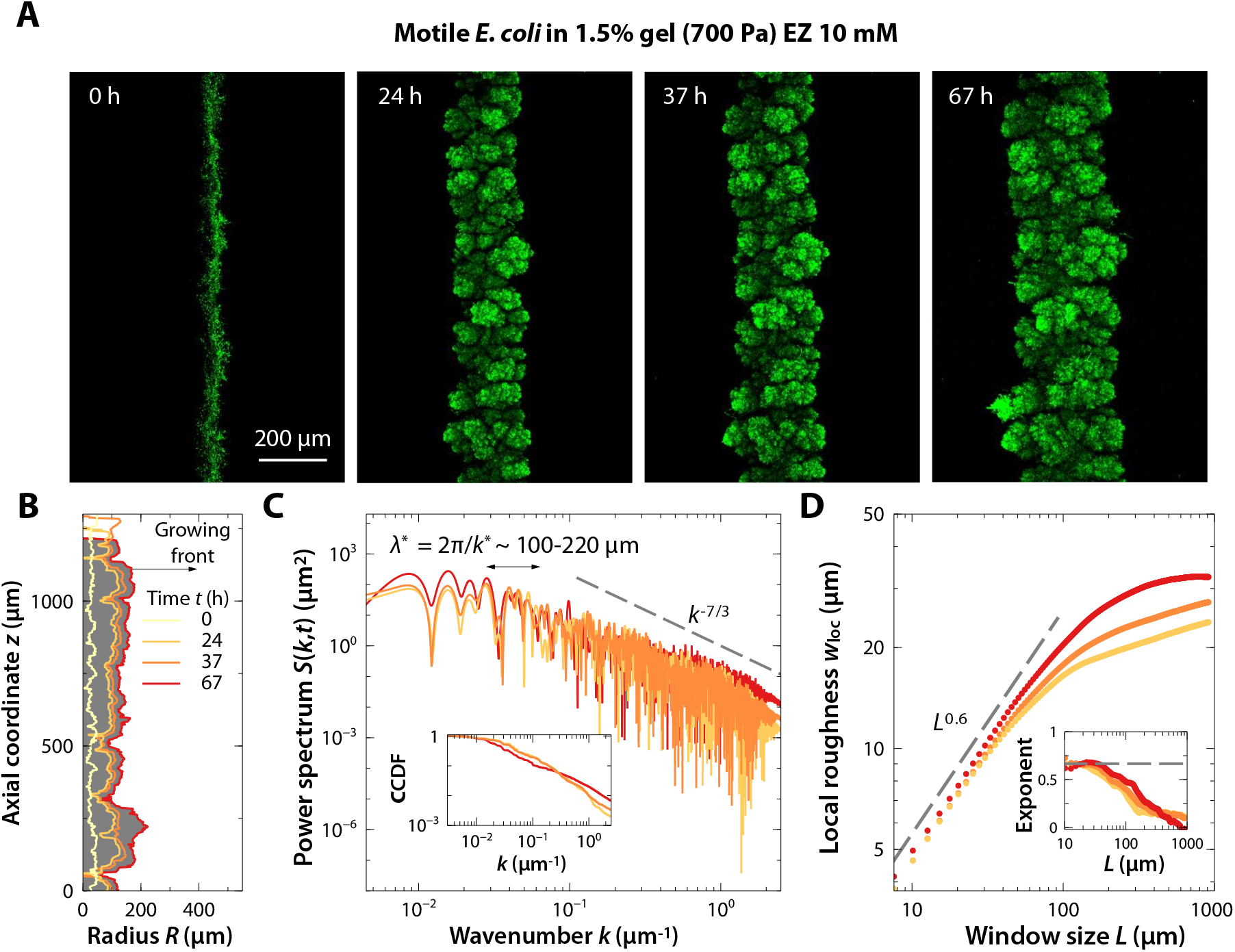
A colony of motile *E. coli* that are immobilized in a tight hydrogel matrix exhibits a roughening instability as it grows outward in 3D. (*A*) Snapshots of the time evolution of a colony of motile *E. coli* displaying a roughening instability as the colony grows in a 3D hydrogel matrix with 1.5% gel. The EZ rich concentration in the medium is 10 mM. The images show bottom-up projections of cellular fluorescence intensity measured using 3D stacks of confocal images taken at different depths (as schematized in Fig. 1*A* of the Main Text). (*B*) Side view of a slice of the colony shown in *A* displaying a section of the leading edge at the same times as in *B*. (*C*) Power spectrum *S*(*k, t*) of the leading edge of the colony in *B* at times *t* = 24, 37, 67 h as a function of the axial wavenumber *k*. The lower and upper limits of the wavenumbers displayed correspond to the size of the domain and the resolution limit, respectively. The power spectra exhibit similar powerlaw decays and characteristic ‘floret’ sizes λ* at the different times. (*Inset*) Cumulative power spectrum as a function of the wavenumber at times *t* = 24, 37, 67 h. (*D*) Local roughness *w*_loc_ as a function of the window size *L* at times *t* = 24, 37, 67 h. Inset shows the corresponding local roughness exponent for each dataset as a function of *L*. We observe similar roughening behavior at the different times.

**FIG. S2.**
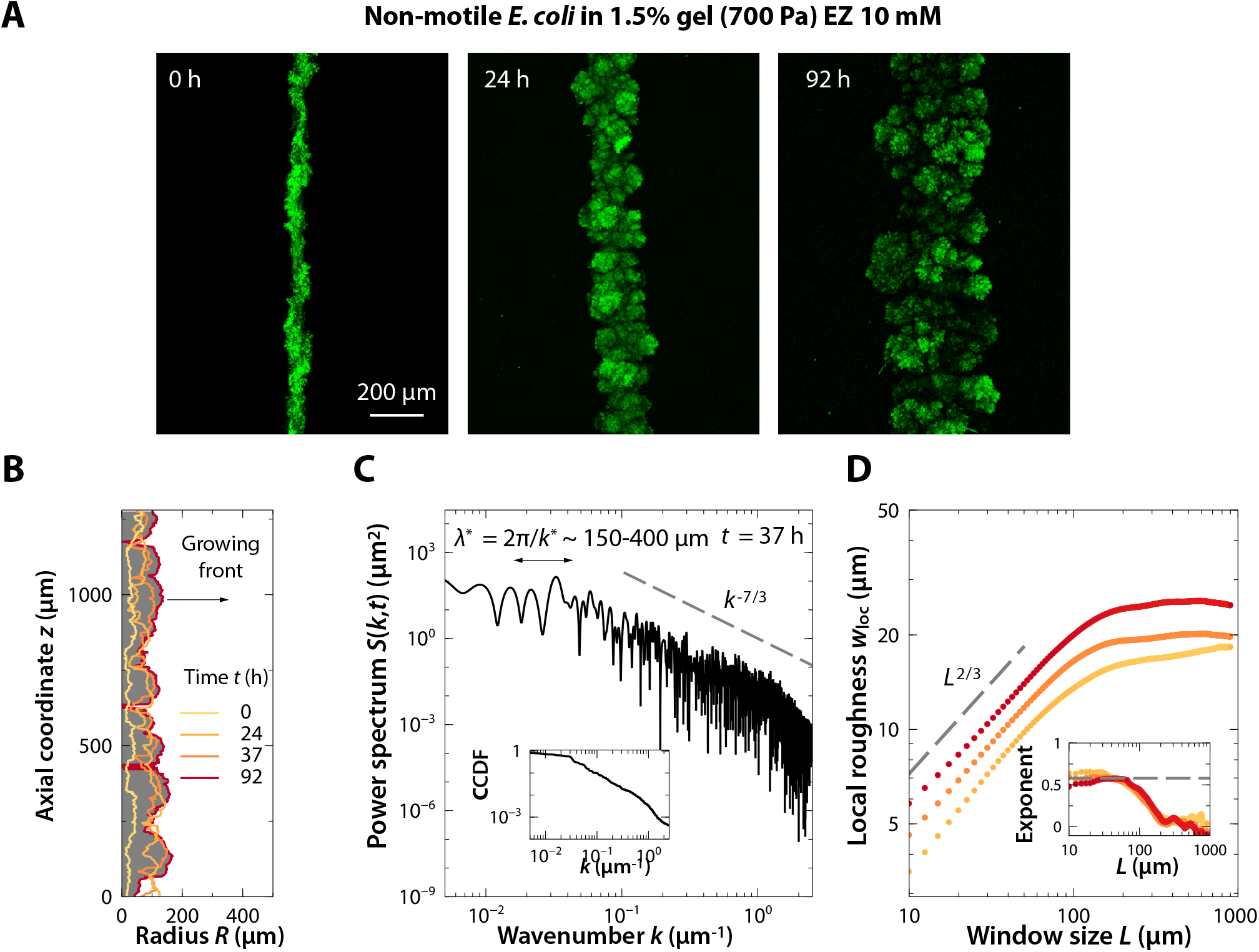
A colony of non-motile *E. coli* in a tight hydrogel matrix (same as Experiment #1) exhibits the same roughening instability as it grows outward in 3D. The comparison with Experiment #1 indicates that cellular motility does not appreciably influence colony expansion and roughening. (*A*) Snapshots of the time evolution of a colony of non-motile *E. coli* displaying a roughening instability as the colony grows in a 3D hydrogel matrix with 1.5% gel. The EZ rich concentration in the medium is 10 mM. The images show bottom-up projections of cellular fluorescence intensity measured using 3D stacks of confocal images taken at different depths (as schematized in Fig. 1*A* of the Main Text). (*B*) Side view of a slice of the colony shown in *A* displaying a section of the leading edge at the same times as in *B*. (*C*) Power spectrum *S*(*k, t*) of the leading edge of the colony in *B* at time *t* = 37 h as a function of the axial wavenumber *k*. The lower and upper limits of the wavenumbers displayed correspond to the size of the domain and the resolution limit, respectively. (*Inset*) Cumulative power spectrum as a function of the wavenumber at *t* = 37 h. (*D*) Local roughness *w*_loc_ as a function of the window size *L* at times *t* = 24, 37, and 92 h. Inset shows the corresponding local roughness exponent for each dataset as a function of *L*. We observe similar roughening behavior at the different times.

**FIG. S3.**
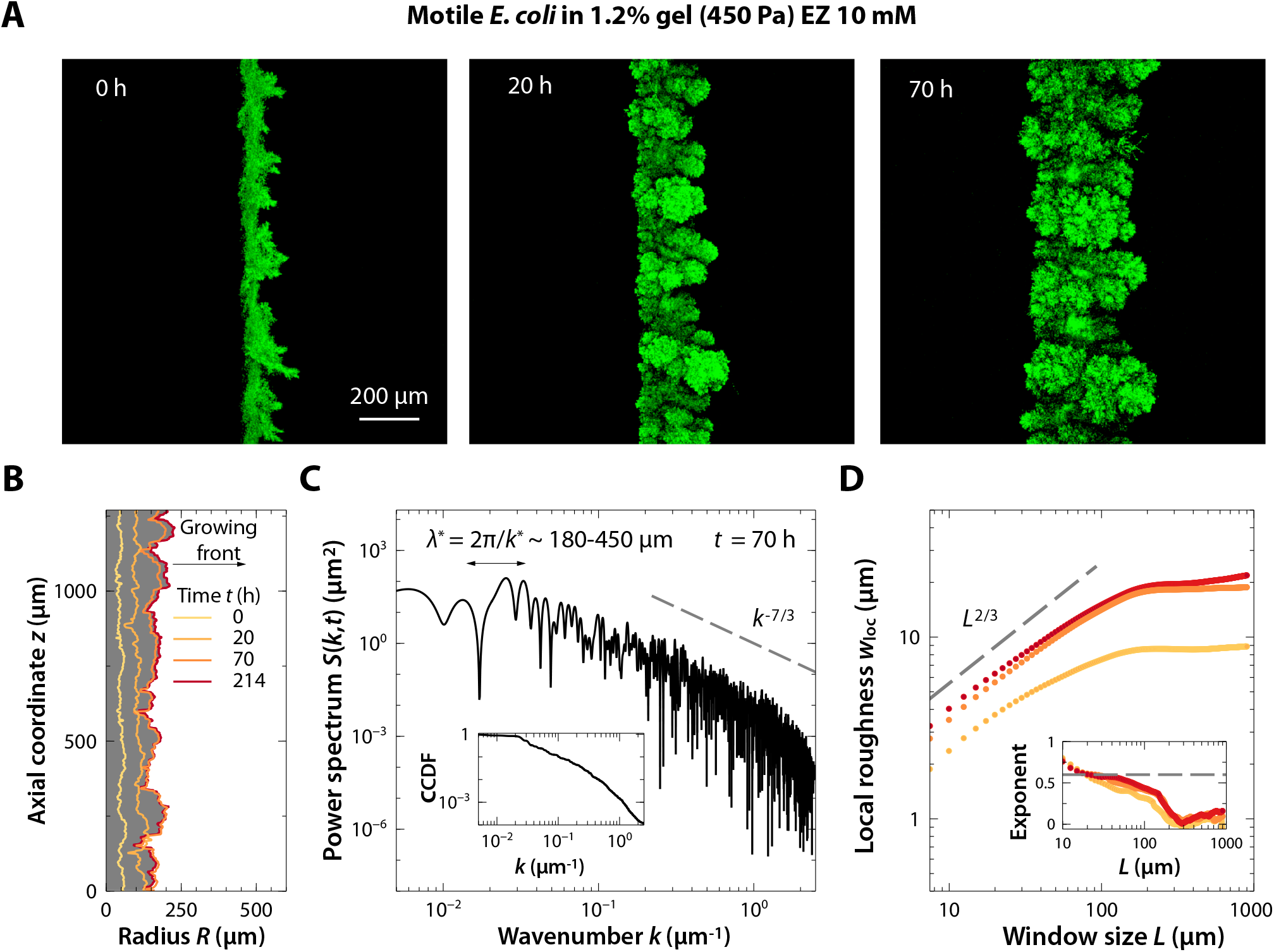
A colony of motile *E. coli* that are immobilized in a tight hydrogel matrix (but looser and softer than that in Experiment #1) exhibits the same roughening instability as it grows outward in 3D. The comparison with Experiment #1 indicates that the matrix pore size and mechanical properties do not appreciably influence colony expansion and roughening. (*A*) Snapshots of the time evolution of a colony of motile *E. coli* displaying a roughening instability as the colony grows in a 3D hydrogel matrix with 1.2% gel. The EZ rich concentration in the medium is 10 mM. The images show bottom-up projections of cellular fluorescence intensity measured using 3D stacks of confocal images taken at different depths (as schematized in Fig. 1*A* of the Main Text). (*B*) Side view of a slice of the colony shown in *A* displaying a section of the leading edge at the same times as in *B*. (*C*) Power spectrum *S*(*k, t*) of the leading edge of the colony in *B* at time *t* = 70 h as a function of the axial wavenumber *k*. The lower and upper limits of the wavenumbers displayed correspond to the size of the domain and the resolution limit, respectively. (*Inset*) Cumulative power spectrum as a function of the wavenumber at *t* = 70 h. (*D*) Local roughness *w*_loc_ as a function of the window size *L* at times *t* = 20, 70, and 214 h. Inset shows the corresponding local roughness exponent for each dataset as a function of *L*. We observe similar roughening behavior at the different times.

**FIG. S4.**
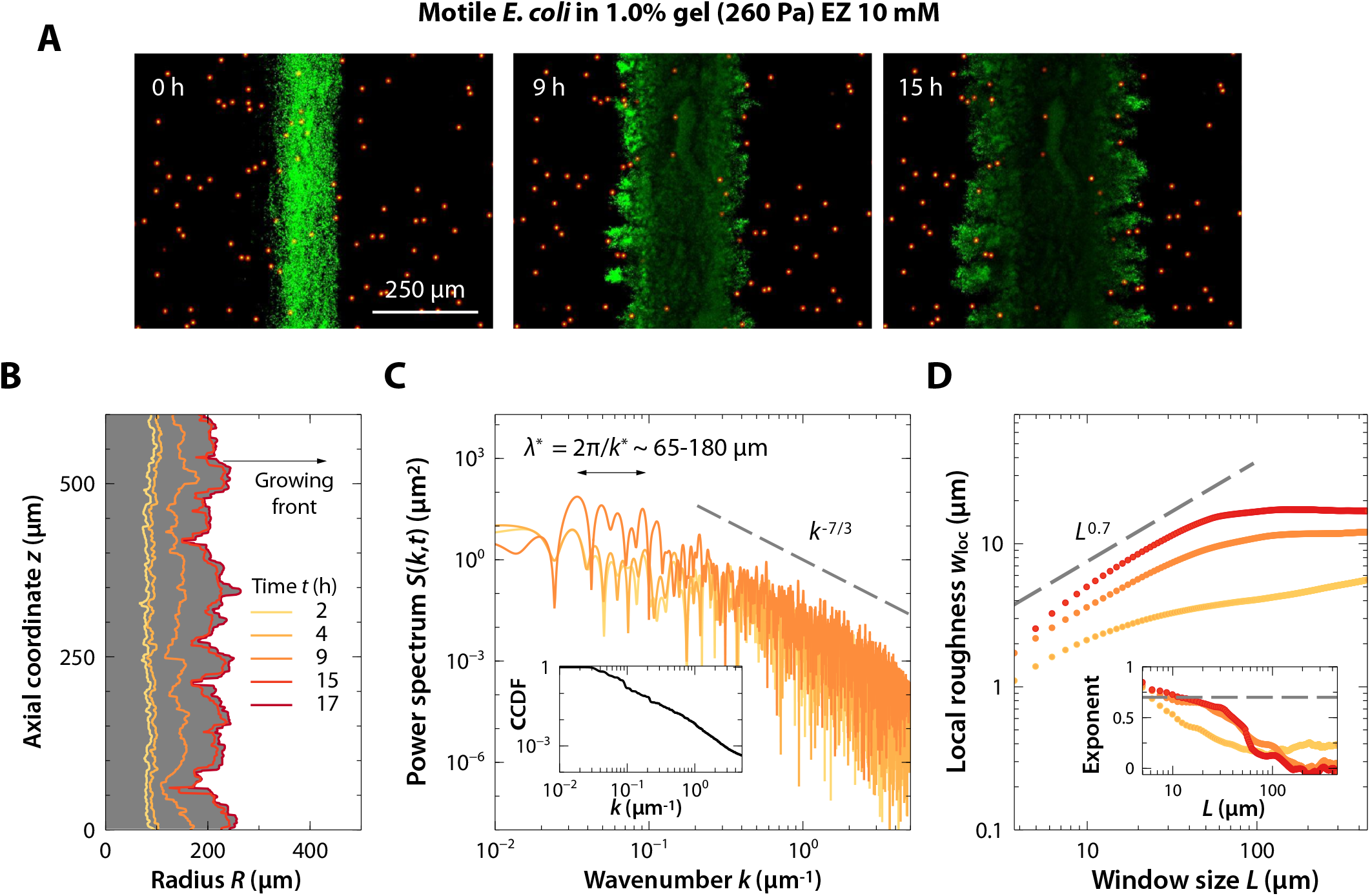
A colony of motile *E. coli* that are immobilized in a tight hydrogel matrix (but looser and softer than that in Experiment #1) exhibits the same roughening instability as it grows outward in 3D. The comparison with Experiment #1 indicates that the matrix pore size and mechanical properties do not appreciably influence colony expansion and roughening. (*A*) Snapshots of the time evolution of a colony of motile *E. coli* displaying a roughening instability as the colony grows in a 3D hydrogel matrix with 1.0% gel. The EZ rich concentration in the medium is 10 mM. The images show bottom-up projections of cellular fluorescence intensity measured using 3D stacks of confocal images taken at different depths (as schematized in Fig. 1*A* of the Main Text). The orange particles show immobilized tracers embedded within the hydrogel matrix as fiducial markers, showing that the matrix locally yields at the surface of the growing colony. (*B*) Side view of a slice of the colony shown in *A* displaying a section of the leading edge at the same times as in *B*. (*C*) Power spectrum *S*(*k, t*) of the leading edge of the colony in *B* at times *t* = 2, 4, and 9 h as functions of the axial wavenumber *k*. The lower and upper limits of the wavenumbers displayed correspond to the size of the domain and the resolution limit, respectively. The power spectra exhibit similar power-law decays and characteristic ‘floret’ sizes λ* at the different times. (*Inset*) Complementary CDF of the power spectrum as a function of the wavenumber at *t* = 9 h. (*D*) Local roughness *w*_loc_ as a function of the window size *L* at times *t* = 4, 9, and 15 h. Inset shows the corresponding local roughness exponent for each dataset as a function of *L*. We observe similar roughening behavior at the different times.

**FIG. S5.**
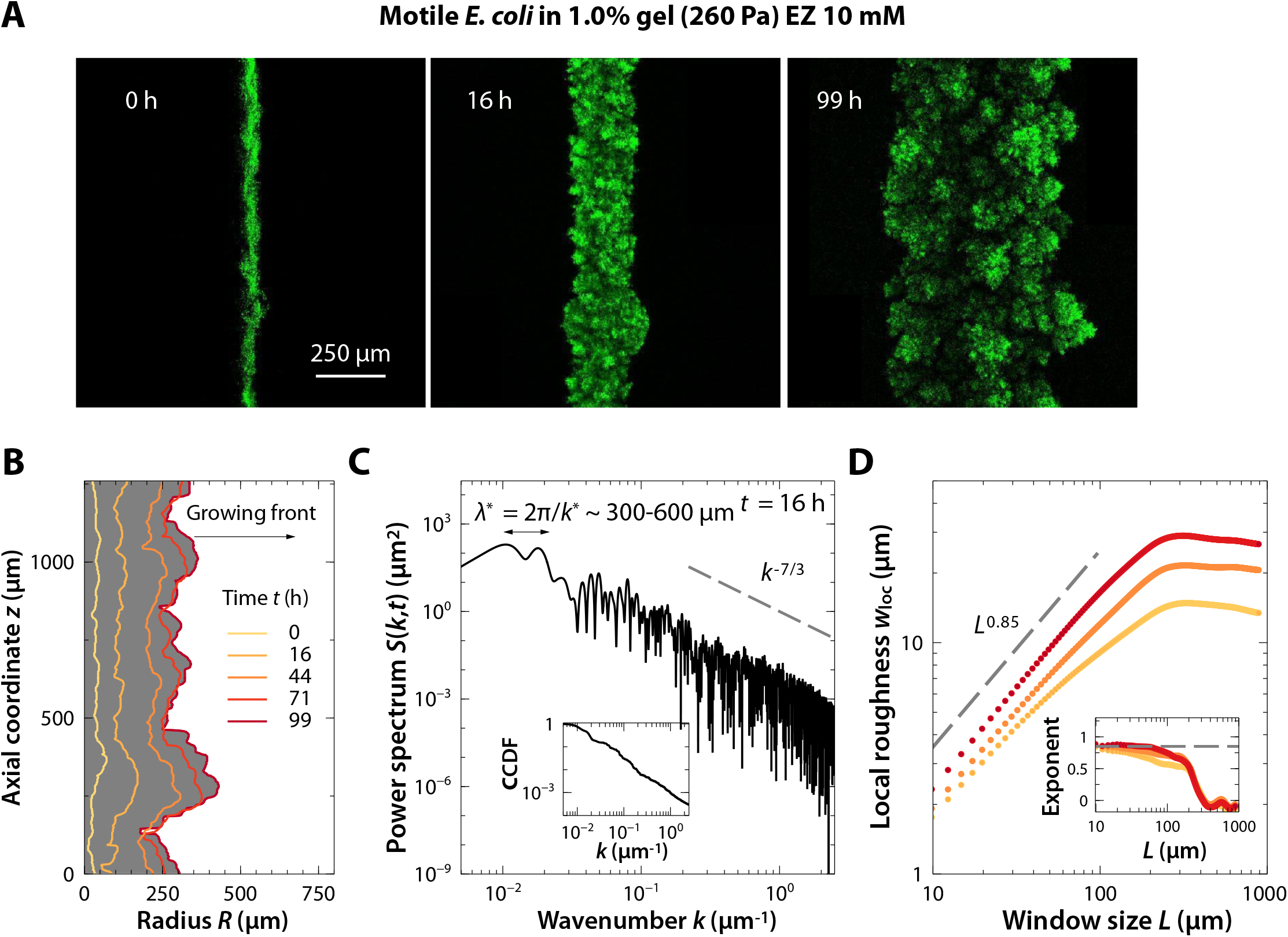
A colony of motile *E. coli* that are immobilized in a tight hydrogel matrix (but looser and softer than that in Experiment #1) exhibits the same roughening instability as it grows outward in 3D. The comparison with Experiment #1 indicates that the matrix pore size and mechanical properties do not appreciably influence colony expansion and roughening. (*A*) Snapshots of the time evolution of a colony of motile *E. coli* displaying a roughening instability as the colony grows in a 3D hydrogel matrix with 1.0% gel. The EZ rich concentration in the medium is 10 mM. The images show bottom-up projections of cellular fluorescence intensity measured using 3D stacks of confocal images taken at different depths (as schematized in Fig. 1*A* of the Main Text). (*B*) Side view of a slice of the colony shown in *A* displaying a section of the leading edge at the same times as in *B*. (*C*) Power spectrum *S*(*k, t*) of the leading edge of the colony in *B* at time *t* = 16 h as a function of the axial wavenumber *k*. The lower and upper limits of the wavenumbers displayed correspond to the size of the domain and the resolution limit, respectively. (*Inset*) Complementary CDF of the power spectrum as a function of the wavenumber at *t* = 16 h. (*D*) Local roughness *w*_loc_ as a function of the window size *L* at times *t* = 16, 44, and 71 h. Inset shows the corresponding local roughness exponent for each dataset as a function of *L*. We observe similar roughening behavior at the different times.

**FIG. S6.**
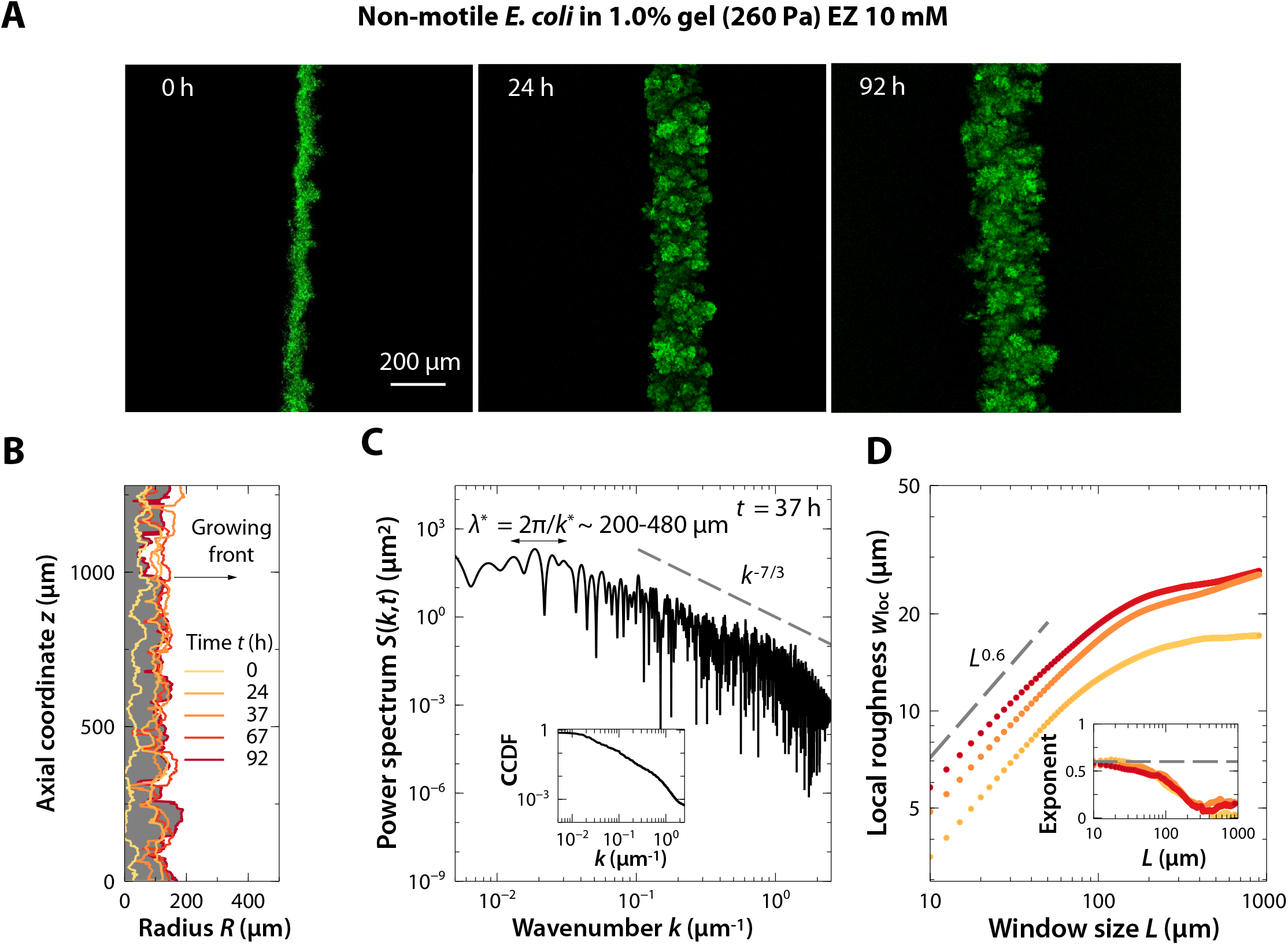
A colony of non-motile *E. coli* that are immobilized in a tight hydrogel matrix (but looser and softer than that in Experiment #2) exhibits the same roughening instability as it grows outward in 3D. The comparison with Experiment #2 indicates that the matrix pore size and mechanical properties do not appreciably influence colony expansion and roughening. (*A*) Snapshots of the time evolution of a colony of non-motile *E. coli* displaying a roughening instability as the colony grows in a 3D hydrogel matrix with 1.0% gel. The EZ rich concentration in the medium is 10 mM. The images show bottom-up projections of cellular fluorescence intensity measured using 3D stacks of confocal images taken at different depths (as schematized in Fig. 1*A* of the Main Text). (*B*) Side view of a slice of the colony shown in *A* displaying a section of the leading edge at the same times as in *B*. (*C*) Power spectrum *S*(*k, t*) of the leading edge of the colony in *B* at time *t* = 37 h as a function of the axial wavenumber *k*. The lower and upper limits of the wavenumbers displayed correspond to the size of the domain and the resolution limit, respectively. (*Inset*) Cumulative power spectrum as a function of the wavenumber at *t* = 37 h. (*D*) Local roughness *w*_loc_ as a function of the window size *L* at times *t* = 24, 37, and 92 h. Inset shows the corresponding local roughness exponent for each dataset as a function of *L*. We observe similar roughening behavior at the different times.

**FIG. S7.**
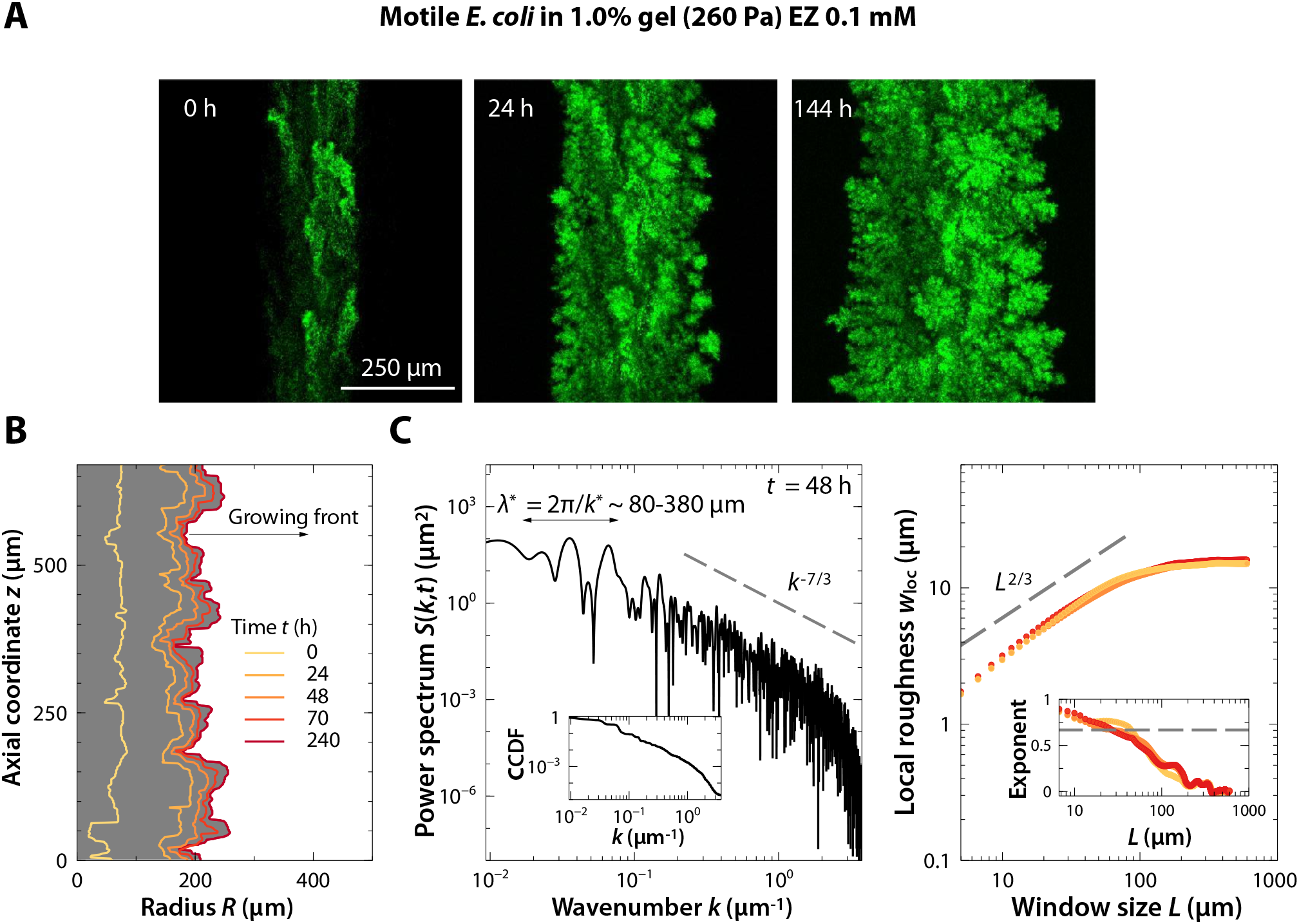
A colony of motile *E. coli* that are immobilized in a tight hydrogel matrix (but nutrient-depleted, compared to the nutrient-replete matrix of Experiment #5) exhibits the same roughening instability as it grows outward in 3D. The comparison with Experiment #5 indicates that the far-field nutrient concentration does not appreciably influence colony expansion and roughening in this regime of growth. (*A*) Snapshots of the time evolution of a colony of motile *E. coli* displaying a roughening instability as the colony grows in a 3D hydrogel matrix with 1.0% gel. The EZ rich concentration in the medium is 0.1 mM. The images show bottom-up projections of cellular fluorescence intensity measured using 3D stacks of confocal images taken at different depths (as schematized in Fig. 1*A* of the Main Text). (*B*) Side view of a slice of the colony shown in *A* displaying a section of the leading edge at the same times as in *B*. (*C*) Power spectrum *S*(*k, t*) of the leading edge of the colony in *B* at time *t* = 48 h as a function of the axial wavenumber *k*. The lower and upper limits of the wavenumbers displayed correspond to the size of the domain and the resolution limit, respectively. (*Inset*) Complementary CDF of the power spectrum as a function of the wavenumber at *t* = 48 h. (*D*) Local roughness *w*_loc_ as a function of the window size *L* at times *t* = 24, 48, and 71 h. Inset shows the corresponding local roughness exponent for each dataset as a function of *L*. We observe similar roughening behavior at the different times.

**FIG. S8.**
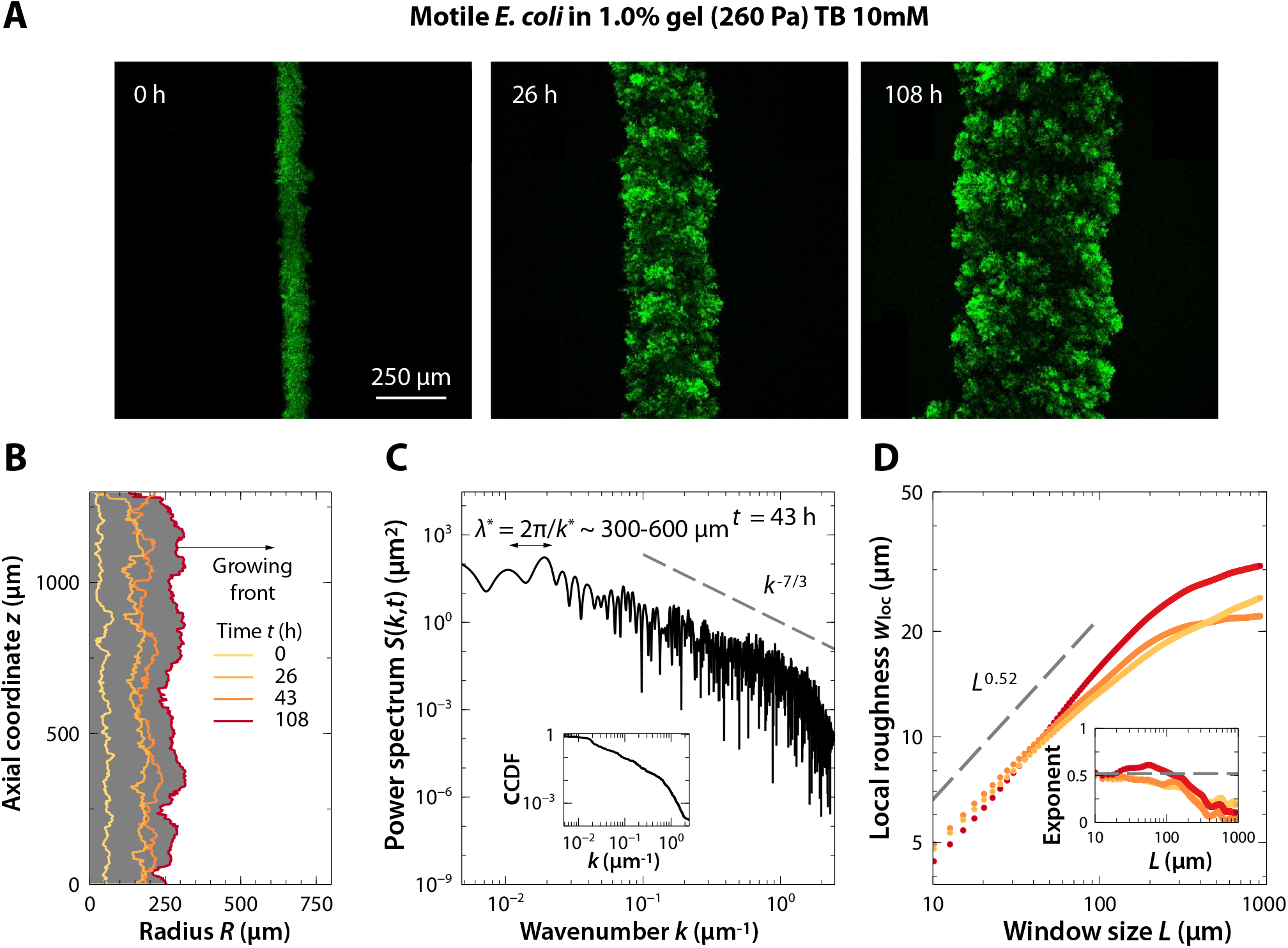
A colony of motile *E. coli* that are immobilized in a tight hydrogel matrix (but formulated with a different liquid cell culture medium compared to the matrix of Experiment #5) exhibits the same roughening instability as it grows outward in 3D. The comparison with Experiment #5 indicates that the exact identity of the liquid cell culture medium does not appreciably influence colony expansion and roughening in this regime of growth. (*A*) Snapshots of the time evolution of a colony of motile *E. coli* displaying a roughening instability as the colony grows in a 3D hydrogel matrix with 1.0% gel. The TB concentration in the medium is 10 mM. The images show bottom-up projections of cellular fluorescence intensity measured using 3D stacks of confocal images taken at different depths (as schematized in Fig. 1*A* of the Main Text). (*B*) Side view of a slice of the colony shown in *A* displaying a section of the leading edge at the same times as in *B*. (*C*) Power spectrum *S*(*k, t*) of the leading edge of the colony in *B* at time *t* = 43 h as a function of the axial wavenumber *k*. The lower and upper limits of the wavenumbers displayed correspond to the size of the domain and the resolution limit, respectively. (*Inset*) Cumulative power spectrum as a function of the wavenumber at *t* = 43 h. (*D*) Local roughness *w*_loc_ as a function of the window size L at times *t* = 26, 43, and 108 h. Inset shows the corresponding local roughness exponent for each dataset as a function of *L*. We observe similar roughening behavior at the different times.

**FIG. S9.**
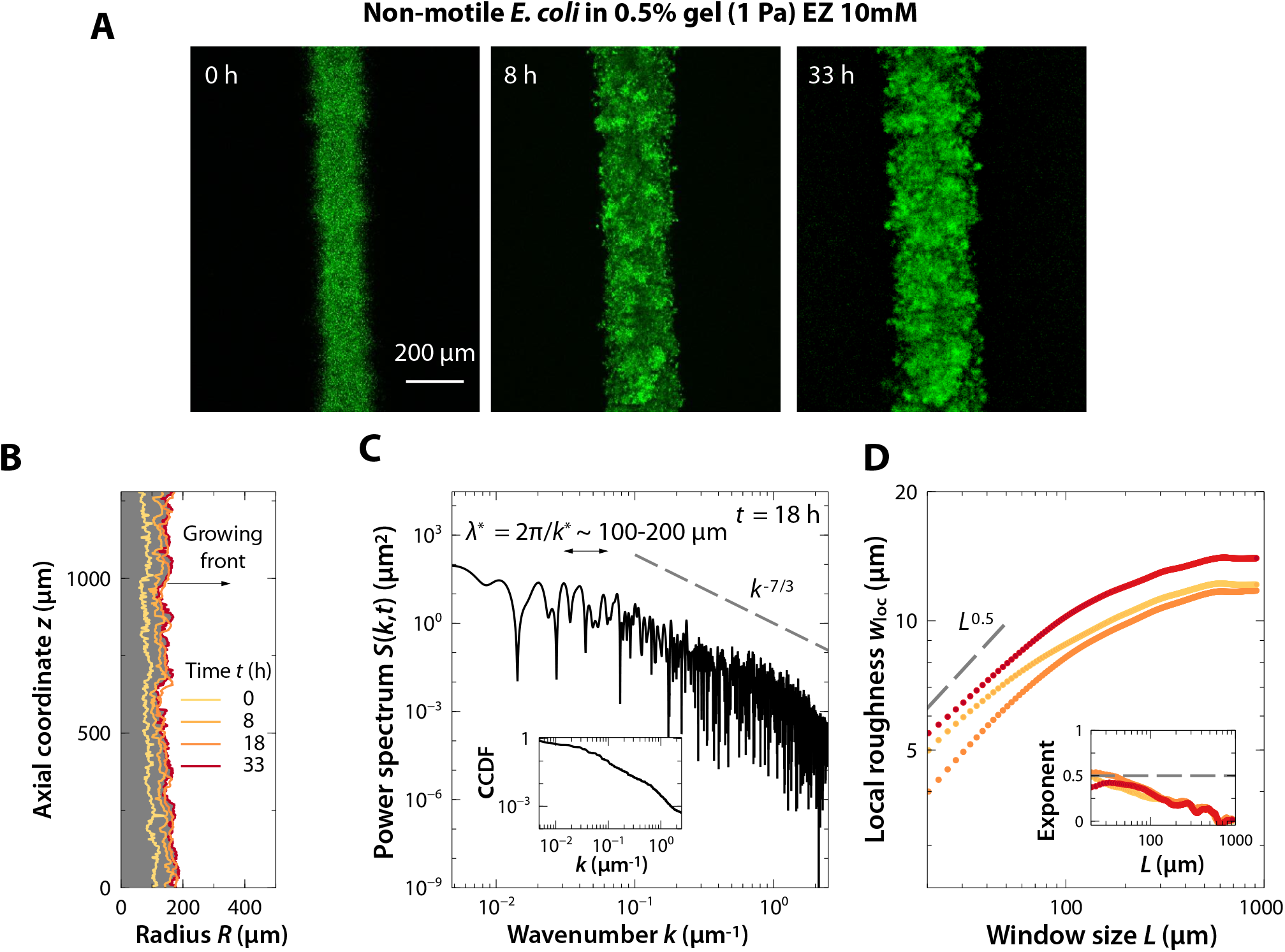
A colony of non-motile *E. coli* that are immobilized in a loose hydrogel matrix (much looser and softer than the matrix of Experiment #2) exhibits the same roughening instability as it grows outward in 3D. The comparison with Experiment #2 indicates that the matrix pore size and mechanical properties do not appreciably influence colony expansion and roughening. (*A*) Snapshots of the time evolution of a colony of non-motile *E. coli* displaying a roughening instability as the colony grows in a 3D hydrogel matrix with 0.5% gel. The EZ rich concentration in the medium is 10 mM. The images show bottom-up projections of cellular fluorescence intensity measured using 3D stacks of confocal images taken at different depths (as schematized in Fig. 1*A* of the Main Text). (*B*) Side view of a slice of the colony shown in *A* displaying a section of the leading edge at the same times as in *B*. (*C*) Power spectrum *S*(*k, t*) of the leading edge of the colony in *B* at time *t* = 18 h as a function of the axial wavenumber *k*. The lower and upper limits of the wavenumbers displayed correspond to the size of the domain and the resolution limit, respectively. (*Inset*) Complementary CDF of the power spectrum as a function of the wavenumber at *t* = 18 h. (*D*) Local roughness w_loc_ as a function of the window size *L* at times *t* = 8, 18, and 33 h. Inset shows the corresponding local roughness exponent for each dataset as a function of *L*. We observe similar roughening behavior at the different times.

**FIG. S10.**
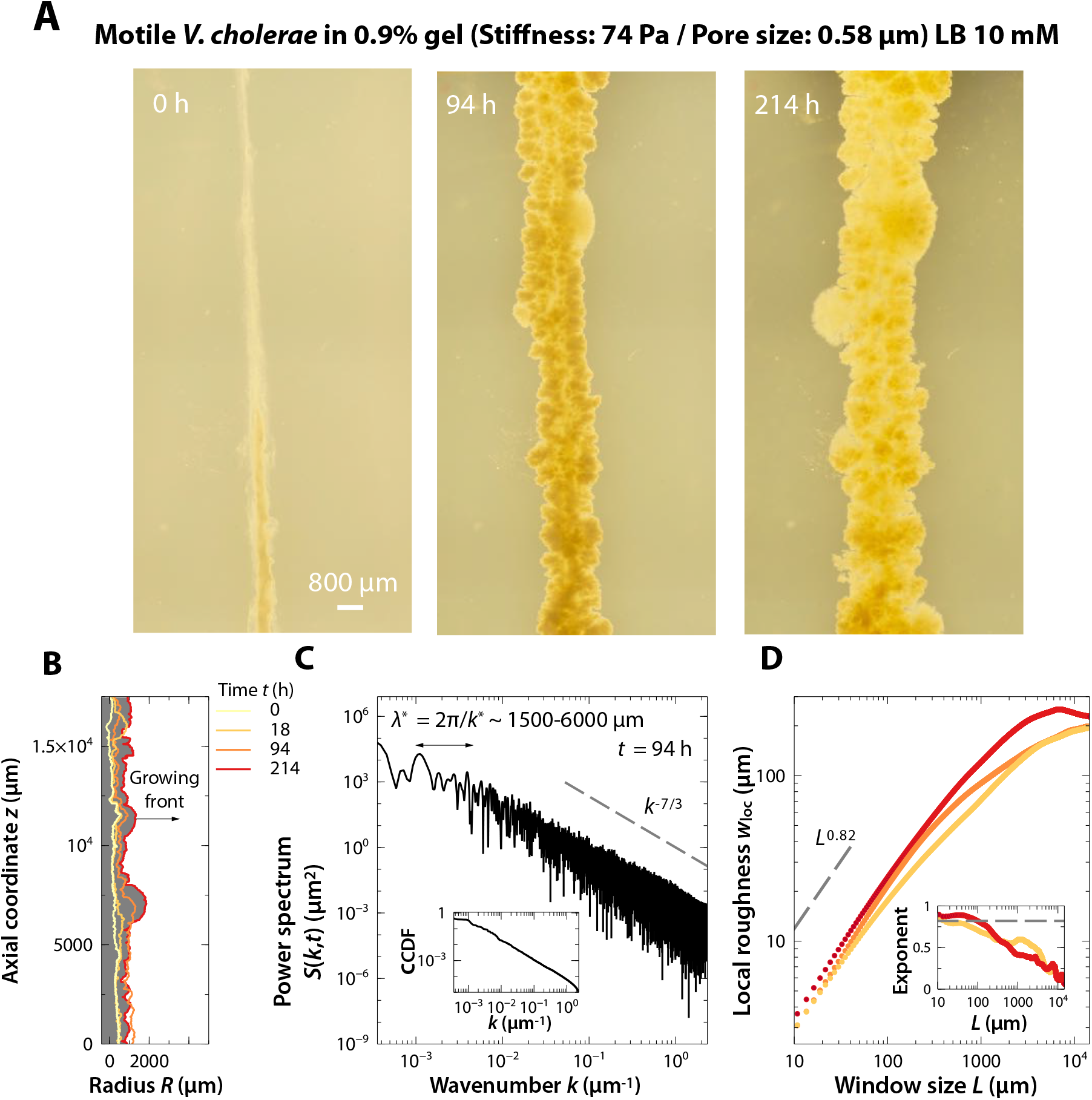
A colony of motile *V. cholerae* that are immobilized in a tight hydrogel matrix exhibits the same roughening instability as it grows outward in 3D. (*A*) Snapshots of the time evolution of a colony of motile *V. cholerae* displaying a roughening instability as the colony grows in a 3D hydrogel matrix with 0.9% gel. The LB medium concentration is 10 mM. The images show side-view photographs taken for a colony that was initially 3D-printed as a vertically-oriented cylinder (t = 0) in a transparent-walled container. (*B*) Side view of a slice of the colony shown in *A* displaying a section of the leading edge at the same times as in *B*. (*C*) Power spectrum *S*(*k, t*) of the leading edge of the colony in *B* at time *t* = 94 h as a function of the axial wavenumber *k*. The lower and upper limits of the wavenumbers displayed correspond to the size of the domain and the resolution limit, respectively. (*Inset*) Cumulative power spectrum as a function of the wavenumber at *t* = 94 h. (*D*) Local roughness *w*_loc_ as a function of the window size *L* at times *t* = 18, 94, and 214 h. Inset shows the corresponding local roughness exponent for each dataset as a function of *L*. We observe similar roughening behavior at the different times.

**FIG. S11.**
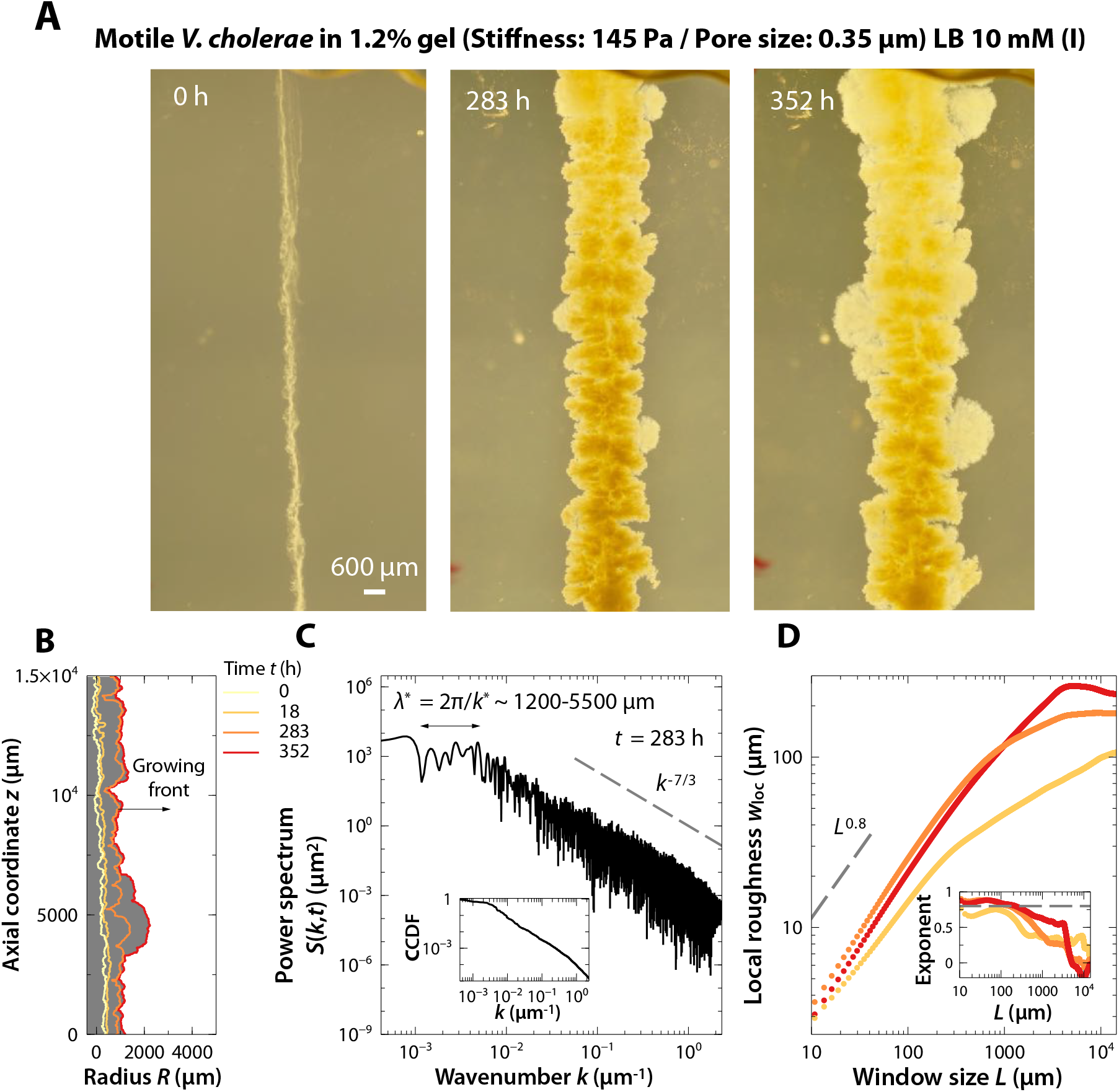
A colony of motile *V. cholerae* that are immobilized in a tight hydrogel matrix (tighter and stiffer than that in Experiment #10) exhibits the same roughening instability as it grows outward in 3D. The comparison with Experiment #10 indicates that the matrix pore size and mechanical properties do not appreciably influence colony expansion and roughening. (*A*) Snapshots of the time evolution of a colony of motile *V. cholerae* displaying a roughening instability as the colony grows in a 3D hydrogel matrix with 1.2% gel. The LB concentration in the medium is 10 mM. The images show side-view photographs taken for a colony that was initially 3D-printed as a vertically-oriented cylinder (*t* = 0) in a transparent-walled container. (*B*) Side view of a slice of the colony shown in *A* displaying a section of the leading edge at the same times as in *B*. (*C*) Power spectrum *S*(*k, t*) of the leading edge of the colony in *B* at time *t* = 283 h as a function of the axial wavenumber *k*. The lower and upper limits of the wavenumbers displayed correspond to the size of the domain and the resolution limit, respectively. (*Inset*) Cumulative power spectrum as a function of the wavenumber at *t* = 283 h. (*D*) Local roughness *w*_loc_ as a function of the window size *L* at times *t* = 18, 283, and 352 h. Inset shows the corresponding local roughness exponent for each dataset as a function of *L*. We observe similar roughening behavior at the different times.

**FIG. S12.**
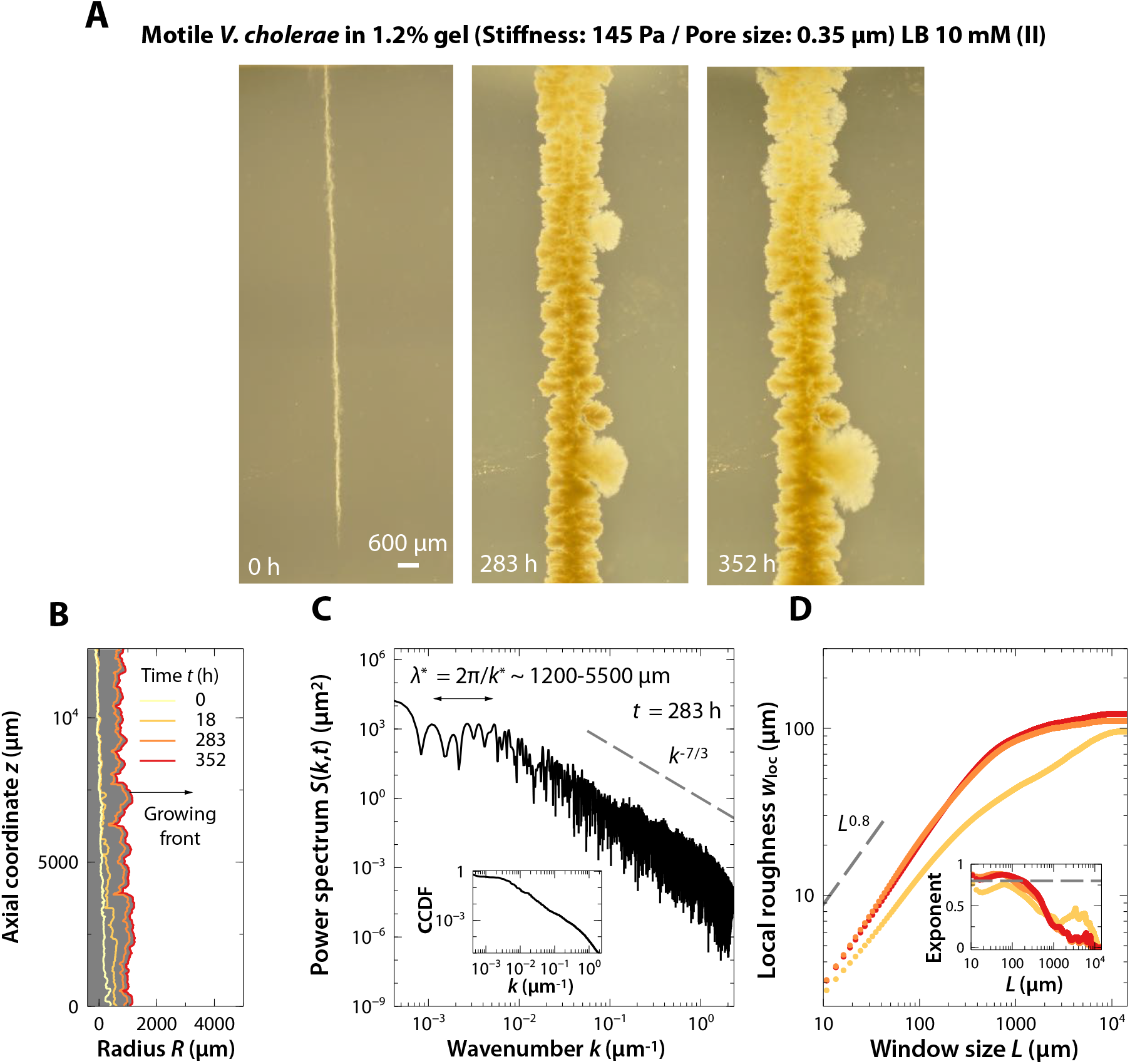
A colony of motile *V. cholerae* that are immobilized in a tight hydrogel matrix (tighter and stiffer than that in Experiment #10) exhibits the same roughening instability as it grows outward in 3D. The comparison with Experiment #10 indicates that the matrix pore size and mechanical properties do not appreciably influence colony expansion and roughening. This experiment is a replicate of Experiment #11, but with a thinner starting colony radius, indicating that the starting radius does not appreciably influence colony expansion and roughening. (*A*) Snapshots of the time evolution of a colony of motile *V. cholerae* displaying a roughening instability as the colony grows in a 3D hydrogel matrix with 1.2% gel. The LB concentration in the medium is 10 mM. The images show side-view photographs taken for a colony that was initially 3D-printed as a vertically-oriented cylinder (t = 0) in a transparent-walled container. (*B*) Side view of a slice of the colony shown in *A* displaying a section of the leading edge at the same times as in *B*. (*C*) Power spectrum *S*(*k, t*) of the leading edge of the colony in *B* at time *t* = 283 h as a function of the axial wavenumber *k*. The lower and upper limits of the wavenumbers displayed correspond to the size of the domain and the resolution limit, respectively. (*Inset*) Cumulative power spectrum as a function of the wavenumber at *t* = 283 h. (*D*) Local roughness *w*_loc_ as a function of the window size *L* at times *t* = 18, 283, and 352 h. Inset shows the corresponding local roughness exponent for each dataset as a function of *L*. We observe similar roughening behavior at the different times.

**FIG. S13.**
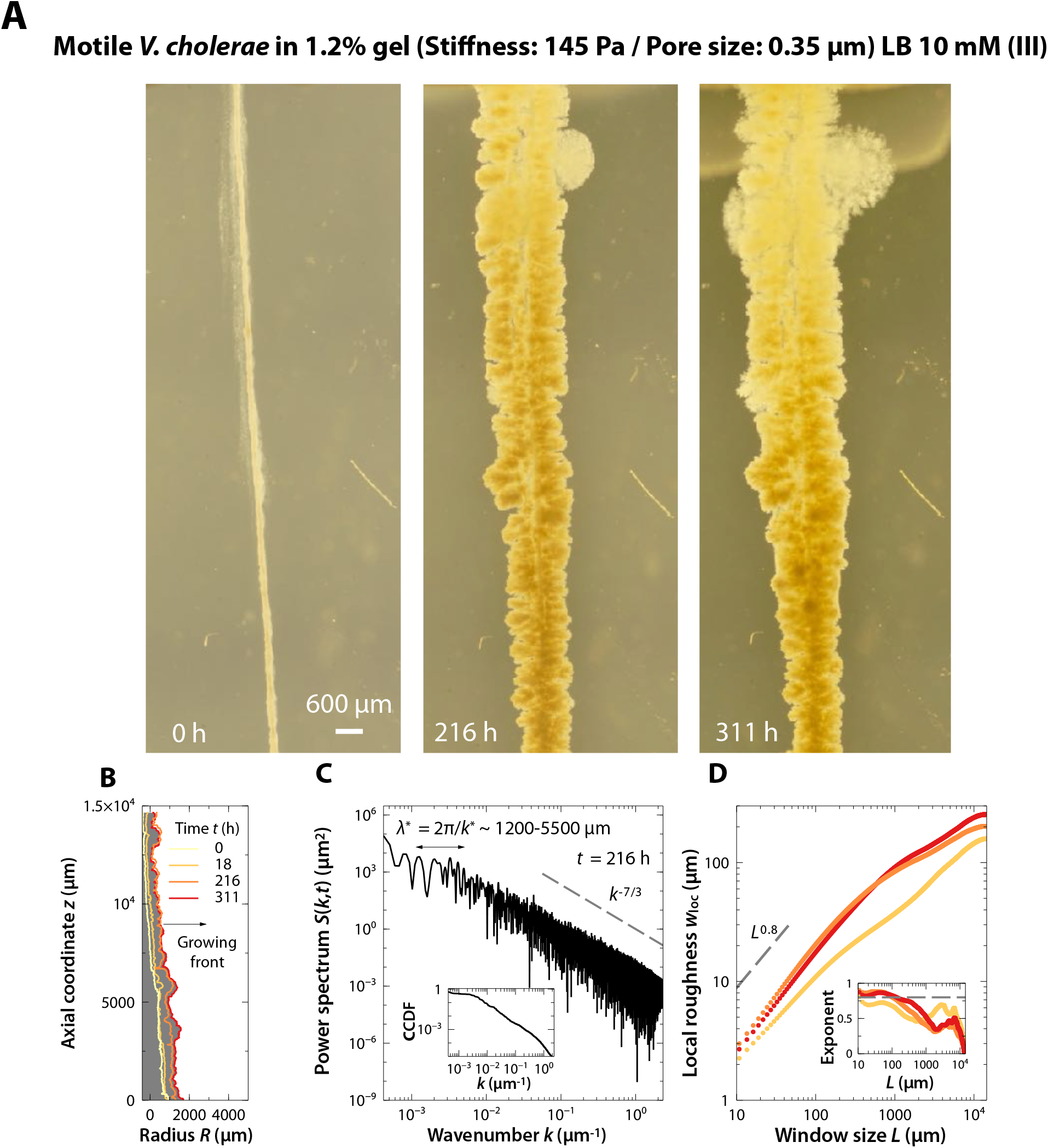
A colony of motile *V. cholerae* that are immobilized in a tight hydrogel matrix (tighter and stiffer than that in Experiment #10) exhibits the same roughening instability as it grows outward in 3D. The comparison with Experiment #10 indicates that the matrix pore size and mechanical properties do not appreciably influence colony expansion and roughening. This experiment is a replicate of Experiments #11 and #12, demonstrating the reproducibility of this phenomenon. (*A*) Snapshots of the time evolution of a colony of motile *V. cholerae* displaying a roughening instability as the colony grows in a 3D hydrogel matrix with 1.5% gel. The LB concentration in the medium is 10 mM. The images show side-view photographs taken for a colony that was initially 3D-printed as a vertically-oriented cylinder (*t* = 0) in a transparent-walled container. (*B*) Side view of a slice of the colony shown in *A* displaying a section of the leading edge at the same times as in *B*. (*C*) Power spectrum *S*(*k, t*) of the leading edge of the colony in *B* at time *t* = 216 h as a function of the axial wavenumber *k*. The lower and upper limits of the wavenumbers displayed correspond to the size of the domain and the resolution limit, respectively. (*Inset*) Cumulative power spectrum as a function of the wavenumber at *t* = 216 h. (*D*) Local roughness *w*_loc_ as a function of the window size *L* at times *t* = 18, 94, and 216 h. Inset shows the corresponding local roughness exponent for each dataset as a function of *L*. We observe similar roughening behavior at the different times.

**FIG. S14.**
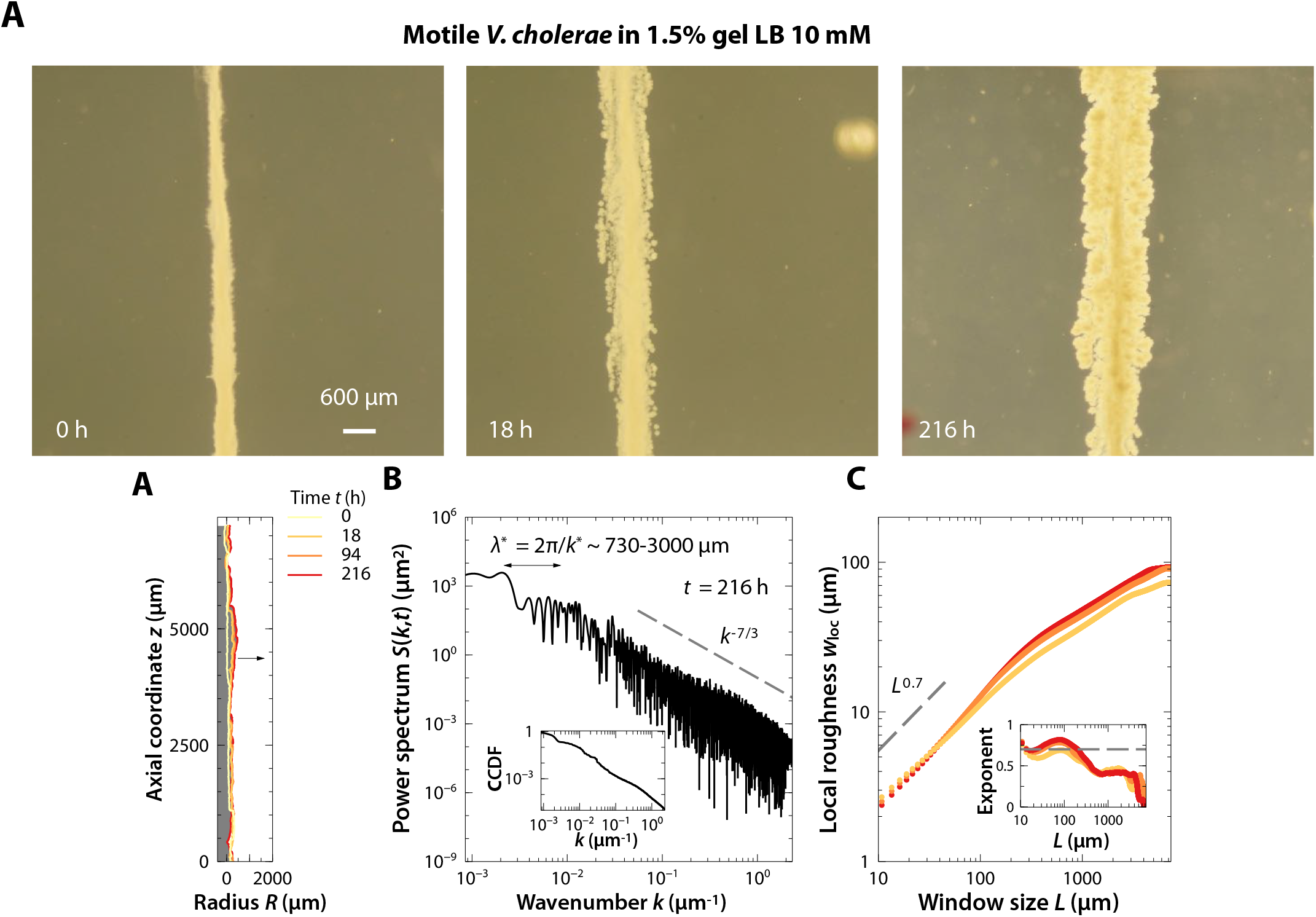
A colony of motile *V. cholerae* that are immobilized in a tight hydrogel matrix (tighter and stiffer than that in Experiment #10) exhibits the same roughening instability as it grows outward in 3D. The comparison with Experiments #10–13 indicates that the matrix pore size and mechanical properties do not appreciably influence colony expansion and roughening. (*A*) Snapshots of the time evolution of a colony of motile *V. cholerae* displaying a roughening instability as the colony grows in a 3D hydrogel matrix with 1.5% gel. The LB concentration in the medium is 10 mM. The images show side-view photographs taken for a colony that was initially 3D-printed as a vertically-oriented cylinder (*t* = 0) in a transparent-walled container. (*B*) Side view of a slice of the colony shown in *A* displaying a section of the leading edge at the same times as in *B*. (*C*) Power spectrum *S*(*k, t*) of the leading edge of the colony in *B* at time *t* = 216 h as a function of the axial wavenumber *k*. The lower and upper limits of the wavenumbers displayed correspond to the size of the domain and the resolution limit, respectively. (*Inset*) Cumulative power spectrum as a function of the wavenumber at *t* = 216 h. (*D*) Local roughness *w*_loc_ as a function of the window size *L* at times *t* = 18, 94, and 216 h. Inset shows the corresponding local roughness exponent for each dataset as a function of *L*. We observe similar roughening behavior at the different times.

**FIG. S15.**
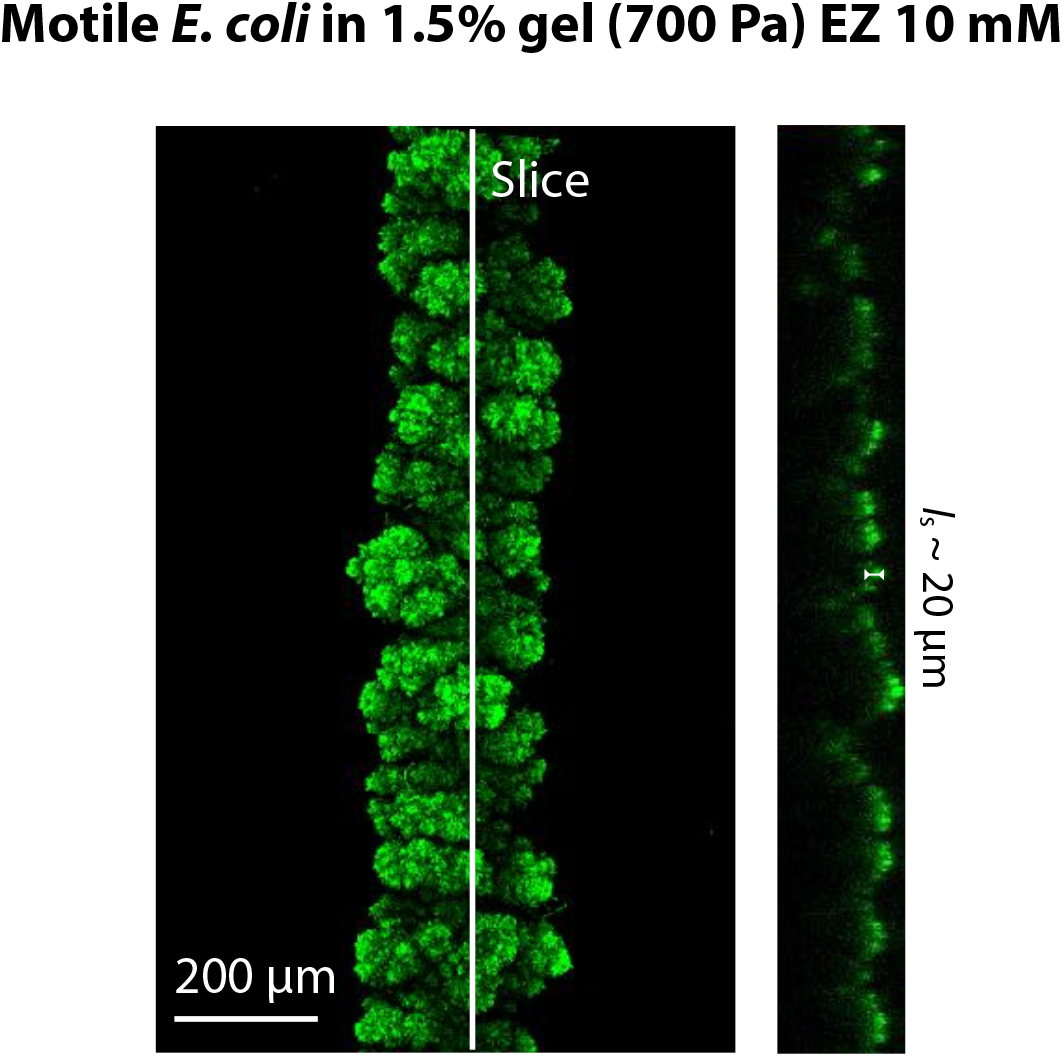
Only a thin layer of cells localized at the surface of a growing 3D colony is metabolically active. (*A*) Left: Snapshot of motile *E. coli* colony growing in 3D corresponding to Fig. 1 of the main text. Right: Fluorescence intensity along the slice indicated in the left panel, which provides a proxy for cellular activity.

**FIG. S16.**
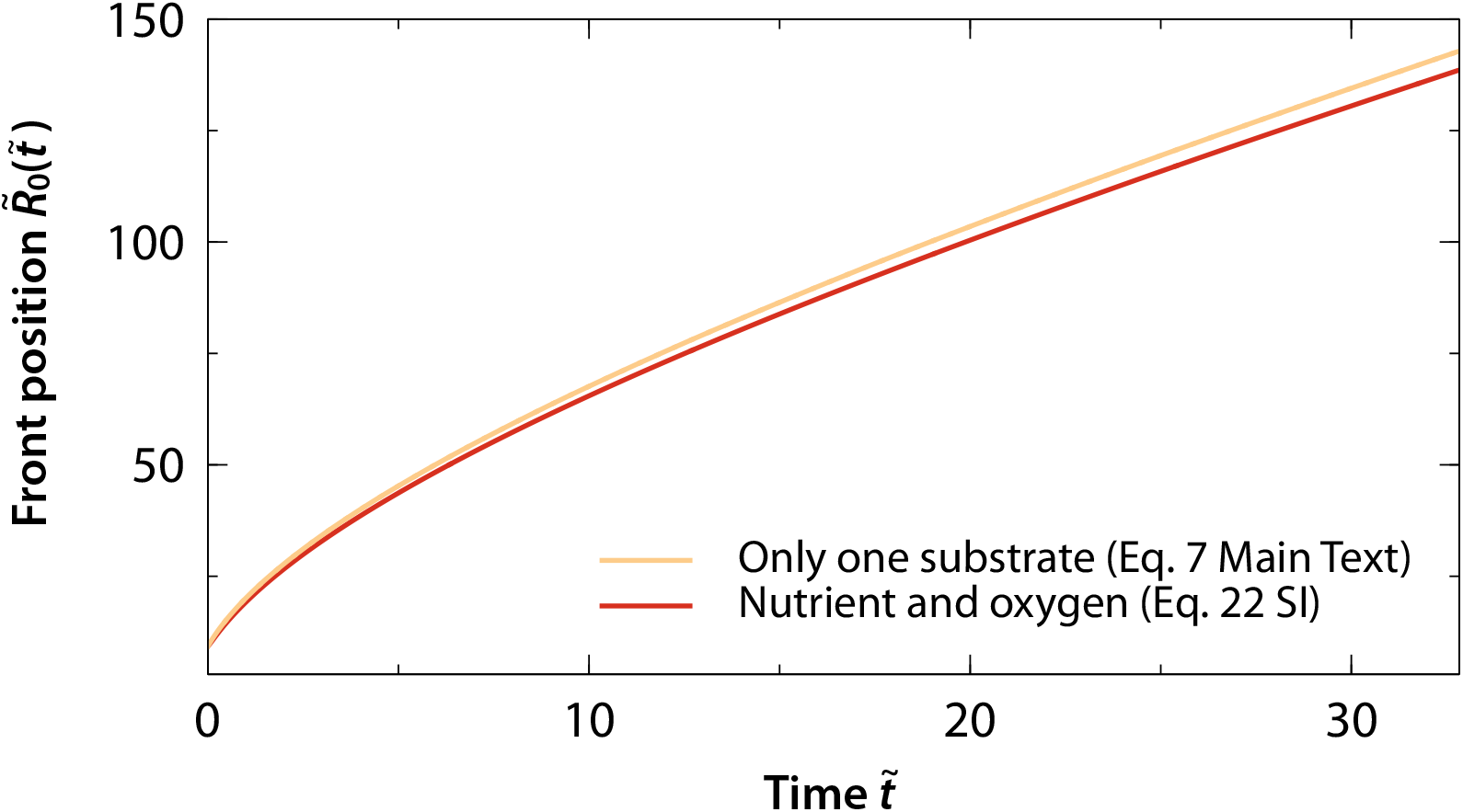
Model predictions are similar when considering either oxygen as the growth-limiting substrate, or both oxygen and nutrient as growth-limiting substrates. Evolution of the advancing cylindrical front 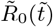 as a function of time, considering either only oxygen as the limiting substrate for growth, i.e. Eq. (7) in the Main Text, or both nutrient and oxygen as substrates required for growth, i.e. Eq. (S22).

**FIG. S17.**
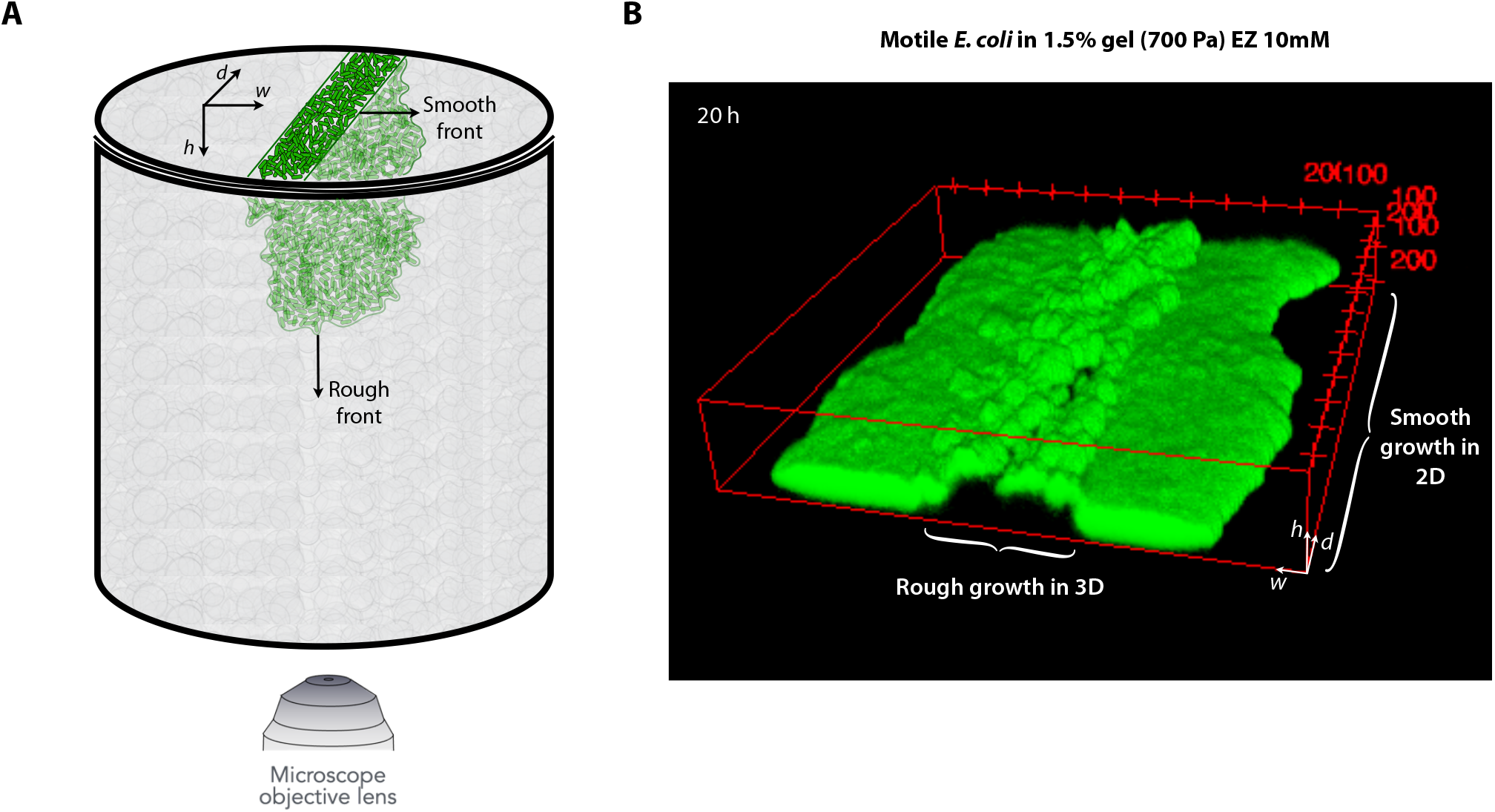
A colony of motile *E. coli* initially located in a thin strip on the surface of a tight hydrogel matrix (identical to that in Experiment #1) shows smooth transverse growth atop the 2D hydrogel surface, but rough growth when it penetrates into the 3D matrix. (*A*) Schematic of the experiment. (*B*) Confocal 3D reconstruction of the colony after *t* = 20 h of growth; note that the vertical orientation is flipped from *A*.

**FIG. S18.**
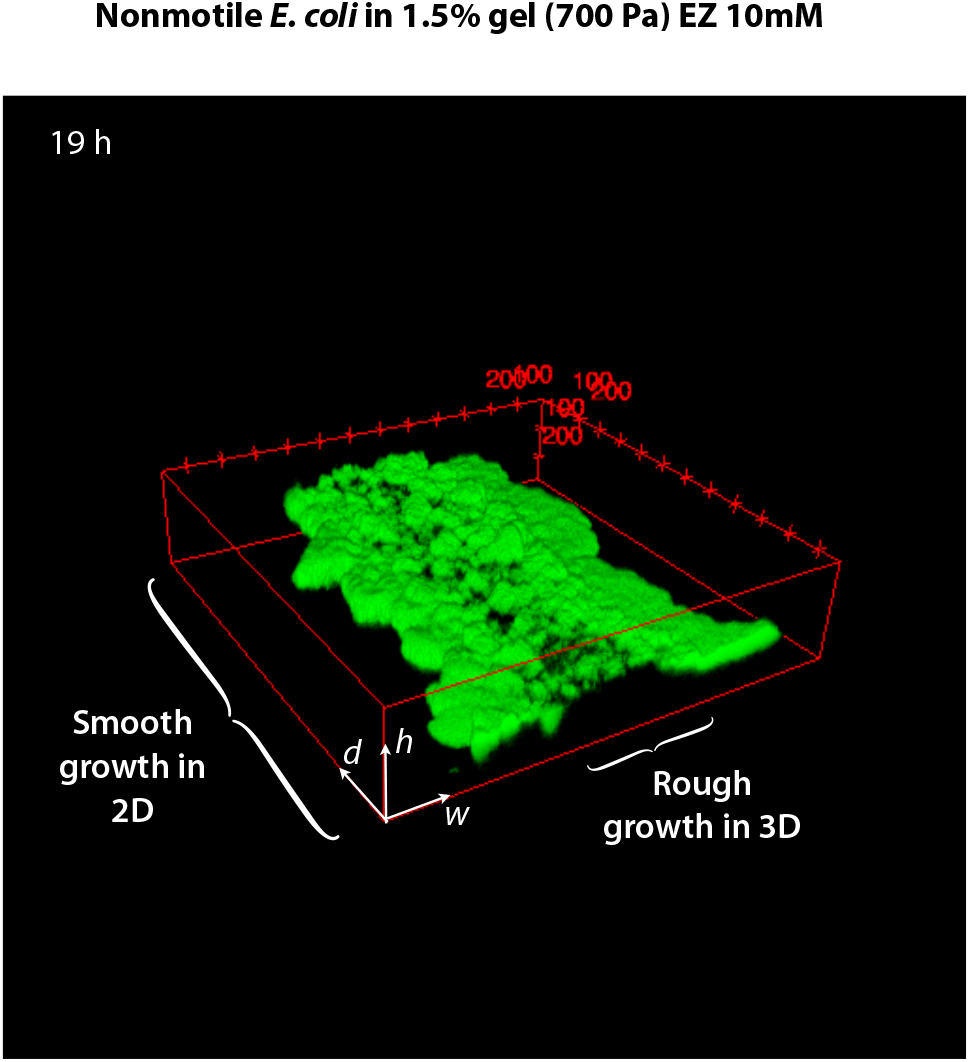
A colony of non-motile *E. coli* initially located in a thin strip on the surface of a tight hydrogel matrix (identical to that in Experiment #1) shows smooth transverse growth atop the 2D hydrogel surface, but rough growth when it penetrates into the 3D matrix. Confocal 3D reconstruction of the colony after *t* = 20 h of growth.

**FIG. S19.**
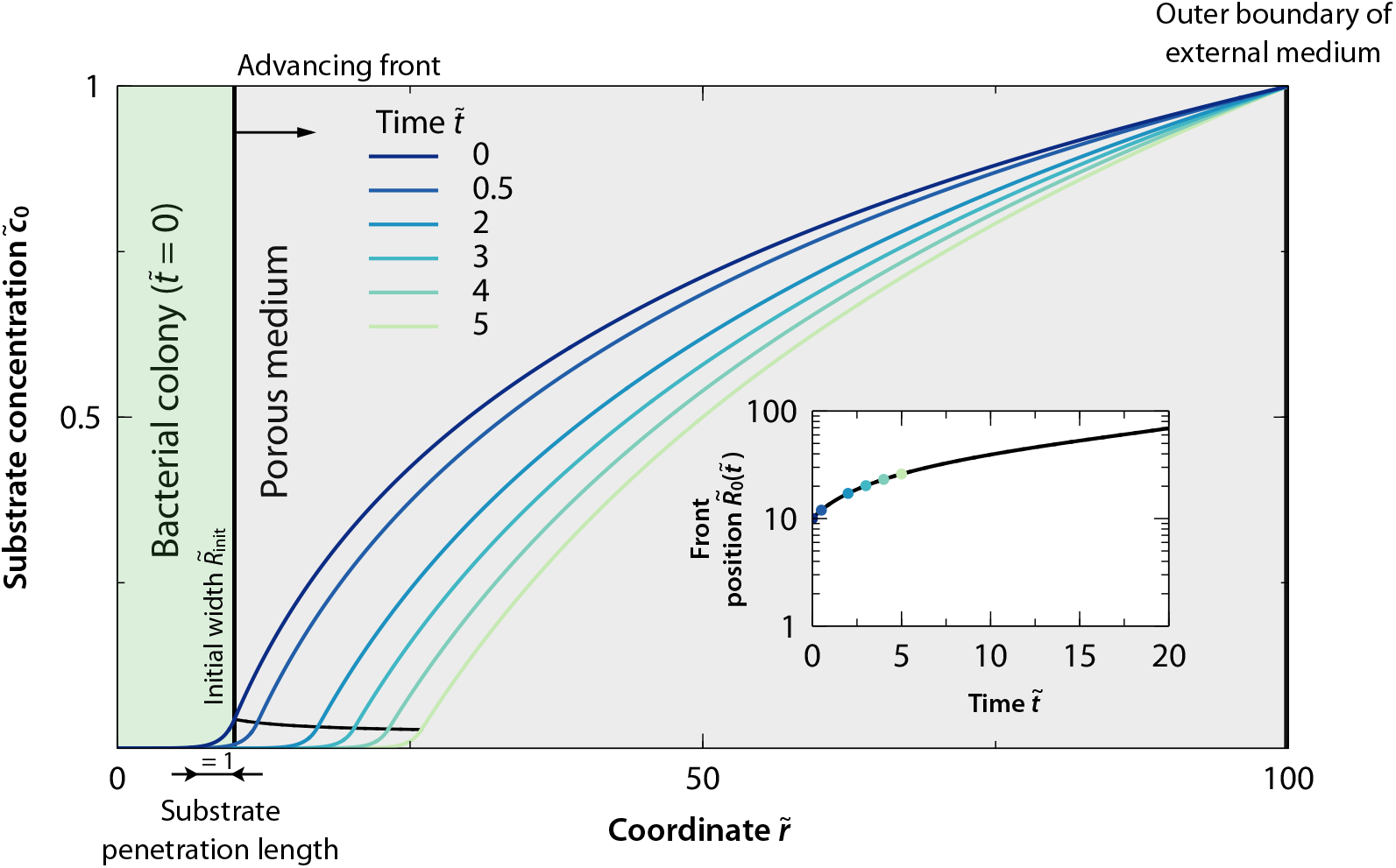
Model prediction for a flat-walled cylindrical colony growing radially outward. 1D dimensionless substrate concentration of cylindrical colony as a function of the radial coordinate 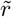 for different times, displaying both the inner and outer solutions, 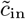 and 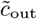, for initial dimensionless colony width 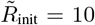 and system width 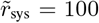, enforcing a cylindrical front. (*Inset*) Evolution of the advancing cylindrical front 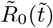 as a function of time.

**FIG. S20.**
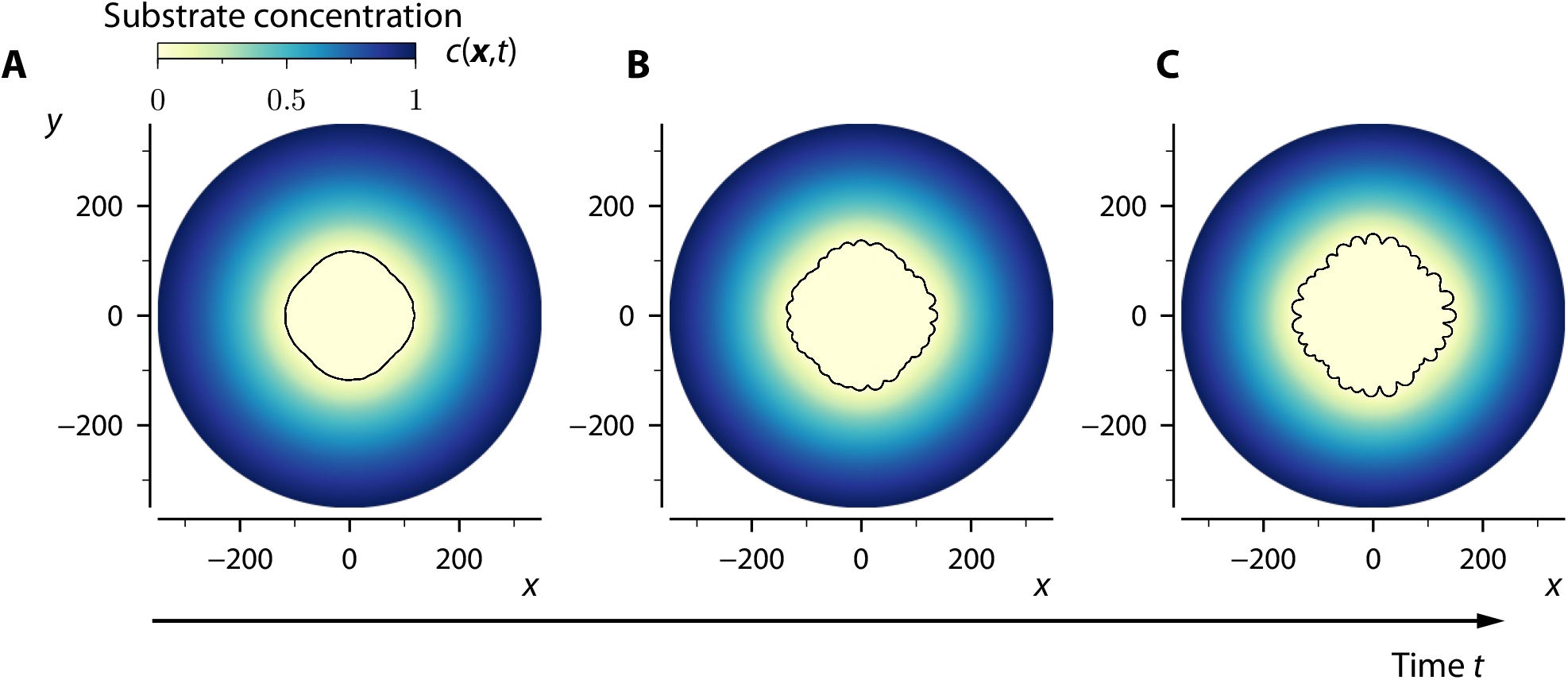
2D numerical simulations recapitulate the incipient stages of the roughening instability. Snapshots of a time-dependent 2D numerical simulation of a circular colony, Eq. (9) in the Main Text, starting with *m* = 4 and 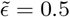 (i.e. only a single mode initially excited) and 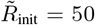, using 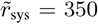 and Γ → 0, at (a) 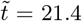, (b) 27.1, and (c) 30.2. Colors indicate the concentration of nutrient inside and outside the bacterial colony.

**FIG. S21.**
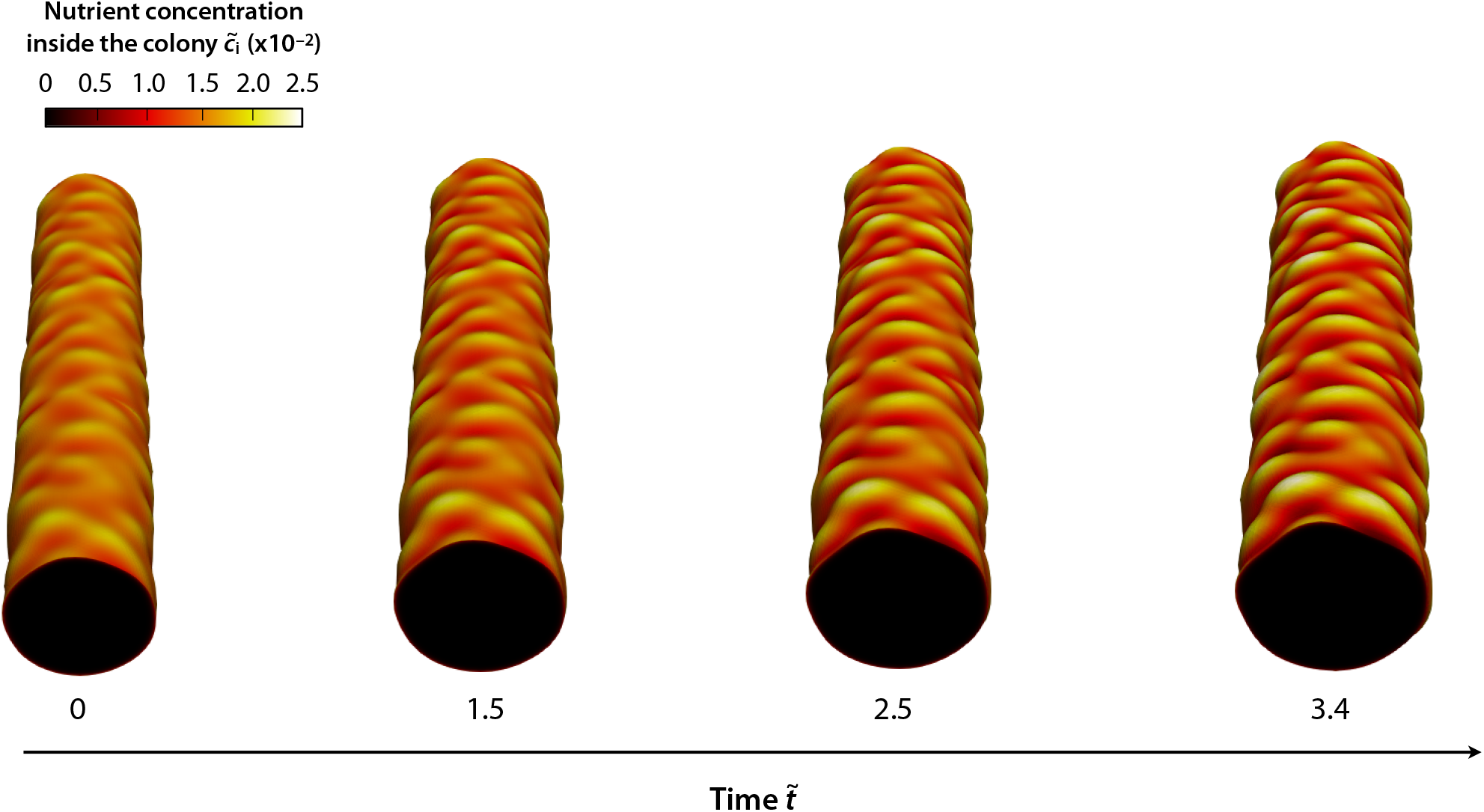
3D numerical simulations recapitulate the incipient stages of the roughening instability. Snapshots of a time-dependent 3D numerical simulation of Eq. (9) in the Main Text, starting with 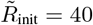, 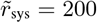, Γ = 0, at times 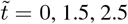, and 3.4. Colors indicate the concentration of nutrient in the interior of the bacterial colony, reveling the depletion of nutrients in the bulk. The cylindrical surface of the colony is initially perturbed with white Gaussian noise with amplitude 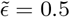.

**FIG. S22.**
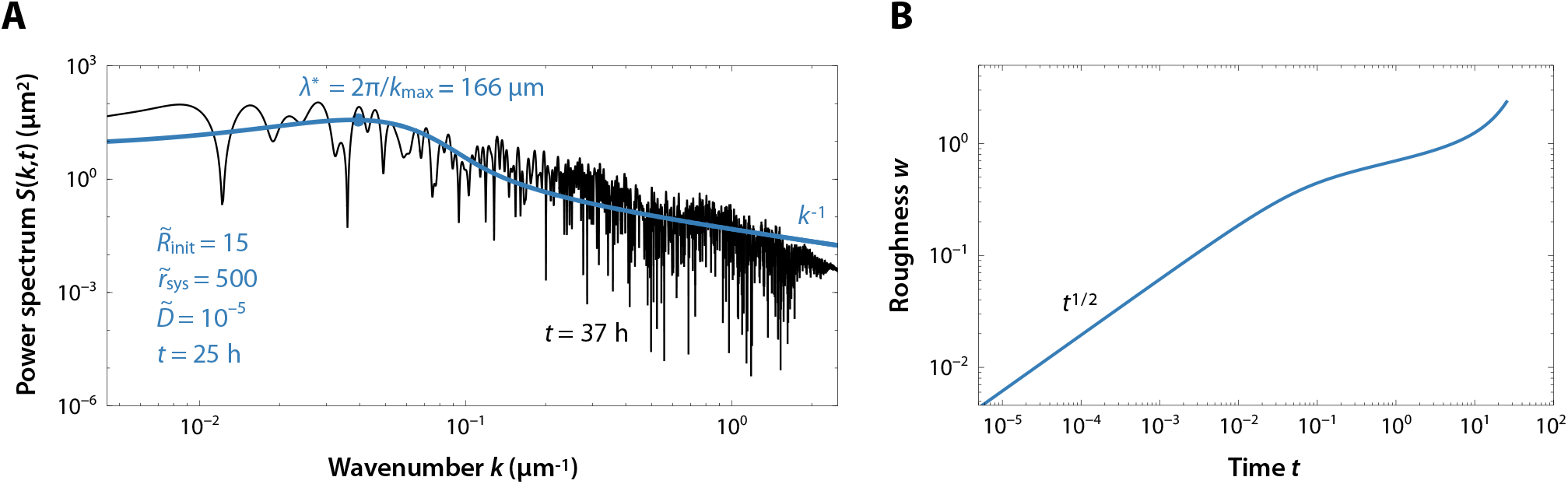
Model qualitatively recapitulates the characteristic ‘floret’ size and multi-scaled nature of colony roughening. (*A*) Experimental (black line) and theoretical (blue line) power spectra at times *t* = 37 h and *t* = 25 h, respectively. The experimental power spectrum corresponds to the *E. coli* colony displayed by Fig. 1*A* of the Main Text and Fig. S1. The values of the dimensionless parameters used for the theoretical result are 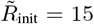, 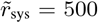, 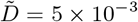, and Γ = 0. The characteristic length scale to compare both results is *l*_c_ = 8.66 μm, thus *D* = 3.7 × 10^-5^ μm^2^ s^-1^ (see estimates for *E. coli* colonies in Table 1 of the Main Text). (*B*) Theoretical global roughness *w* as a function of time *t* for the same values used in *A*. (*Inset*) The structure factor at long wavelengths, *k* ≪ 1, as a function of time.

**FIG. S23.**
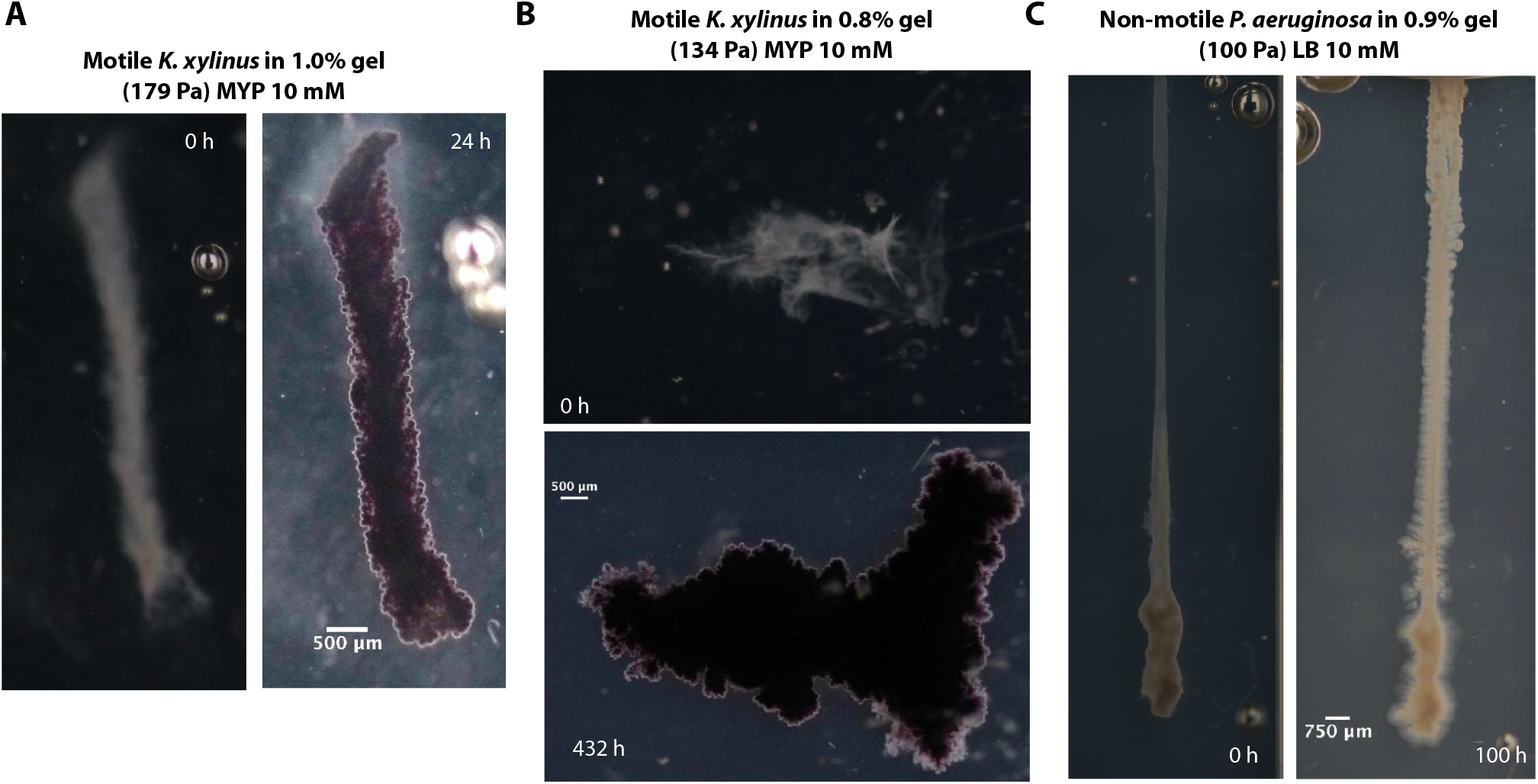
Colonies of motile *K. sucrofermentans* or non-motile *P. aeruginosa* that are immobilized in tight hydrogel matrices exhibit the same roughening instability as they grow outward in 3D. Images show snapshots taken after *t* = 24, 432, and 100 h of growth (*A–C*, respectively). The images show side-view photographs taken for colonies 3D-printed in transparent-walled containers. The comparison with Experiments #1–14 indicates that the specific cell type does not appreciably influence colony expansion and roughening.

## Notes

### Competing Interest Statement

The experimental platform used to 3D print and image bacterial communities in this publication is the subject of a patent application filed by Princeton University on behalf of T.B. and S.S.D.

## References

[1] P. Stoodley, K. Sauer, D. G. Davies, and J. W. Costerton, Annu. Rev. Microbiol. 56, 187 (2002).

[2] D. B. Kearns, Nat. Rev. Microbiol. 8, 634 (2010).

[3] A. Persat, C. D. Nadell, M. K. Kim, F. Ingremeau, A. Siryaporn, K. Drescher, N. S. Wingreen, B. L. Bassler, Z. Gitai, and H. A. Stone, Cell 161, 988 (2015).

[4] H.-C. Flemming, J. Wingender, U. Szewzyk, P. Steinberg, S. A. Rice, and S. Kjelleberg, Nat. Rev. Microbiol. 14, 563 (2016).

[5] A. Martínez-Calvo, C. Trenado-Yuste, and S. S. Datta, arXiv preprint: 2108.07011-, (2021).

[6] P. E. Kolenbrander, Annu. Rev. Microbiol. 54, 413 (2000).

[7] K. Werdan, S. Dietz, B. Löffler, S. Niemann, H. Bushnaq, R.-E. Silber, G. Peters, and U. Müller-Werdan, Nat. Rev. Cardiol. 11, 35 (2014).

[8] C. M. Dejea, P. Fathi, J. M. Craig, A. Boleij, R. Taddese, A. L. Geis, X. Wu, C. E. D. Shields, E. M. Hechenbleikner, D. L. Huso, et al., Science 359, 592 (2018).

[9] P. D. Schloss and J. Handelsman, PLoS Comput. Biol. 2, e92 (2006).

[10] R. Hayat, S. Ali, U. Amara, R. Khalid, and I. Ahmed, Ann. Microbiol. 60, 579 (2010).

[11] K. Drescher, Y. Shen, B. L. Bassler, and H. A. Stone, Proc. Nat. Acad. Sci. U.S.A. 110, 4345 (2013).

[12] R. Rusconi, M. Garren, and R. Stocker, Annu. Rev. Biophys. 43, 65 (2014).

[13] T. Liu, Y. F. Cheng, M. Sharma, and G. Voordouw, J. Pet. Sci. Eng. 156, 451 (2017).

[14] H. Fujikawa and M. Matsushita, J. Phys. Soc. Jpn. 58, 3875 (1989).

[15] E. Ben-Jacob, H. Shmueli, O. Shochet, and A. Tenenbaum, Physica A 187, 378 (1992).

[16] E. Ben-Jacob, O. Schochet, A. Tenenbaum, I. Cohen, A. Czirok, and T. Vicsek, Nature 368, 46 (1994).

[17] J. A. Shapiro, BioEssays 17, 597 (1995).

[18] D. A. Kessler and H. Levine, Nature 394, 556 (1998).

[19] M. Matsushita, J. Wakita, H. Itoh, I. Rafols, T. Matsuyama, H. Sakaguchi, and M. Mimura, Phys. A: Stat. Mech. Appl. 249, 517 (1998).

[20] E. Ben-Jacob, I. Cohen, and H. Levine, Adv. Phys. 49, 395 (2000).

[21] A. Seminara, T. E. Angelini, J. N. Wilking, H. Vlamakis, S. Ebrahim, R. Kolter, D. A. Weitz, and M. P. Brenner, Proc. Natl. Acad. Sci. U.S.A. 109, 1116 (2012).

[22] C. Fei, S. Mao, J. Yan, R. Alert, H. A. Stone, B. L. Bassler, N. S. Wingreen, and A. Košmrlj, Proc. Nat. Acad. Sci. U.S.A. 117, 7622 (2020).

[23] L. Xiong, Y. Cao, R. Cooper, W.-J. Rappel, J. Hasty, and L. Tsimring, eLife 9, e48885 (2020).

[24] J. Müller and W. Van Saarloos, Phys. Rev. E 65, 061111 (2002).

[25] R. Tokita, T. Katoh, Y. Maeda, J.-I. Wakita, M. Sano, T. Matsuyama, and M. Matsushita, J. Phys. Soc. Jpn. 78, 074005 (2009).

[26] F. Beroz, J. Yan, Y. Meir, B. Sabass, H. A. Stone, B. L. Bassler, and N. S. Wingreen, Nat. Phys. 14, 954 (2018).

[27] J. Yan, C. Fei, S. Mao, A. Moreau, N. S. Wingreen, A. Košmrlj, H. A. Stone, and B. L. Bassler, eLife 8, e43920 (2019).

[28] N. Luo, S. Wang, J. Lu, X. Ouyang, and L. You, Mol. Syst. Biol. 17, e10089 (2021).

[29] E. Ben-Jacob, D. S. Coffey, and H. Levine, Trends Microbiol. 20, 403 (2012).

[30] F. Farrell, O. Hallatschek, D. Marenduzzo, and B. Waclaw, Phys. Rev. Lett. 111, 168101 (2013).

[31] X. Wang, H. A. Stone, and R. Golestanian, New J. Phys. 19, 125007 (2017).

[32] R. J. Allen and B. Waclaw, Rep. Prog. Phys. 82, 016601 (2018).

[33] M. Mattei, L. Frunzo, B. D’acunto, Y. Pechaud, F. Pirozzi, and G. Esposito, J. Math. Biol. 76, 945 (2018).

[34] T. Bhattacharjee, D. B. Amchin, J. A. Ott, F. Kratz, and S. S. Datta, Biophys. J. 120, 3483 (2021).

[35] T. Bhattacharjee, D. B. Amchin, J. A. Alert, R. and Ott, and S. S. Datta, Elife 11, e71226 (2022).

[36] R. Alert, A. Martínez-Calvo, and S. Datta, Phys. Rev. Lett. 128, 148101 (2022).

[37] I. Klapper and J. Dockery, SIAM J. Appl. Math. 62, 853 (2002).

[38] D. Volfson, S. Cookson, J. Hasty, and L. S. Tsimring, Proc. Nat. Acad. Sci. U.S.A. 105, 15346 (2008).

[39] I. Klapper and J. Dockery, SIAM Rev. 52, 221 (2010).

[40] D. R. Espeso, A. Carpio, and B. Einarsson, Phys. Rev. E 91, 022710 (2015).

[41] C. Giverso, M. Verani, and P. Ciarletta, J. R. Soc. Interface 12, 20141290 (2015).

[42] C. Giverso and P. Ciarletta, Eur Phys J. E 39, 1 (2016).

[43] D. Dell’Arciprete, M. L. Blow, A. T. Brown, F. D. C. Farrell, J. S. Lintuvuori, A. F. McVey, D. Marenduzzo, and W. C. K. Poon, Nat. Commun. 9, 1 (2018).

[44] S. Srinivasan, C. N. Kaplan, and L. Mahadevan, eLife 8, e42697 (2019).

[45] P. Pearce, B. Song, D. J. Skinner, R. Mok, R. Hartmann, P. K. Singh, H. Jeckel, J. S. Oishi, K. Drescher, and J. Dunkel, Phys. Rev. Lett. 123, 258101 (2019).

[46] R. Hartmann, P. K. Singh, P. Pearce, R. Mok, B. Song, F. Díaz-Pascual, J. Dunkel, and K. Drescher, Nat. Phys. 15, 251 (2019).

[47] M. R. Shaebani, A. Wysocki, R. G. Winkler, G. Gompper, and H. Rieger, Nat. Rev. Phys. 2, 181 (2020).

[48] Y. Yang and H. Levine, Phys. Biol. 17, 046003 (2020).

[49] S. C. Takatori and K. K. Mandadapu, arXiv preprint arXiv:2003.05618-, (2020).

[50] Y. G. Pollack, P. Bittihn, and R. Golestanian, bioRxiv-, (2021).

[51] D. Rodriguez, B. Einarsson, and A. Carpio, Phys. Rev. E 86, 061914 (2012).

[52] G. Melaugh, J. Hutchison, K. N. Kragh, Y. Irie, A. Roberts, T. Bjarnsholt, S. P. Diggle, V. D. Gordon, and R. J. Allen, PLoS One 11, e0149683 (2016).

[53] F. D. Farrell, M. Gralka, O. Hallatschek, and B. Waclaw, J. R. Soc. Interface 14, 20170073 (2017).

[54] M. R. Warren, H. Sun, Y. Yan, J. Cremer, B. Li, and T. Hwa, eLife 8, e41093 (2019).

[55] M. J. I. Müller, B. I. Neugeboren, D. R. Nelson, and A. W. Murray, Proc. Nat. Acad. Sci. U.S.A. 111, 1037 (2014).

[56] C. D. Nadell, K. Drescher, and K. R. Foster, Nat. Rev. Microbiol. 14, 589 (2016).

[57] W. Liu, T. A. Tokuyasu, X. Fu, and C. Liu, Curr. Opin. Microbiol. 63, 109 (2021).

[58] M. Whiteley, M. G. Bangera, R. E. Bumgarner, M. R. Parsek, G. M. Teitzel, S. Lory, and E. P. Greenberg, Nature 413, 860 (2001).

[59] T.-F. Mah, B. Pitts, B. Pellock, G. C. Walker, P. S. Stewart, and G. A. O’Toole, Nature 426, 306 (2003).

[60] D. Nguyen, A. Joshi-Datar, F. Lepine, E. Bauerle, O. Olakanmi, K. Beer, G. McKay, R. Siehnel, J. Schafhauser, Y. Wang, et al., Science 334, 982 (2011).

[61] W. O. H. Hughes and J. J. Boomsma, Evolution 58, 1251 (2004).

[62] A. R. Hughes, B. D. Inouye, M. T. J. Johnson, N. Underwood, and M. Vellend, Ecol. Lett. 11, 609 (2008).

[63] T. B. H. Reusch, A. Ehlers, A. Hämmerli, and B. Worm, Proc. Nat. Acad. Sci. U.S.A. 102, 2826 (2005).

[64] O. Hallatschek, P. Hersen, S. Ramanathan, and D. R. Nelson, Proc. Nat. Acad. Sci. U.S.A. 104, 19926 (2007).

[65] O. Hallatschek and D. R. Nelson, Theoretical population biology 73, 158 (2008).

[66] O. Hallatschek and D. R. Nelson, Evolution: International Journal of Organic Evolution 64, 193 (2010).

[67] K. S. Korolev, M. Avlund, O. Hallatschek, and D. R. Nelson, Rev. Mod. Phys. 82, 1691 (2010).

[68] J. Kayser, C. F. Schreck, M. Gralka, D. Fusco, and O. Hallatschek, Nature ecology & evolution 3, 125 (2019).

[69] S. Chu, M. Kardar, D. R. Nelson, and D. A. Beller, J. Theor. Biol. 478, 153 (2019).

[70] G. T. Fortune, N. M. Oliveira, and R. E. Goldstein, Phys. Rev. Lett. 128, 178102 (2022).

[71] R. Rusconi, S. Lecuyer, L. Guglielmini, and H. A. Stone, J. R. Soc. Interface 7, 1293 (2010).

[72] Q. Zhang, J. Li, J. Nijjer, H. Lu, M. Kothari, R. Alert, T. Cohen, and J. Yan, Proc. Nat. Acad. Sci. U.S.A. 118 (2021).

[73] G. A. Stooke-Vaughan and O. Campàs, Curr. Opin. Genet. Dev. 51, 111 (2018).

[74] K. Goodwin and C. M. Nelson, Dev. cell 56, 240 (2021).

[75] D. Wirtz, K. Konstantopoulos, and P. C. Searson, Nat. Rev. Cancer 11, 512 (2011).

[76] A. M. J. Valencia, P.-H. Wu, O. N. Yogurtcu, P. Rao, J. DiGiacomo, I. Godet, L. He, M.-H. Lee, D. Gilkes, S. X. Sun, et al., Oncotarget 6, 43438 (2015).

[77] C. D. Paul, W.-C. Hung, D. Wirtz, and K. Konstantopoulos, Annu. Rev. Biomed. Eng. 18, 159 (2016).

[78] H. Ahmadzadeh, M. R. Webster, R. Behera, A. M. J. Valencia, D. Wirtz, A. T. Weeraratna, and V. B. Shenoy, Proc. Natl. Acad. Sci. U. S. A. 114, E1617 (2017).

[79] P.-H. Wu, D. M. Gilkes, and D. Wirtz, Annu. Rev. Biophys. 47, 549 (2018).

[80] J. N. Wilking, T. E. Angelini, A. Seminara, M. P. Brenner, and D. A. Weitz, MRS Bull. 36, 385 (2011).

[81] M. Schaffner, P. A. Rühs, F. Coulter, S. Kilcher, and A. R. Studart, Sci. Adv. 3, eaao6804 (2017).

[82] C. Douarche, A. Buguin, H. Salman, and A. Libchaber, Phys. Rev. Lett. 102, 198101 (2009).

[83] X. Fu, S. Kato, J. Long, H. H. Mattingly, C. He, D. C. Vural, S. W. Zucker, and T. Emonet, Nat. Commun. 9, 1 (2018).

[84] O. A. Croze, G. P. Ferguson, M. E. Cates, and W. Poon, Biophys. J. 101, 525 (2011).

[85] T. Bhattacharjee, S. M. Zehnder, K. G. Rowe, S. Jain, R. M. Nixon, W. G. Sawyer, and T. E. Angelini, Sci. Adv. 1, e1500655 (2015).

[86] T. Bhattacharjee, C. P. Kabb, C. S. O’Bryan, J. M. Uruena, B. S. Sumerlin, W. G. Sawyer, and T. E. Angelini, Soft Matter 14, 1559 (2018).

[87] T. Bhattacharjee and S. Datta, Nat. Commun. 10, 1 (2019).

[88] T. Bhattacharjee and S. Datta, Soft Matter 15, 9920 (2019).

[89] A.-L. Barabási and H. E. Stanley, Fractal concepts in surface growth (Cambridge university press, 1995).

[90] M. Kardar, G. Parisi, and Y.-C. Zhang, Phys. Rev. Lett. 56, 889 (1986).

[91] J. A. Bonachela, C. D. Nadell, J. B. Xavier, and S. A. Levin, J. Stat. Phys. 144, 303 (2011).

[92] C. D. Nadell, V. Bucci, K. Drescher, S. A. Levin, B. L. Bassler, and J. B. Xavier, Proc. R. Soc. B: Biol. Sci. 280, 20122770 (2013).

[93] E. Young, G. Melaugh, and A. R. J., bioRxiv-, (2022).

[94] C. Blanch-Mercader and J. Casademunt, Phys. Rev. Lett. 110, 078102 (2013).

[95] M. J. Bogdan and T. Savin, R. Soc. Open Sci. 5, 181579 (2018).

[96] A. Mongera, P. Rowghanian, H. J. Gustafson, E. Shelton, D. A. Kealhofer, E. K. Carn, F. Serwane, A. A. Lucio, J. Giammona, and O. Campàs, Nature 561, 401 (2018).

[97] S. P. Banavar, E. K. Carn, P. Rowghanian, G. Stooke-Vaughan, S. Kim, and O. Campàs, Sci. Rep. 11, 1 (2021).

[98] Y. Guo, M. Nitzan, and M. P. Brenner, PLoS Comput. Biol. 17, e1009576 (2021).

[99] J. Li, S. K. Schnyder, M. S. Turner, and R. Yamamoto, Phys. R. X 11, 031025 (2021).

[100] M. Martin and T. Risler, New J. Phys. 23, 033032 (2021).

[101] L. Michaelis, M. L. Menten, et al., Biochem. Z. 49, 352 (1913).

[102] K. A. Johnson and R. S. Goody, Biochemistry 50, 8264 (2011).

[103] T. Vicsek, M. Cserző, and V. K. Horváth, Phys. A: Stat. Mech. Appl. 167, 315 (1990).

[104] S. N. Santalla and S. C. Ferreira, Phys. Rev. E 98, 022405 (2018).

[105] S. N. Santalla, J. Rodríguez-Laguna, J. P. Abad, I. Marín, M. del Mar Espinosa, J. Muñoz-García, L. Vázquez, and R. Cuerno, Phys. Rev. E 98, 012407 (2018).

[106] T. R. Thomas, Rough surfaces (London, London: Logman).

[107] F. Family and T. Vicsek, J. Phys. A Math. Theor. 18, L75 (1985).

[108] F. Family, J. Phys. A Math. Theor. 19, L441 (1986).

[109] M. Rubio, C. Edwards, A. Dougherty, and J. Gollub, Phys. Rev. Lett. 63, 1685 (1989).

[110] V. K. Horváth, F. Family, and T. Vicsek, J. Phys. A Math. Theor. 24, L25 (1991).

[111] S. Buldyrev, A.-L. Barabási, F. Caserta, S. Havlin, H. Stanley, and T. Vicsek, Phys. Rev. A 45, R8313 (1992).

[112] S. He, G. L. Kahanda, and P.-Z. Wong, Phys. Rev. Lett. 69, 3731 (1992).

[113] J. Zhang, Y.-C. Zhang, P. Alstrøm, and M. Levinsen, Phys. A: Stat. Mech. Appl. 189, 383 (1992).

[114] T. Halpin-Healy and Y.-C. Zhang, Phys. Rep. 254, 215 (1995).

[115] R. Cuerno and A.-L. Barabási, Phys. Rev. Lett. 74, 4746 (1995).

[116] J. Krug, Adv. Phys. 46, 139 (1997).

[117] J. Soriano, J. J. Ramasco, M. A. Rodríguez, A. Hernández-Machado, and J. Ortín, Phys. Rev. Lett. 89, 026102 (2002).

[118] R. M. Sutherland, Science 240, 177 (1988).

[119] J. Soriano, A. Mercier, R. Planet, A. Hernández-Machado, M. Rodríguez, and J. Ortín, Phys. Rev. Lett. 95, 104501 (2005).

[120] A. Birovljev, L. Furuberg, J. Feder, T. Jssang, K. Mly, and A. Aharony, Phys. Rev. Lett. 67, 584 (1991).

[121] L.-H. Tang and H. Leschhorn, Phys. Rev. A 45, R8309 (1992).

[122] L.-H. Tang, M. Kardar, and D. Dhar, Phys. Rev. Lett. 74, 920 (1995).

[123] G. Parisi, EPL 17, 673 (1992).

[124] J. M. López, M. A. Rodriguez, and R. Cuerno, Phys. Rev. E 56, 3993 (1997).

[125] J. M. López, Phys. Rev. Lett. 83, 4594 (1999).

[126] J. J. Ramasco, J. M. López, and M. A. Rodríguez, Phys. Rev. Lett. 84, 2199 (2000).

[127] U. Böckelmann, W. Manz, T. R. Neu, and U. Szewzyk, FEMS Microbiol. Ecol. 33, 157 (2000).

[128] K. Busch, S. Endres, M. H. Iversen, J. Michels, E.-M. Nöthig, and A. Engel, Front. Mar. Sci 4, 166 (2017).

[129] B. Zäncker, A. Engel, and M. Cunliffe, J. Plankton Res. 41, 561 (2019).

[130] D. A. Rosen, T. M. Hooton, W. E. Stamm, P. A. Humphrey, and S. J. Hultgren, PLoS Med. 4, e329 (2007).

[131] E. Szabó, R. Liébana, M. Hermansson, O. Modin, F. Persson, and B.-M. Wilén, Front. Microbiol. 8, 770 (2017).

[132] B.-M. Wilén, B. Jin, and P. Lant, Water Res. 37, 3632 (2003).

[133] B. Bottura, L. M. Rooney, P. A. Hoskisson, and G. McConnell, bioRxiv-, (2021).

[134] A. Prindle, J. Liu, M. Asally, S. Ly, J. Garcia-Ojalvo, and G. M. Süel, Nature 527, 59 (2015).

[135] J. Liu, A. Prindle, J. Humphries, M. Gabalda-Sagarra, M. Asally, D.-Y. D. Lee, S. Ly, J. Garcia-Ojalvo, and G. M. Süel, Nature 523, 550 (2015).

[136] J. Humphries, L. Xiong, J. Liu, A. Prindle, F. Yuan, H. A. Arjes, L. Tsimring, and G. M. Süel, Cell 168, 200 (2017).

[137] J. Liu, R. Martinez-Corral, A. Prindle, D.-Y. D. Lee, J. Larkin, M. Gabalda-Sagarra, J. Garcia-Ojalvo, and G. M. Süel, Science 356, 638 (2017).

[138] J. W. Larkin, X. Zhai, K. Kikuchi, S. E. Redford, A. Prindle, J. Liu, S. Greenfield, A. M. Walczak, J. Garcia-Ojalvo, A. Mugler, et al., Cell Sys. 7, 137 (2018).

[139] R. Martinez-Corral, J. Liu, G. M. Süel, and J. Garcia-Ojalvo, Proc. Nat. Acad. Sci. U.S.A. 115, E8333 (2018).

[140] J. A. Schwartzman, A. Ebrahimi, G. Chadwick, Y. Sato, V. Orphan, and O. X. Cordero, bioRxiv-, (2021).

[141] K. Tong, G. O. Bozdag, and W. C. Ratcliff, Curr. Opin. Microbiol. 67, 102141 (2022).

[142] A. Bonforti and R. Sole, Preprints-, (2021).

[143] W. C. Ratcliff, R. F. Denison, M. Borrello, and M. Travisano, Proc. Natl. Acad. Sci. U.S.A. 109, 1595 (2012).

[144] S. Jacobeen, J. T. Pentz, E. C. Graba, C. G. Brandys, W. C. Ratcliff, and P. J. Yunker, Nat. Phys. 14, 286 (2018).

[145] S. Jacobeen, E. C. Graba, C. G. Brandys, T. C. Day, W. C. Ratcliff, and P. J. Yunker, Phys. Rev. E 97, 050401 (2018).

[146] S. A. Zamani-Dahaj, A. Burnetti, T. C. Day, P. J. Yunker, W. C. Ratcliff, and M. D. Herron, bioRxiv-, (2021).

[147] T. C. Day, S. S. Höhn, S. A. Zamani-Dahaj, D. Yanni, A. Burnetti, J. Pentz, A. R. Honerkamp-Smith, H. Wioland, H. R. Sleath, W. C. Ratcliff, et al., eLife 11, e72707 (2022).

[148] J. Yan, A. G. Sharo, H. A. Stone, N. S. Wingreen, and B. L. Bassler, Proceedings of the National Academy of Sciences 113, E5337 (2016).

[149] T. Bhattacharjee, C. J. Gil, S. L. Marshall, J. M. Urueña, C. S. O’Bryan, M. Carstens, B. Keselowsky, G. D. Palmer, S. Ghivizzani, C. P. Gibbs, et al., ACS Biomater. Sci. Eng. 2, 1787 (2016).

